# Magnetic printing and actuation of stretchable muscle tissue

**DOI:** 10.1101/2025.02.19.638972

**Authors:** Noam Demri, Lise Morizur, Simon Dumas, Giacomo Gropplero, Cécile Martinat, Stéphanie Descroix, Claire Wilhelm

**Affiliations:** Laboratoire Physico Chimie Curie, PCC, CNRS UMR168, Institut Curie, Sorbonne University, PSL University, 75005 Paris, France; INSERM/UEVE UMR 861, Université Paris Saclay, I-STEM, Corbeil-Essonnes, France

## Abstract

While the link between tissue organization, stimulation, and function is now acknowledged as crucial for tissue development, engineering tissues with precise, long-lasting shapes and the capability for mechanical stimulation remains challenging. This study addresses this challenge by developing a next-generation magnetic bioprinting approach to create anisotropic, shape-controlled, scaffold-free, and stretchable skeletal muscle constructs.

Murine skeletal muscle cells and human induced pluripotent stem cell-derived skeletal muscle cells, labeled with iron oxide nanoparticles, were magnetically bioprinted into wrench-shaped tissues. Their magnetic properties allowed these tissues to be clipped onto magnetic needles, preserving their shape over two weeks of culture while promoting anisotropic differentiation and myoblast fusion. Additionally, the magnetic tissues could be stretched by up to 100%, enhancing their anisotropy and improving muscle maturation.

This magnetic toolbox demonstrates significant advancements in muscle tissue engineering, as evidenced by enhanced indicators of myoblast differentiation, including cell fusion, increased myogenic maturation and contractility. These findings highlight the potential of magnetic-based techniques for developing advanced muscle-on-chip systems and other complex tissue constructs.

## Introduction

Due to the inability of monolayer cell cultures to accurately replicate the complexity of tissues, there is an increasing need for systems that can fully capture the *in vivo* tissue organization^1,2^. This challenge is currently being addressed through the development of organ-on-chip and bioprinting technologies. Microfluidic systems, used to create organs-on-a-chip^3–9^, enable culturing cells in chambers that mimic the *in vivo* microenvironment, typically within a three-dimensional (3D) scaffold, to achieve a high level of control in recapitulating certain organ functions, in drug screening and to dissect tissue biology. However, current organ-on-a-chip approaches still lack control over the overall tissue shape, which remains a critical challenge in tissue engineering^10–13^. In contrast, bioprinting addresses the essential link between tissue shape and function^14,15^ by allowing precise control over tissue geometrical cues. Bioprinting typically involves the use of bioink filled with cells, which is either extruded^16,17^ or photopolymerized^18^ in the desired shape. Despite their advantages, bioinks can be physiologically limiting and restrict cell density and cell-to-cell interactions. Furthermore, while bioprinting methods can accurately control the initial shape of bioprinted tissues, they do not guarantee the maintenance of this geometry throughout tissue maturation.

And yet, many tissues, such as skeletal muscle tissue^19^, require both strong geometrical cues^20^ and a densely packed cellular network over an extended period for proper tissue maturation^21^. Myogenesis occurs when a dense network of myoblasts, mononucleated muscle precursor cells, fuse together to form parallel giant multinucleated myofibers. This geometry is crucial for optimal maturation and functional force generation *in vivo.* The increasing contractility of differentiating cells also induces significant stresses that can deform engineered *in vitro* tissues, thereby limiting their maturation and, consequently, their physiological relevance.

Preserving the original anisotropic geometry of the tissue is therefore essential for skeletal muscle constructs. Aside from 2D systems on anisotropic substrates^22^, most muscle-on-chip models constrain the tissue by anchoring it between two pillars^23–27^. Such approach favors the maintenance of 3D anisotropy and can facilitate contractibility measurements through post deflection ^25,26,28^. However, these systems generally require the use of a scaffold to maintain anisotropy. Additionally, achieving consistent reproducibility and precise shape control over time remains challenging.

The advent of magnetic-based tissue engineering techniques has recently introduced new possibilities for creating complex, scaffold-free tissues. Magnetic forces can effectively direct the organization of cells that have internalized magnetic nanoparticles^29,30^. Previously used for theragnostic purposes - such as imaging contrast agents, drug delivery vectors and anticancer hyperthermal therapies^31,32^, these nanoparticles have also shown promise in remotely controlling cells within a tissue engineering context. Initiated by magnetic sheet engineering^33^, magnetic bioprinting approaches have successfully created matrix-free tissues in specific shapes ^29,34–36^. However, these magnetic tissues have not yet achieved the complexity seen in bioprinting. Nevertheless, magnetic remote actuation has proven remarkably effective for 3D spheroid assembly^29,37^, mechanical stimulation ^38^, and trapping cell aggregates^36,39,40^.

Herein, we present a next generation magnetic bioprinting approach for skeletal muscle constructs, engineered to be anisotropic, shape-controlled, scaffold-free, and with inherent capacity for physical stimulation. Indeed, functionalizing the shape of the tissues enabled magnetic actuation, allowing them to be remotely clipped onto two magnetic needles while preserving their shape overtime, and enabling controlled stretching. Once optimized with C2C12 mouse myoblasts, this “clip-on” two-fold magnetic process was successfully applied to myoblasts derived from human induced pluripotent stem cells (M-hiPSC). In both cases, stretching the tissues improved their maturation and differentiation, resulting in the rapid formation of fascicle-like skeletal muscle tissue.

## Results

### Magnetic bioprinting of wrench-shaped ’clip-on’ muscle tissues

C2C12 mouse myoblasts were magnetically labeled by incubating them sequentially with maghemite nanoparticles. As demonstrated in a previous study^29^, this process led to the internalization of approximatively 20pg of iron per cell through endocytosis^41^, without affecting the cells viability.

The maghemite nanoparticles are superparamagnetic: upon internalization, they enable the remote control of the cells using magnetic forces without any residual magnetism once the magnetic field is removed. Our approach aims to create a magnetic muscle tissue with controllable anisotropy and shape that can be further manipulated by magnetic forces for stretching stimulation. We envisioned a wrench-shaped form (Fig. 1A) with the ends designed as two clamps to clip onto magnetic attractors, such as needles, which could then be moved to enable stretching. To create the shape, we conceived a magnetic patterning strategy that could potentially replace the printing pattern during bioprinting. We employed photolithography and electrodeposition to create soft NiFe (permalloy) micropatterns in any desired shape (Supplementary Fig. 1A-B). The selected wrench-shaped pattern (Fig. 1A) features a 500 µm wide and 2 mm long central fiber that mimics the anisotropic muscle structure and two clamps for functional purposes, while remaining large enough for effective manipulation. Once the pattern is positioned between two strong magnets (Supplementary Fig. 1C), where the magnetic field is uniform, it magnetizes, generating a high magnetic gradient around the pattern, as demonstrated by finite element simulations (Fig 1B, Supplementary Fig. 1D-L). By placing a suspension of magnetically labeled cells in a well, positioned on top of the patterns, cells are exposed to high magnetic forces at the bottom of the well. Simulations indicate that the magnetic gradient there varies between 40 and 800 T.m^-1^, resulting in forces ranging from 0.05 to 1.5 nN on a single cell (with a magnetic moment at saturation of 1.7 x 10^-12^ A.m^2^). These forces caused the cells to migrate and align with the shape of the patterns within seconds (Supplementary movie 1). The cells were maintained under the magnetic field for three hours to ensure cohesiveness. As a result, the tissues could be easily collected once the patterns were removed from the magnetic field (Fig 1C). Thus, this magnetic bioprinting technique results in the rapid formation of cohesive tissues in the exact shape of the magnetic patterns (Fig. 1D). Unlike most other bioprinting methods, which typically rely on a gel or matrix, the resulting tissue is composed solely of cells. Importantly, live dead assays on freshly patterned tissues (Fig. 1E, Supplementary Fig. 2) demonstrated that this magnetic bioprinting process does not induce cell death, with 94±2% of live cells after tissue formation. Interestingly, while cells follow the 2D pattern, the resulting tissue is three-dimensional (3D) (Fig. 1F-G, Supplementary Fig. 3), with cells aggregating in a densely packed manner on top of one another, resulting in a 3D type of bioprinting. Varying the cell density from 10^5^ to 4×10^5^ cells per mm^2^ of pattern resulted in an increase in the width and height of the central fiber, which on average grew from 388 µm and 300 µm to 692 µm and 547 µm, respectively (Fig. 1H and 1I). Cell densities can therefore be used to control the thickness of the tissue, while the ratio between width and height remains constant (Fig. 1J). In the following experiments, tissues were created with a cell density of 10^5^ cells per mm^2^ of pattern, as this density minimizes the number of required cells and results in tissues that closely match the size of the pattern without compromising oxygen and nutrient diffusion within the tissue^42^. The magnetic properties of the bioprinted tissues were monitored over time, as they are central to the entire magnetic bioprinting process. Additionally, for potential future clinical applications, it is important to eventually eliminate the presence of nanoparticles within the tissues through cellular assimilation involving iron metabolism^41^. To investigate whether such assimilation could take place in the bioengineered magnetic muscular tissues, bioprinted tissues were cultured in non-adherent 500µL tubes and their magnetic signals were regularly measured using a benchtop magnetometer^43^. Consistent with previous studies indicating that cells degrade magnetic nanoparticles over time^41,43^, the samples exhibited a linear decrease in the magnetic signal with a total loss of about 21% from day 1 to day 20 (Fig. 1K).

**Figure 1.**
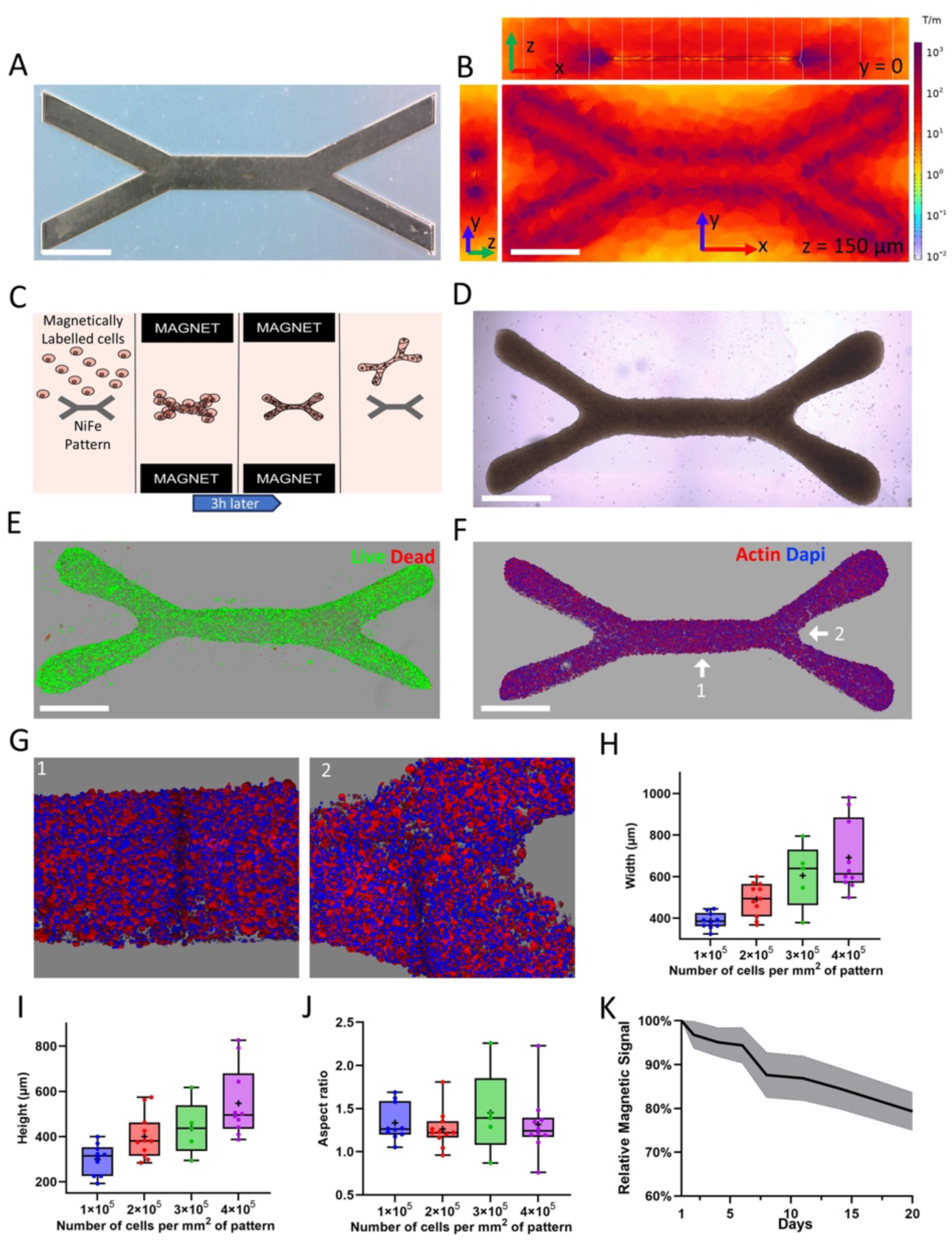
Magnetic micropatterns generate viable, shape-controlled 3D tissues. **A** Wrench-shaped magnetic micropattern. **B** Magnetic field gradient generated when the pattern is placed between two strong magnets. **C** Patterning procedure. **D** Typical bright-field image of a magnetically bioprinted tissue. **E** 3D reconstitution from confocal imaging of a live/dead assay on a tissue immediately after patterning (live cells in green, dead cells in red). **F**-**G** 3D reconstitutions from confocal microscopy of a tissue (actin in red, nuclei in blue): whole tissue (F), zoomed-in views of zones 1 and 2 (G). **H-J** Boxplots of the width (H), height (I) and aspect ratio (J) of the section of the central fiber of the tissue after patterning (whiskers as min and max, box as the interquartile range, center line as median and mean as +). **K** Variation of the magnetic signal of bioprinted tissues with time (n=9) plotted relative to the signal at day 1 until day 20. Scale bars = 1 mm.

### Trapping the magnetic-printed muscle tissue between two magnetic needles

Once removed from the pattern, each tissue is transferred into one of the eight microfabricated chambers of a chip filled with differentiation medium and made of a soft silicone – Ecoflex material (Fig 2A, Supplementary Fig. 4), which will allow stretching later on. Each compartment contains two steel needles facing one another at a 2 mm distance, magnetized by a pair of adjacent magnets attracting each other. The magnetic field gradient (Fig. 2B) is focused on the tips of the needles: when a tissue is pipette-deposited in the center of a chamber containing the two facing needles, it immediately attaches to the needles with its two clamps (Fig. 2C, supplementary movie 2), transforming the muscle construct into a clip-on tissue. Remarkably, the clip-on process is both efficient and reproducible, with a nearly 100% success rate (Fig. 2D).

**Figure 2.**
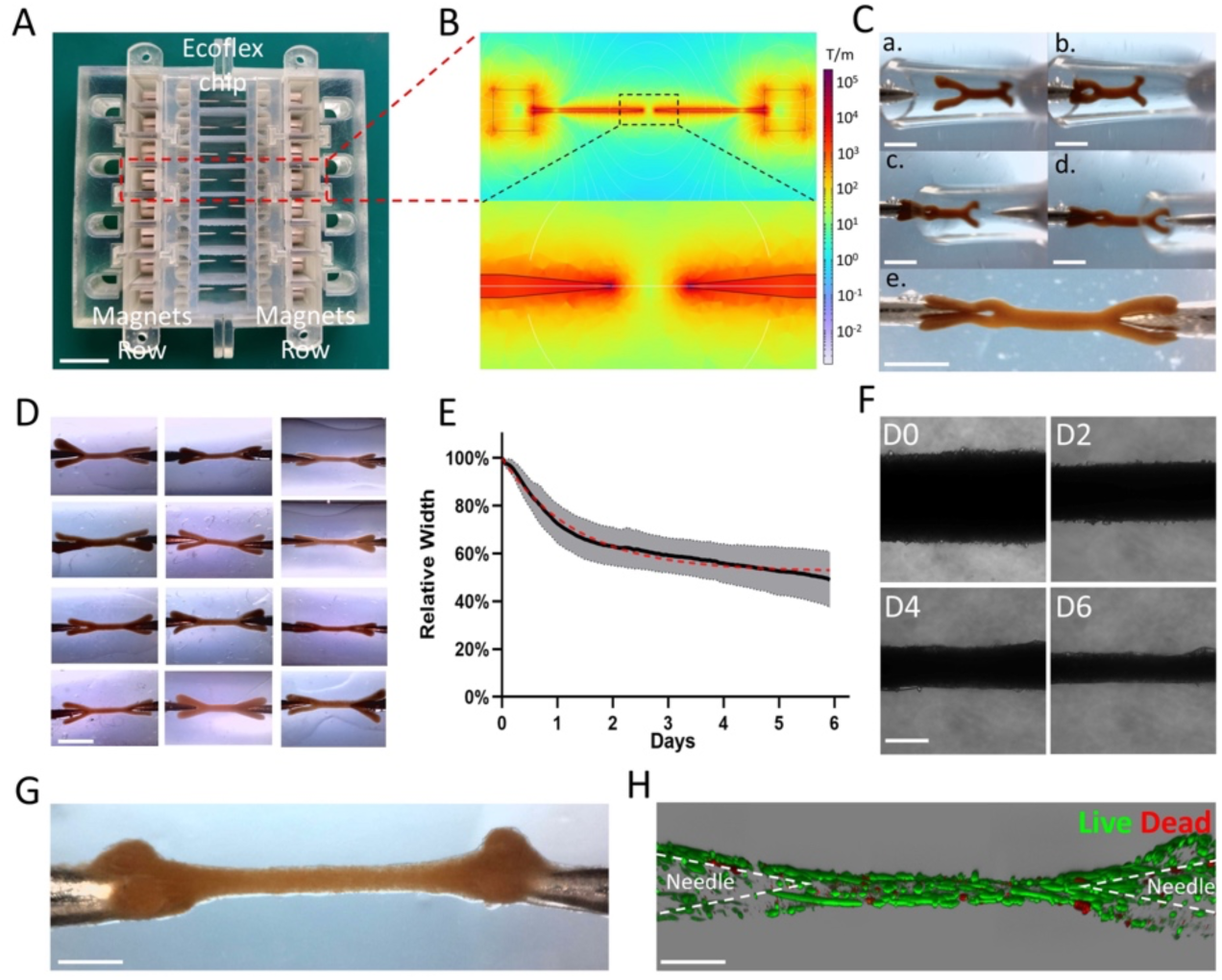
Tissue shape is preserved by magnetic trapping. **A** Ecoflex chip (schematics in supplementary Fig. 4B) with 8 compartments of magnetized needles and the adjacent magnet holders (schematics in supplementary Fig. 4C). Scale bar = 2 cm. **B** Magnetic field gradient around the magnets and needles, with a close-up between two needles. **C** Trapping procedure to clip a tissue on two needles. Scale bar = 1 mm. **D** Tissues made from C2C12 cells clipped on the needles. Scale bar = 2 mm. **E-F** Evolution of the central fiber’s width (E) plotted relative to the width at day 0 over 6 days (n=4) and (F) illustrated with bright-field microscopy. Scale bar = 200 µm. **G-H** Fiber made from C2C12 imaged after 6 days of differentiation (G) from the top with a binocular magnifier and (H) with confocal imagery after a live/dead assay (live cells in green, dead cells in red). Scale bar = 500 µm.

After the tissues are magnetically trapped, they are left to adhere overnight to the needles, with the permanent magnets in place to maintain the attraction forces. It is important to note that these forces are present only near the tips of the needles while the magnetic gradient is close to 0 between the needles. For reference, at a distance of 100 µm from the tips, simulations indicate that a single magnetized cell experiences an 8000 T.m^-1^ gradient (Fig. 2B, Supplementary Fig. 5), resulting in an attractive force of approximately 15 nN on the cell. The next day, the magnets are removed, releasing the initial magnetic forces that facilitated tissue placement, initial attachment, and cohesiveness of the tissue clamps. Since the needles are coated with polydopamine and collagen, tissues remain attached to their respective anchor needles.

As soon as muscle differentiation is initiated on day 0 by replacing the medium with differentiation medium, the central fibers of the trapped tissues exhibit significant thinning over the following days (Fig. 2E-F, Supplementary Fig. 6). This thinning initially occurs at an approximately linear rate before gradually plateauing. This evolution can be described by a decreasing exponential curve, characterized by a time constant of approximately 30 hours. Tissues are maintained between the needles for a maximum of 14 days. Thanks to the magnetic trapping and subsequent cellular adhesion, the overall tissue anisotropy is systematically preserved (Fig. 2G, Supplementary Fig. 7). In contrast, similar tissues cultured in a non-adherent well, free from constrains, quickly lose their shape and eventually become more spherical (Supplementary Fig. 7D) to minimize surface tension. Importantly, tissues cultured on the needles for 6 days exhibited almost no cell death with 94±2% of live cells, even at the needle sites, demonstrating the long-term viability of the trapping procedure (Fig. 2H, Supplementary Fig. 8).

### Preserving tissue anisotropy for myogenesis

The muscle differentiation is initially assessed by examining internal C2C12 cell reorganization. Figure 3A and Supplementary Figure 9 show that on day 0 cells are round, mononucleated and organized isotropically. Over time, they progressively fuse together and become increasingly elongated and aligned in the direction of the overall tissue (Fig. 3B-3C, Supplementary Fig. 10A-10C, 11A-11B and 12A-12C). By day 6, the central fiber of the tissue resembles a fascicle, with giant fused myotubes all aligned in the same direction (Fig. 3D-3E, Supplementary Fig. 13A-13D). Regarding cell fusion, from the original population of myoblasts at day 0, only 36% remains mononucleated at day 6, while the remaining 64% fused to form multinucleated myotubes with at least two nuclei (Fig. 3F). Notably, 28 % of the original myoblasts developed into multinucleated cells with more than 10 nuclei, some of which contained over 60 nuclei and approached millimeter-scale sizes (Supplementary Fig. 14A). At this stage, 53% of the actin filaments are oriented within ±15° of the fiber’s direction, compared to only 18% on day 0. C2C12 cells cultured in 2D showed no overall anisotropy at day 6, and no matter the direction chosen, the percentage of actin filaments within a ±15° range did not exceed 21% at best (Fig. 3G, Supplementary Fig. 15A-15C). Intriguingly, while the 3D tissues become increasingly anisotropic over time (Fig. 3G), some samples on day 6 show a slight shift in the direction of cell alignment. 3D reconstruction reveals that these cells appear to be twisting along the tissue axis, possibly to better distribute forces within the tissue. This shift appears to slightly persist in tissues cultured up to 14 days. However, overall, from day 4 to day 14, the distribution of actin filaments alignment remains mostly constant, with 47% of the filaments still aligned within ±15° of the fiber’s direction on day 14. One noticeable change observed in samples cultured for two weeks (Fig. 3H) is the width of the multinucleated myotubes. The average width of myotubes with more than 10 nuclei at day 6 is 35 µm, compared to 11µm for mononucleated myoblasts at day 0, and further increases to 42 µm after 2 weeks (Fig. 3I).

**Figure 3.**
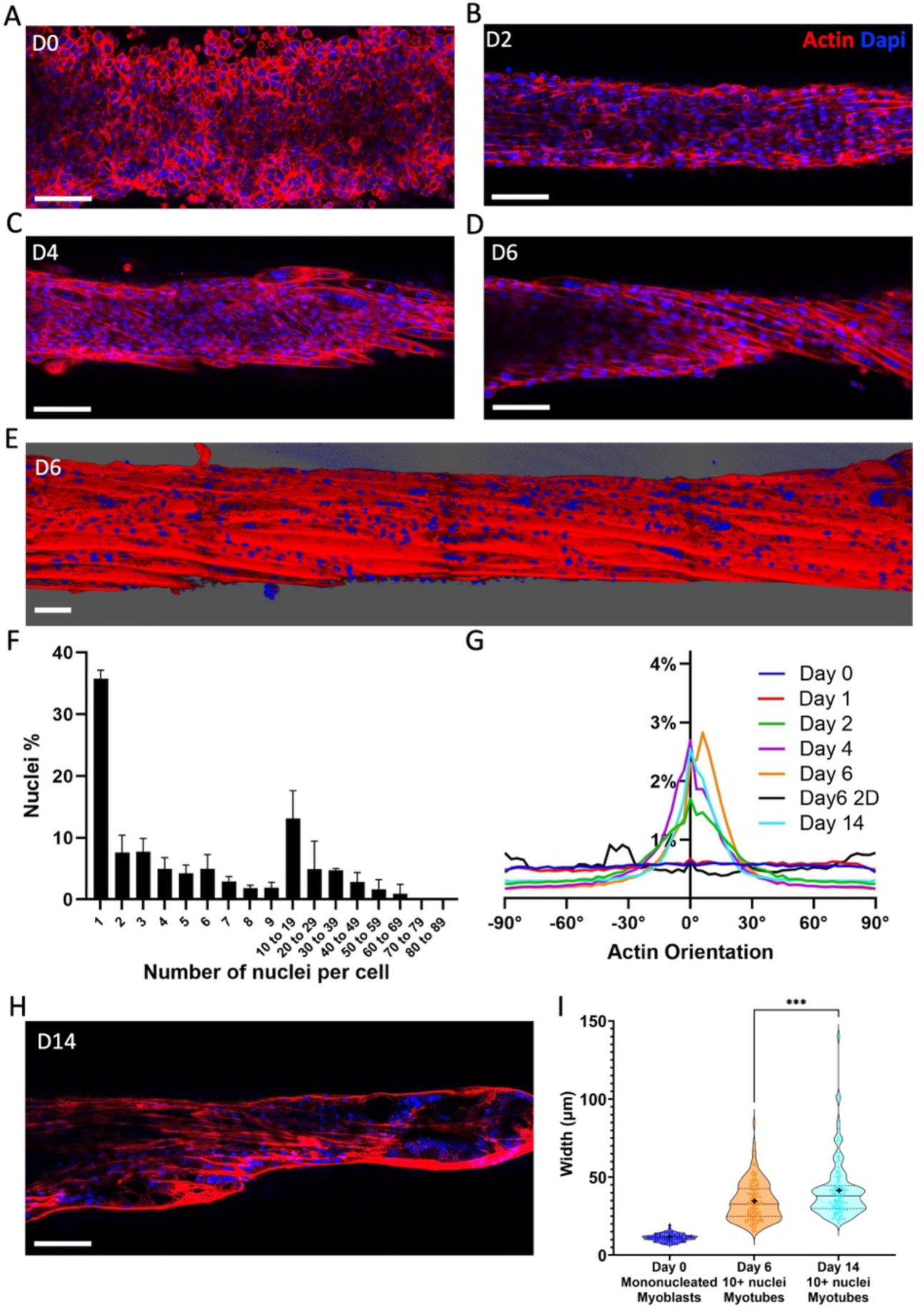
Constraining the tissue induces anisotropic differentiation and cell fusion. **A-D** Confocal imaging of a z slice of the internal cellular structure of a tissue made from C2C12 cells at day 0 (A), day 2 (B), day 4 (C) and day 6 (D). **E** 3D reconstruction from confocal imaging of a fiber at day 6 (actin in red and nuclei in blue). **F** Distribution of nuclei fusion in terms of nuclei percentage at day 6 in the 3D tissues (quantified for 3 samples, 9216 nuclei in total). **G** Distribution of actin filaments orientation in 3D tissues on day 0 (n=3), day 1 (n=4), day 2 (n=3), day 4 (n=4), day 6 (n=5), day 14 (n=4) and in 2D on day 6 (n=3). For each condition, the curve represents the average distribution across all samples. **H** Confocal imaging of a z-slice of the internal cellular structure of a tissue made from C2C12 cells at day 14. **I** Distributions of the widths of mononucleated myoblasts in tissues that were just bioprinted on day 0, and of myotubes with more than 10 nuclei in 3D tissues cultured between two needles for 6 (n=3) and 14 days (n=4). Each data point on the graph represents the measurement of one cell width. Scale bars = 100 µm.

### Stretching improves muscle differentiation

Mechanical cues are omnipresent in skeletal muscle tissue maturation^44–47^, and muscle development and functioning largely depend on being subjected to mechanical stretching constraints. The advantage of the magnetic tissue trapping system is its ability to stretch the anisotropic magnetic tissue between the needles at will. To achieve this, the chip containing the eight compartments was mounted on a translation stage with 3D-printed parts (Fig. 4A, Supplementary Fig. 16), allowing the distance between the needles to be increased, thereby stretching the clamped tissues. Tissue constructs were stretched by up to 100% in a single step on day 1. Remarkably, the central fiber can double in length without detaching or breaking (Fig. 4B, supplementary movie 3), leading to an immediate reorganization of the tissue on day 1 (Fig. 4C). Cells that are still isotropically organized in the absence of stretching (Fig. 4D, Supplementary Fig. 10A-10C) rapidly elongate and align when the tissue is stretched by 50% (Fig. 4E, Supplementary Fig. 10D-10G), with even more pronounced alignment at 100% strain (Fig. 4F, Supplementary Fig. 10H-10J). At 100% stretching on day 1, the distribution of actin filament orientation resembles that observed on day 6 without stretching, with 52 % of the actin filaments oriented within ±15° of the fiber. This indicates that stretching places the tissue in conditions that match a more advanced stage of differentiation. As the stretched tissues continue to mature overtime (Supplementary Fig. 11C-11J, 12D-12H), they not only become thinner by day 6 (Fig. 4G-4I, Supplementary Fig. 6, 13E-13K, and 17), but also display better alignment and cell fusion compared to tissues that were not stretched. Furthermore, twisting was no longer observed under stretching, and the distribution of actin filament orientation became narrower, with 66 % of the filaments aligned within ±15° of the fiber at day 6 (Fig 4J). Conversely, tissues cultured in a non-adherent well, totally free from constraints, exhibit no anisotropy in actin orientation (less than 19% of the filaments within ±15° no matter the reference direction chosen) and show minimal fusion (Supplementary Fig. 15D-15I). In tissues stretched by 100%, at day 6, 24% of the original myoblasts are still mononucleated while 76% have fused into myotubes (Fig. 4K), representing a significant improvement compared to non-stretched tissues. Stretching also increased the proportion of nuclei belonging to cells with more than 10 nuclei, reaching 38%. Maximum cell fusion and elongation are also improved, with some cells containing over 80 nuclei and measuring up to 2 mm in length (Supplementary Fig. 14B).

**Figure 4.**
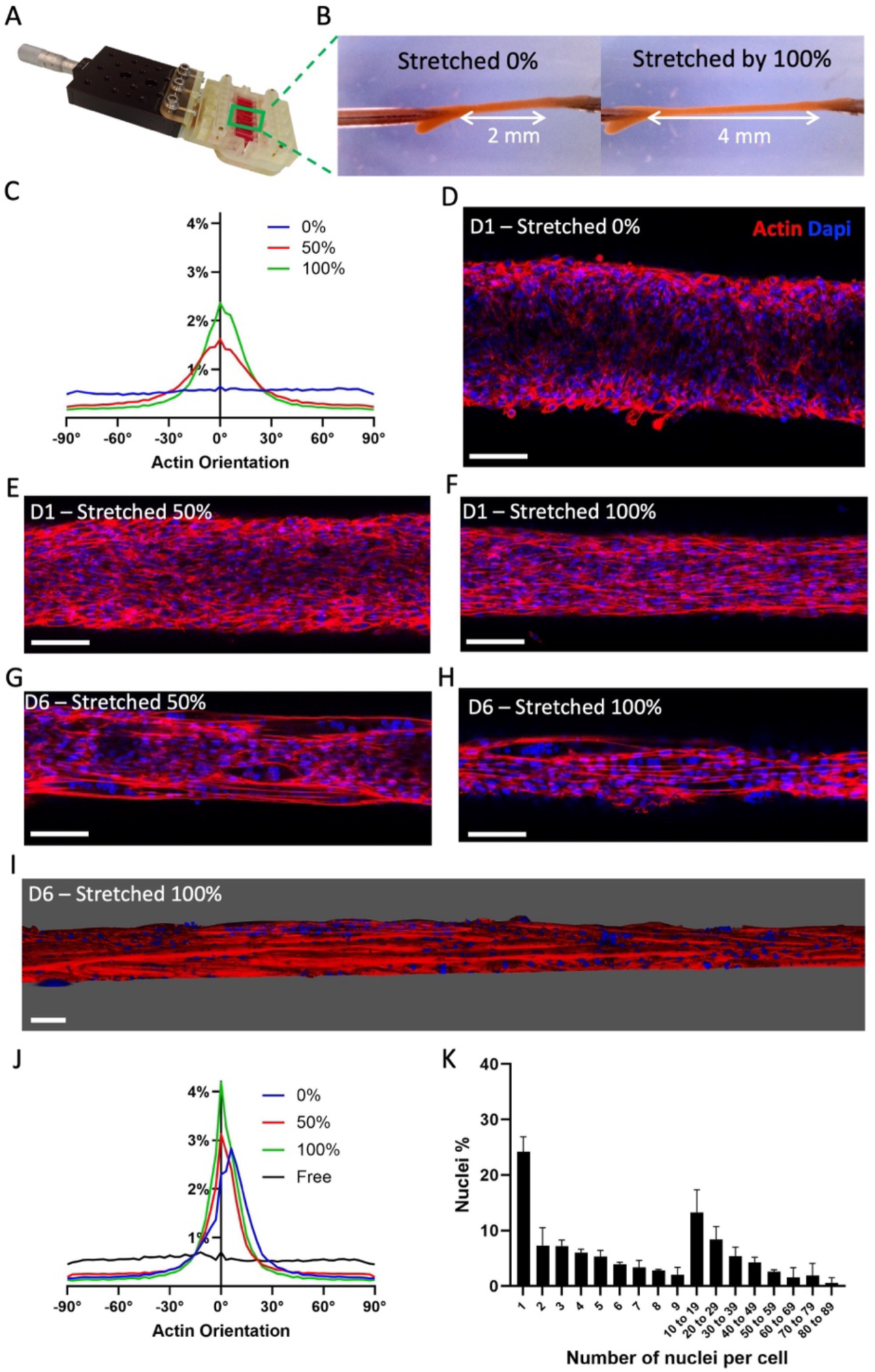
Anisotropy and cell fusion are enhanced by stretching the tissues. **A-B.** Stretching system used to stretch the chips (A) and therefore the tissues (B). **C** Distribution of the actin filaments orientation in tissues stretched by 0% (n=4), 50% (n=5) and 100% (n=4) at day 1. For each condition, the curve represents the average distribution across all samples. **D-H** Confocal imaging of a z slice showing the internal structure of a C2C12 tissue at day 1 stretched by 0% (D), 50% (E), or 100% (F) and at day 6 after 50% (G) and 100% (H) stretching at day 1 (actin in red and nuclei in blue). **I**. 3D reconstruction from confocal imaging of a tissue stretched by 100 % on day 6. **J** Distribution of the actin filaments orientation at day 6 in trapped 3D tissues stretched since day 1 by 0% (n=5), 50% (n=3) and 100% (n=4) and in 3D tissues cultured free of constraints (n=3). For each condition, the curve represents the average distribution across all samples. **K** Distribution of nuclei fusion in terms of nuclei percentage at day 6 in 3D tissues stretched by 100% (quantified for 3 samples, 12522 nuclei in total). Scale bars = 100 µm.

To further assess the mechanical stresses in the trapped tissues and examine their role in myogenesis, we embedded within the tissues deformable 42±5 µm spherical polyacrylamide beads coated with fluorescent poly-L-lysine for cell adhesion (Fig. 5A-5D, Supplementary Fig. 18). These deformable beads were previously used to measure internal stresses in tissues, such as in zebra-fish embryos^48^. Confocal imaging provided the aspect ratio (Fig. 5E) and orientation of the beads relative to the direction of the tissue (Fig. 5F). Notably, stretching the tissue at day 1 increases the aspect ratio of the beads to 1.80±0.14, and the orientation distribution of the beads becomes more aligned with the tissue direction. Remarkably, the aspect ratio of beads in non-stretched conditions at day 6 was 1.88±0.31, close to the 1.80 value found at day 1 immediately after stretching, confirming that stretching places the tissue in a more advanced maturation stage, at least from a mechanical point of view.

**Figure 5.**
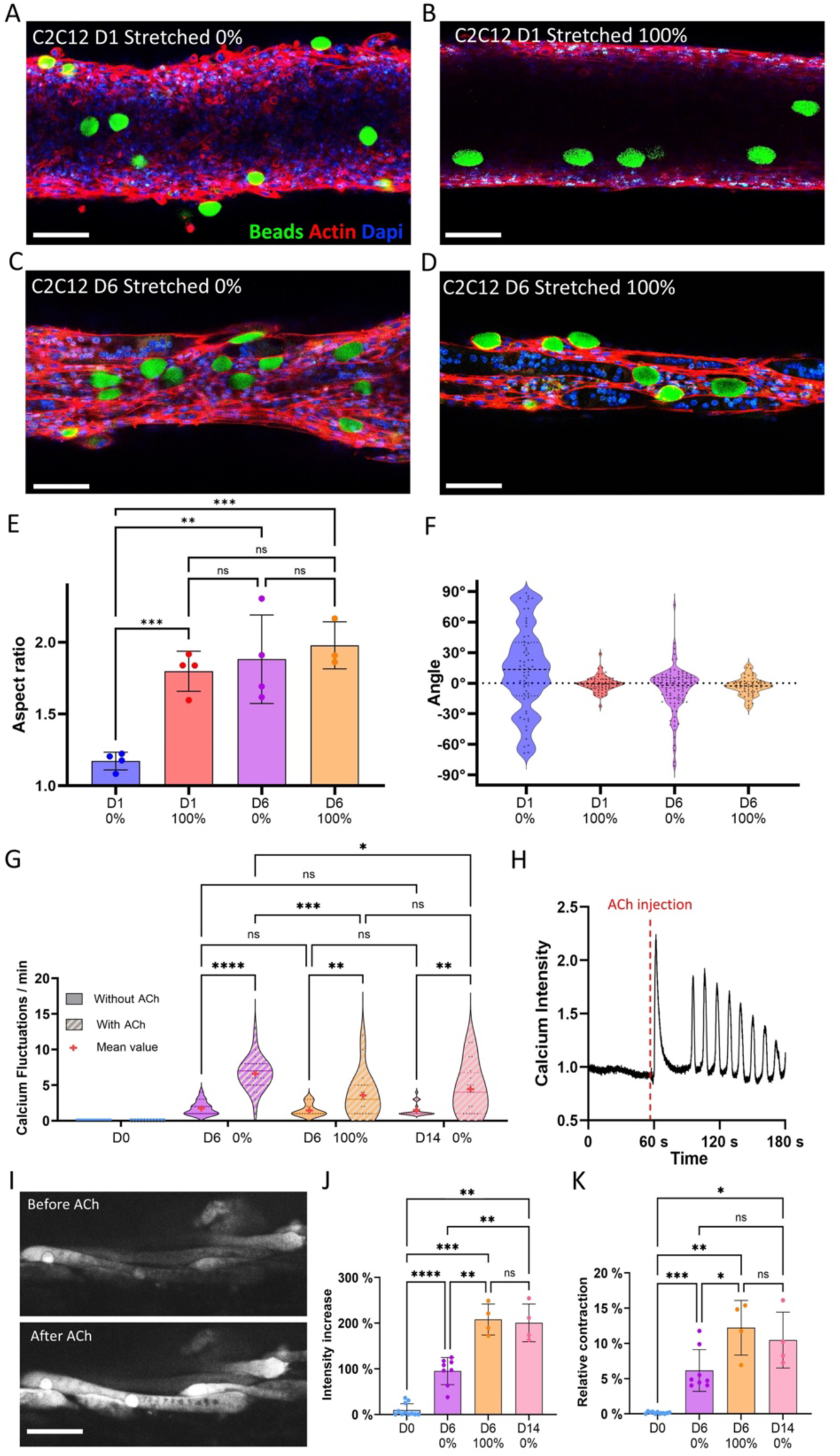
Deformable beads and calcium imaging reveal stresses and contractility are similar in early stretched tissues and differentiated tissues. **A-D** Confocal images of polyacrylamide beads in tissues made from C2C12 cells on day 1 stretched by 0% (A) and 100% (B), and on day 6 with an initial stretching of 0% (C) and 100% (D) (beads in green, actin in red and nuclei in blue). **E-F** Aspect ratio (E) and distribution of the orientation (F) of beads in tissues on days 1 and 6 whether they were stretched by 100% or not. Data points represent the average bead aspect ratio within one sample, and the angles of all beads across all samples. **G** Distribution of the frequencies of cellular calcium fluctuations in tissues on day 0, day 6 (whether stretched by 0% or 100%), and day 14 (non-stretched) before and just after treating them with 1mM of ACh. For each tissue, measurements were taken from 5 different cells exhibiting fluctuations, before and after ACh treatment, and each of these measurements was represented as one point on the plot. Samples on day 0 showed no fluctuations, but a line at 0 was placed for reference. **H-I** Kymograph (H) of the calcium signal in a cell and spinning disk microscope imaging (I) from a non-stretched tissue at day 6 before and after exposure to 1mM of ACh. **J-K** Relative increase of signal intensity (J) and tissue contraction (K) after treating tissues collected on day 0, day 6 (whether stretched by 0% or 100%) and day 14 (non-stretched) with 1mM ACh. Scale bars = 100 µm.

Generation of contractile force and calcium transients in response to chemical stimulation is also a key functional signature of skeletal muscles. To complement the mechanical characterization provided by the beads with information on functionality, calcium imaging was performed on tissues with and without the addition of acetylcholine (ACh), a neurotransmitter that induces an influx of calcium ions in skeletal muscle, triggering contraction^49^. For optimal visualization, tissues were detached from the needles and imaged with a spinning-disk microscope. While the calcium signal in cells from tissues imaged immediately after bioprinting (day 0) remained constant, the tissues exhibited some fluctuations in their calcium activity after 6 days of culture (Fig. 5G), even without ACh. After treating them with 1mM of ACh, the control samples at day 0 showed no response, except for a slight increase in signal (Supplementary Fig. 19A). However, in 6 and 14-day old tissues, ACh treatment not only induced increased calcium fluctuations, as shown in the kymographs in Figure 5H and supplementary Fig.19, but also enhanced the overall intensity of the calcium signal within the tissues (Fig. 5I-5J) and triggered tissue contraction (Fig. 5I and 5K). While the frequency of fluctuations increased the most in non-stretched tissues at day 6, with an average of 6.7 fluctuations per minute (Fig. 5G), both the relative increase in signal intensity and tissue contraction following ACh treatment were twice as high in stretched tissues. Notably, non-stretched tissues cultured for 14 days exhibited responses closer to those of stretched tissues, supporting the hypothesis that stretching could help tissues reach later stages of differentiation faster. One possible explanation for the greater increase in fluctuations in non-stretched tissues is that in stretched tissues, while some cells exhibited increased calcium fluctuations (Supplementary Fig. 19B), others showed significantly higher calcium intake (Supplementary Fig. 19C). This could lead to tetanic contraction and suggests that stretched tissues may be more sensitive to ACh.

### Transposing to myoblasts derived from human induced pluripotent stem cells

The two-fold method of magnetic bioprinting and tissue trapping was then applied to myoblasts derived from human induced pluripotent stem cells (hiPSC) to validate its effectiveness in a more clinically relevant model. A transgene-free differentiation protocol was used to generate a homogenous population of myogenic progenitor cells from a healthy hiPSC line within 4 weeks. Magnetic labeling of these hiPSC-derived myogenic cells resulted in each cell internalizing approximately 17 pg of iron (Supplementary Fig. 20). Figures 6A and 6B, along with supplementary Figure 21 demonstrate that M-hiPSC can be magnetically bioprinted and that the resulting tissues are viable (94±2% of live cells). These tissues derived from M-hiPSC could be clipped onto the magnetic needles (Fig 6C) and successfully cultured for 6 days (Fig. 6D, Supplementary Fig. 22), proving the successful transposition of the technique to M-hiPSC. The central fibers once again exhibited a decrease in width over time (Fig. 6E), however this reduction was more pronounced than in tissues derived from mouse myoblasts, reaching about 40% of their original width after 3 days, compared to approximately 60% in the latter (Fig. 2E). Immunostainings were performed at day 6 to assess specific skeletal muscle markers. Figure 5F and supplementary figure 23A-E confirmed the presence of desmin, a muscle-specific protein^50^, throughout the tissue, as well as myosin heavy chains (Supplementary Fig. 24A). Pax7, an indicator of satellite-like cells essential for muscle development and repair^51^,was found in 12.1±0.5% of cells in the 6-day-old 3D tissues (Fig. 6G-6H, Supplementary Fig. 25A-B), a proportion similar to that observed in 2D controls (Supplementary Fig. 26). To evaluate the number of myoblasts that started differentiating in the tissues, myogenin, a transcription factor for muscle differentiation^52^, was analyzed (Fig. 5I, supplementary Figures 27A-C and 28A-B). At day 6, 73.7±7.2% of cells were positive for myogenin, representing a significant increase compared to 2D controls (Fig. 6J, Supplementary Fig. 29). To gain deeper insight into the internal architecture of the cells, Figure 6K and supplementary Figure 30 show the alpha-actinin filaments, which highlight the presence of striations - the hallmark of the sarcomeres^53^, the contractile units of muscle tissue. Finally, about half of the 3D samples showcased sporadic spontaneous contractions by day 6 (Supplementary Movie 4), further validating the presence of the contractile apparatus in the muscle cells.

**Figure 6.**
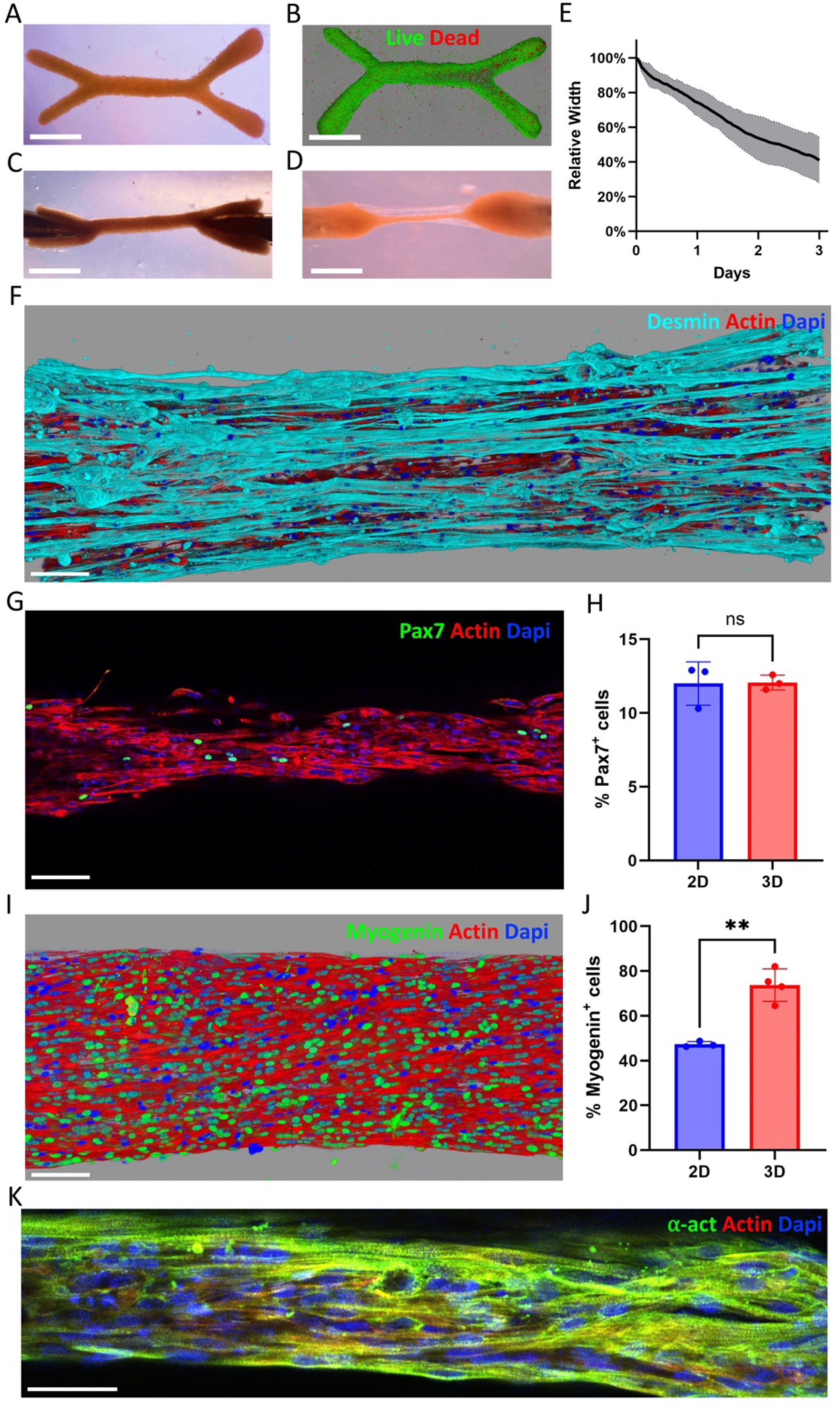
Differentiation is improved in 3D tissues bioprinted from M-hiPSC. **A-D** Wrench shaped tissue patterned with M-hiPS imaged (A) after patterning, (B) with confocal imaging after a live dead assay (live cells in green, dead cells in red), trapped between needles at (C) day 0 and (D) day 6. Scale bar = 1 mm. **E** Evolution of the central fiber’s width from day 0 to day 3 (n=3). **F** 3D reconstructions from confocal imaging of a 6-day old tissue immunostained in cyan for desmin. **G-H** Confocal imaging of a 6-day old tissue immunostained in green for Pax 7 (G) and proportions of Pax 7 positive cells (H). **I-J** 3D reconstruction from confocal imaging of a 6-day old tissue immunostained in green for myogenin (I) and proportions of myogenin positive cells (J). Scale bar = 100µm. **K** Confocal imaging of a 6-day old tissue immunostained in green for alpha-actinin. Scale bar = 50 µm. (Actin in red, dapi in blue).

The deformation of elastic beads embedded in tissues made of M-hiPSC (Supplementary Fig. 31A-H) also evidenced the stresses the cells were developing within the tissues over time. Although the beads exhibited the highest aspect ratio (Supplementary Fig. 31I) and anisotropy (Supplementary Fig. 31J) at day 6, most of their deformation along the tissue’s axis appeared to occur within the first 3 days of differentiation.

### Stretching improves functionality and specialization

Interestingly, some samples exhibited sporadic spontaneous contractions as early as day 3. Combined with the rapid thinning of the tissue (Fig. 7A) and the significant deformation of the beads by day 3, this points to fast differentiation. We thus investigated the effect of stretching on tissues cultured for 3 days, across three different M-hiPS cell lines (referred to as M-hiPSC 1, for the first cell line studied, 2 and 3 for the two additional ones). Magnetic trapping of tissues bioprinted with M-hiPSC 2 and 3 are illustrated in Supplementary Fig. 32. Tissues made from all three M-hiPS cell lines were successfully stretched by 100% on day 1(Fig 7B) and formed thin fascicle like structures by day 3. Alpha-actinin striations were observed in all conditions at day 3 (Fig. 7C-7D), indicating that the contractile apparatus is already in place after 3 days, which was further assessed through calcium imaging. At day 0, fluctuations were present in all cell lines, with an average frequency within flickering cells of 4.7 fluctuations per minute (Fig. 7E, supplementary Fig. 33A and Movie 5). This value significantly increased after 3 days, reaching 13.9 and 10.2 fluctuations per minute for samples stretched by 0% and 100%, respectively (Fig. 7E, supplementary Fig. 33B and Movie 6). Upon 1mM ACh treatment, fluctuations consistently increased for non-stretched tissues, reaching more than 20 fluctuations per minute on average at day 3. ACh treatment also led to an increase in signal intensity (Fig. 7F, supplementary Fig. 33C) and contraction (Fig. 7G-7H) of all 3-day old tissues, with effects being twice as high on average in stretched tissues (Supplementary Movie 7) compared to non-stretched tissues.

**Figure 7.**
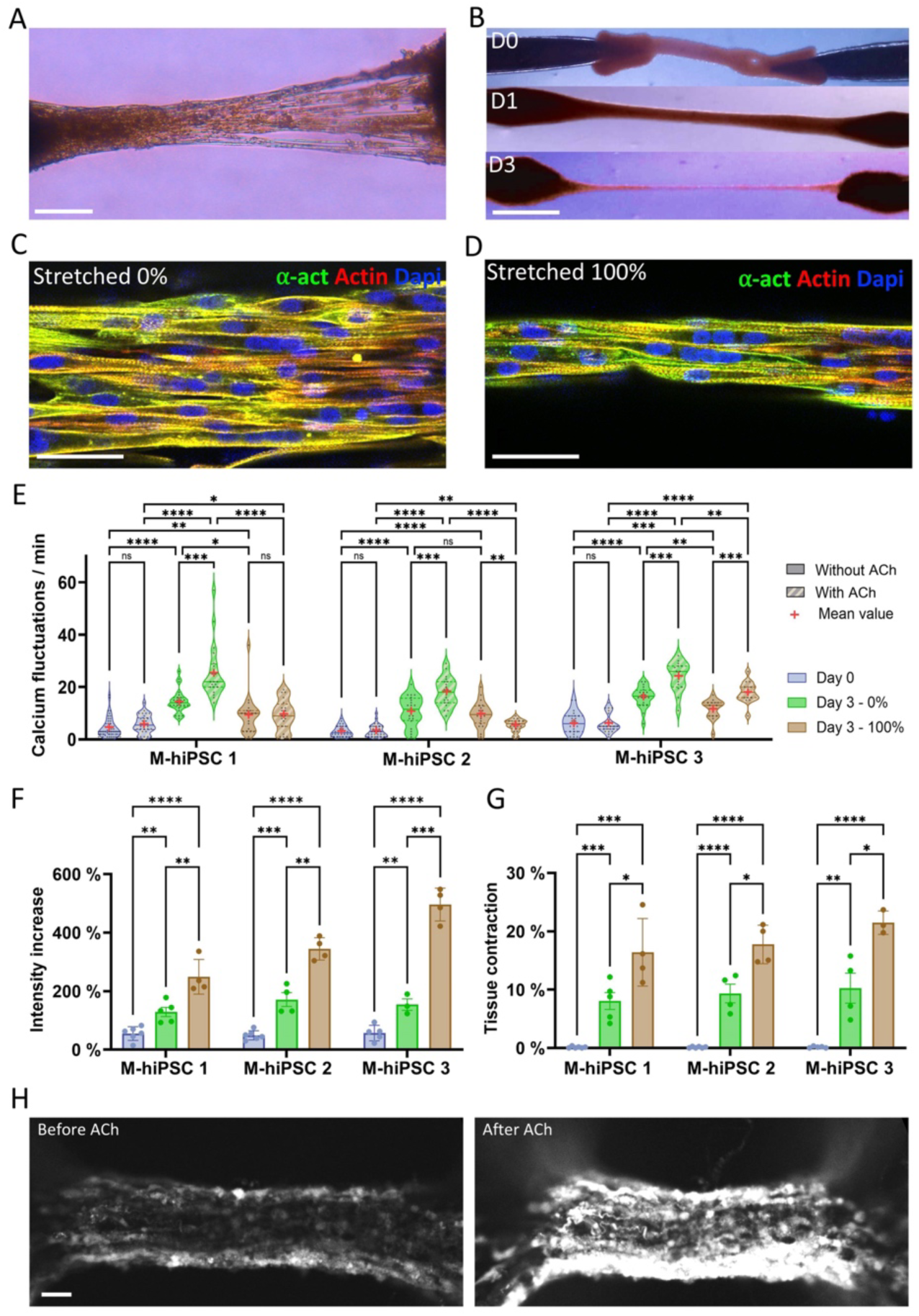
Stretching tissues made from the three different M-hiPS cell lines increases tissue contractility. **A** 3D tissue made with M-hiPSC-1 on day 3. Scale bar = 200 µm. **B** Trapped tissue on day 0, and after 100% stretching on days 1 and 3. Scale bar = 1 mm. **C-D** Confocal imaging of 3-day-old M-hiPSC 1 tissues immunostained in green for alpha-actinin with (C) 0% and (D) 100% stretching. **E** Distribution of the frequencies of calcium fluctuations in M-hiPS cells from tissues on day 0 and on day 3 whether they were stretched by 0% or 100%, before and just after treating them with 1mM of ACh. For each tissue, measurements were taken from 5 different cells exhibiting fluctuations, before and after ACh treatment, and each of these measurements was represented as one point on the plot. **F-G** Relative increase in signal intensity (F) and tissue contraction (G) after 1mM ACh treatment in tissues collected on day 0 and day 3 whether they were stretched by 0 or 100%. **H** Spinning-disk microscope imaging from a M-hiPSC 2 tissue stretched by 100% at day 3, before and after exposure to 1mM of ACh. Scale bar = 50 µm.

Immunofluorescence staining and confocal imaging were next performed to better characterize 3-day old tissues. Cells are expressing desmin (Fig. 8A-C, Supplementary Fig. 23F-Q) and myosin heavy chains (Supplementary Fig. 24.B-D) in all conditions. Stretching then appears to significantly impact the populations of myogenin positive (Fig 8D-F, Supplementary Fig. 27D-O and 28C-K) and Pax7 positive (Fig 8G-I, Supplementary Fig. 25C-J) cells. In tissues made from M-hiPSC 1, the number of myogenin positive cells at day 3 increases with stretching, rising from 64.8±4.3% without stretching to 77.1±3.2% with 100% stretching (Fig. 8J), and stretching doubles the number of Pax7 positive cells, which goes from 11.2±1.1% without stretching to 21.7±1.5% with 100% stretching (Fig. 8K). These increases were also observed in M-hiPSC 2 and 3 (Fig. 8J-8K). EdU staining confirmed that most Pax7 positive cells in stretched samples are quiescent (Fig. 8L), with only 9.9±2.7% proliferative cells among Pax7-positive cells across the three cell lines, which is similar to the 7.9±1.3% in non-stretched 3-day old samples and 9.6±0.9% at day 0. This further reinforces the idea that stretching not only increases the number of cells starting to differentiate but also promotes the presence of satellite-like cells.

**Figure 8.**
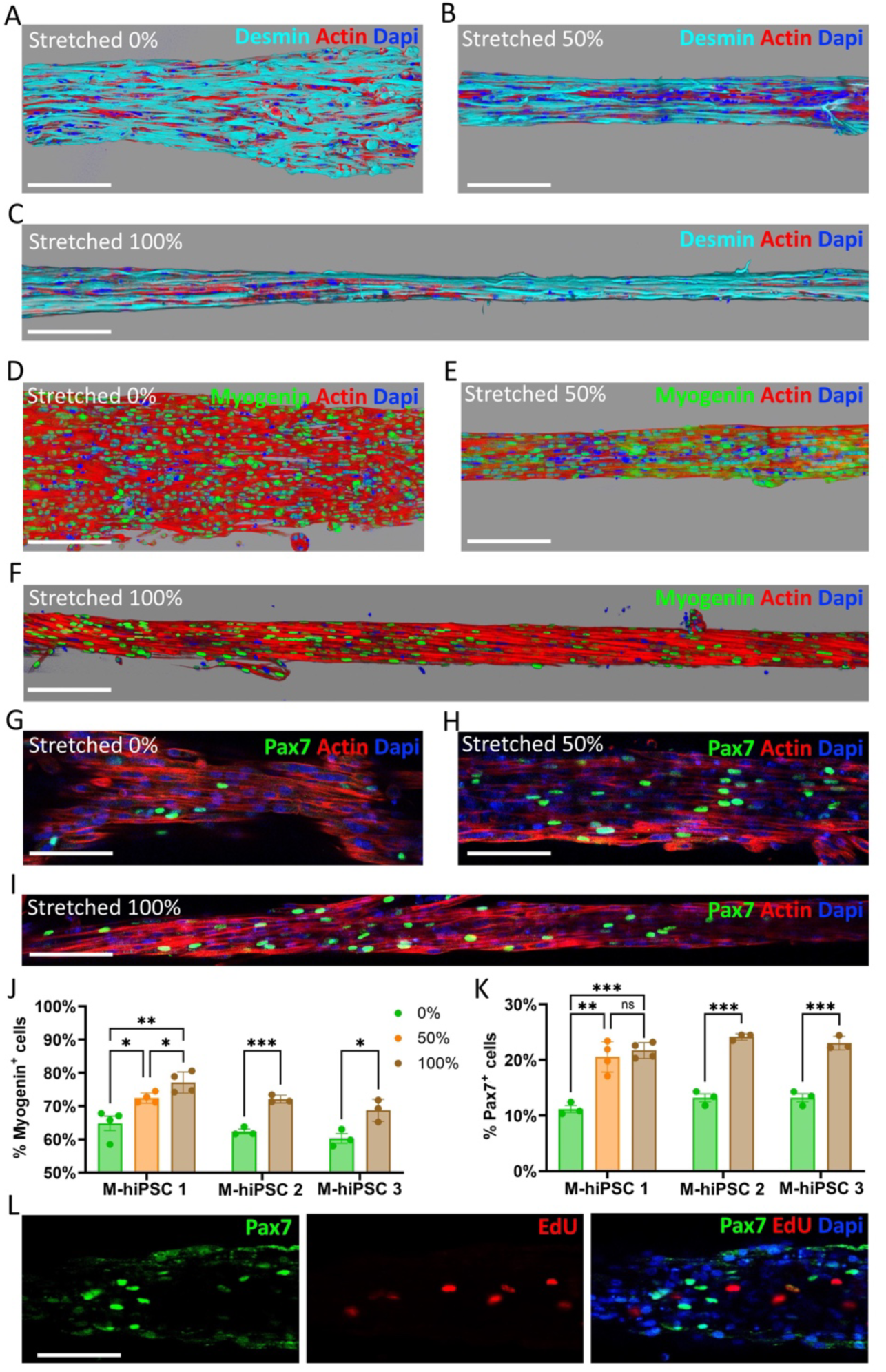
Stretching tissues made from M-hiPSC increases the number of myogenin and Pax7 positive cells. **A-C** 3D reconstruction from confocal imaging of 3-day old M-hiPSC 1 tissues immunostained in cyan for desmin with (A) 0%, (B) 50% and (C) 100% stretching. Scale bar = 200 µm. **D-F** 3D reconstruction from confocal imaging of 3-day M-hiPSC 1 old tissues immunostained in green for myogenin with (D) 0%, (E) 50% and (F) 100% stretching. Scale bar = 200 µm. **G-I** Confocal imaging of 3-day old M-hiPSC 1 tissues immunostained in green for Pax 7 with (G) 0%, (H) 50% and (I) 100% stretching. Scale bar = 100µm. (Actin in red, nuclei in blue). **J-K** Proportions of (J) myogenin positive cells and (K) Pax7 positive cells at day 3 in tissues made from all three M-hiPS cell lines whether they were stretched by 0%, 50% or 100% at day 3. **P** Confocal imaging of the different colour channels of a 3-day old M-hiPSC 1 tissue stretched by 100% with proliferative cells labelled with EdU and then immunostained for Pax 7 (Pax7 in green, EdU in red, nuclei in blue).

Finally, gene expression analysis was performed at day 3 on M-hiPSC 1 cells to quantify the levels of four different isoforms of myosin heavy chains: one embryonic form (Myh3), one neonatal form (Myh8) and two adult forms (Myh1 and Myh2) (Supplementary Fig. 34). Both the embryonic and neonatal isoforms were upregulated in non-stretched 3D samples, but their expression decreased in tissues stretched by 100%. Without stretching, Myh1 expression was similar in 2D and 3D, while Myh2 was slightly lower in 3D. However, stretching positively impacted the expression of these two adult isoforms of myosin, as both Myh1 and Myh2 were upregulated by stretching 3D samples by 100%.

## Discussion

In this work, we demonstrate the engineering of 3D mature muscle tissue using magnetic forces. These forces serve (1) to precisely bioprint the tissue into any desired shape without the need for a supporting matrix, and (2) to maintain its shape or even stretch it, thereby promoting muscle maturation.

The next-generation magnetic bioprinting method with high-resolution NiFe patterns produces shape-controlled 3D tissues in just 3 hours. The use of NiFe patterns as customized magnets enhances previous magnetic bioprinting methods^29,34–36^ by providing improved control over the geometry and competes effectively with traditional bioprinting approaches^16–18^ enabling precise shape control. Despite the significant advancements brought by bioink-based bioprinting to 3D tissue engineering, it still faces limitations, such as external stresses applied to cells during the printing process and the lack of physiological relevance of certain bioinks. Additionally, matrix can limit cell density^24^, posing a challenge when differentiating muscle precursor cells into muscle fibers. Magnetic bioprinting offers distinct advantages. Although the absence of matrix makes tissue cohesion more challenging and necessitates the use of higher number of cells, this approach allows for the creation of tissues composed with densely packed cells. This high cell density reduces the dependency on an extracellular matrix, relying instead on adhesive molecules produced by the cells themselves. Consequently, magnetic bioprinting enables the fabrication of tissues that more closely mimic the cellular composition and density of native tissues.

The magnetic properties of the cells were not only used for bioprinting tissues with controlled geometry but also for trapping the tissue between two magnetic attractors. This magnetic clipping function complements remote magnetic actuation techniques, which have primarily been applied for targeting therapeutic agents, controlling cell migration^30,54,55^ or triggering mechanotransduction^36,38,39^. One major advantage of magnetic clipping is its ability to maintain the tissue’s geometry over time. Preserving the anisotropy of a tissue, such as muscle, is crucial for its function, but remains challenging^10^ for contractile tissue, as evidenced in Supplementary Figures 7D and 15D-I without the clipping function. Systems where the tissue is molded in a tube or extruded typically provide good anisotropy over the entire tissue^56,57^. However, molded gels can be colonized^29^, may not be biomimetic, or are simply used as sacrificial molds^57^. The magnetic clipping approach shares similarities with systems where tissue matures around two posts^16,23–27,58–64^. This post-method, introduced over two decades ago^24^, has become a stapple of muscle tissue engineering. Despite numerous variations, few alternative approaches have emerged, aside mostly from bioprinting^17^ and tube molding^56,57^ (Table 1). All these methods rely on extracellular matrix for tissue cohesion. For those that do not, only tissues at much smaller scale - typically in the range of a few hundred micrometers^26^ - have been successfully achieved (Table 1). Thus, having a novel method to produce scaffold-free tissues at the centimeter scale with precise control over both their initial and final shape using remote magnetic actuation could be beneficial for skeletal muscle tissue engineering. This is particularly advantageous when tissue maturation can be further enhanced through stretching.

**Table 1.** Table comparing various parameters of the different skeletal muscle tissue engineering constructs referenced in this study.

| Ref.<sup>a</sup> | Cell type<sup>b</sup> | Days<sup>c</sup> | Tissue engineering Approach | Scaffold | Size<sup>d</sup> (mm) | Mechanical stimulations | Other Stimulations | Force evaluation |
| --- | --- | --- | --- | --- | --- | --- | --- | --- |
| <sup>16</sup> | Mouse mesoangioblasts - Muscle-derived human mesenchymal stem cells | 30 | Extrusion of photopolymerized fiber on C-shape support | PEG-Fibrinogen (8mg.mL<sup>-1</sup>), 0.1% w.v<sup>-1</sup> Irgacure 2959 | 40 | No | No | No |
| <sup>17</sup> | Human muscle progenitor cell | 9 | Extrusion based bioprinting with sacrificial gelatin bioink | Fibrinogen (20 mg.mL<sup>-1</sup>), gelatin (35 mg.mL<sup>-1</sup>), hyaluronic acid (3 mg.mL<sup>-1</sup>), 10% (v/v) glycerol | 15 | No | No | No |
| <sup>23</sup> | Primary rat myoblasts | 42 | Seeding cells on a fibrin gel with two anchors | 500μL medium with thrombin (10U.mL<sup>-1</sup>) with 200μL fibrinogen (20 mg.mL<sup>-1</sup>) | 12 | No | Electrical (15V, 1.2ms single pulse & 1s train of 1.2ms 10-150Hz pulses) | Force transducer |
| <sup>24</sup> | Primary human skeletal muscle cells | 16 | Casting a cell-loaded gel in a well with two pins | Collagen I (0.8mg.mL<sup>-1</sup>), Matrigel (1:6 v/v) | 25 | 10% / 4 days strain & later cycles of 3 sets of 5 2s pulses (5% strain for 2 days, 10% for 2 days, 15% for 4 days) | Cytochalasin D (2-5μM) | Force transducer |
| <sup>25</sup> | C2C12 - M-hiPS cells | 28 | Injection of cell loaded gel in microfluidic chamber with two pillars, with motoneuron spheroid in another chamber | Collagen I (2.4mg.mL<sup>-1</sup>), Matrigel (10%) | 1.5 | No | Electrical (0.5-4Hz, 12V) Glutamic acid (0.1-5mM), tetrodotoxin (1μM), bosutinib (100 μM) & rapamycin (200 nM) | Pillar deflection |
| <sup>26</sup> | Immortalized (healthy and pathological) and primary human myoblasts | 14 | Seeding cells on pillars micropatterned with Matrigel | No matrix | 0.2 | No | ACh (20μM) | Pillar deflection |
| <sup>27</sup> | C2C12 cells modified for optogenetic control | 21 | Injection of a cell loaded gel in a well with two pillars | Collagen I (2mg.mL<sup>-1</sup>), Matrigel (10%) | 1 | No | Optical (1.5 W collimated blue light or 300 W lamp with 472nm filter) | Pillar deflection |
| <sup>29</sup> | C2C12 | 7 | Magnetic alignment of cells and spheroids in a gel | Collagen I (2mg.mL<sup>-1</sup>), Matrigel (10%) | 10 | 20% strain | No | No |
| <sup>56</sup> | C2C12 | 28 | Seeding cells in a microfluidic tube molded in a collagen gel | Collagen I (6mg.mL<sup>-1</sup>) | 4 | No | No | No |
| <sup>57</sup> | C2C12 | 14 | Molding a cell laden gel in a microfluidic tube in a sacrificial gel | Fibrinogen (5 mg.mL<sup>-1</sup>), Thrombin (1 U.mL<sup>-1</sup>), Matrigel (20% v/v) | 5 | No | No | No |
| <sup>58</sup> | C2C12 | 13 | Seeding cells onto DegraPol® strips | DegraPol® strips | 31 | 48h stretching at 0.02 mm/h, then cycles of 3 sets of 5 1mm 2s pulses; Max. strain: 6.7% | No | No |
| <sup>59</sup> | C2C12 | 8 | Casting a cell-loaded gel in a cylindrical shape with two anchors | Fibrinogen (3 mg.mL<sup>-1</sup>), aprotinin (5 mg.mL<sup>-1</sup>) mixed 1:1 with thrombin (2 U.mL<sup>-1</sup>) | 15 | 50% strain | No | No |
| <sup>60</sup> | C2C12 | 9 | Molding a cell-loaded gel in a ring shape and mounting it onto spools | Fibrinogen (20 mg.mL<sup>-1</sup>), thrombin (0.6 U.mL<sup>-1</sup>) | 33 | Cycles of 10% strain for 6 h, 18 h at 3%, for 6 days | No | No |
| <sup>61</sup> | C2C12 | 16 | Casting a cell-loaded gel in a well with two pillars | 85% collagen I (2 mg.mL<sup>-1</sup>) | 24 | 15% / 1h strain, 15% for 2h and then 0% | Electrical (3.6 V/mm 1.2 ms pulse & 1s 100 Hz) | Force transducer |
| <sup>49</sup> | Muscle progenitor cells induced from hiPS cells - Primary human muscle cells | 28 | Casting a cell-loaded gel in semi-cylindrical mold and anchoring it to nylon frame | Fibrinogen (4 mg.mL<sup>-1</sup>), Matrigel (20% v/v), thrombin (0.2U.mg<sup>-1</sup>) | 7 | No | Electrical (40 V/cm, 10 ms pulse & 1s 5-40 Hz) ACh (10mM) | Force transducer |
| <sup>62</sup> | Muscle progenitor cells induced from hiPS cells - Primary human myoblasts | 7 | Casting a cell loaded gel in a chamber with pillars | Fibrinogen (1-2 mg.mL<sup>-1</sup>), Matrigel (20% v/v), Thrombin (0.8U.mL<sup>-1</sup>) | 11 | No | Electrical (2.45 V, 1 or 20 Hz) Caffeine (100μM), verapamil (10μM), cardiotoxin (1.2μM), chloroquine (100μM) | Pillar deflection |
| <sup>63</sup> | Primary human myoblasts | 14 | Casting a cell loaded gel in wells of a 96-well plate with PDMS pillars | Fibrinogen (4 mg.mL<sup>-1</sup>, 40% v/v), Geltrex<sup>TM</sup> (20% v/v), thrombin (0.2 U.mL<sup>-1</sup>) | 4 | No | Electrical stimulation (10 V/cm, 0.5-20 Hz) Dexamethasone, IGF -1, cerivastatin (1-100nm), gemcitabine (32-320μM), d-tubocurarine (25 μM), ACh (2mM), Irinotecan (16-72μM) | Pillar deflection |
| <sup>64</sup> | Primary rat skeletal myocytes | 28 | Casting a cell loaded gel in circular mold, later transferred to holders | collagen I (0.4mg), Matrigel (10% v/v) | 8 | Mechanical “crush” injury | Electrical stimulation (4ms, 2.5-100Hz) 25 μg.mL<sup>-1</sup> cardiotoxin, 0.5 μmol/L γ-secretase inhibitor, 5 μmol/L IWR-1, 1 μmol/L CHIR99021, 100 mol/L Oxytocin, 100 nmol/L Angiotensin-II | Force transducer |
| * | C2C12 - M-hiPSC | 14 | Magnetic bioprinting and trapping | No matrix | 8 | Strain 100% | ACh (1mM) | Deformable beads |

<sup>a</sup>Reference of the study presenting this construct
<sup>b</sup>Skeletal muscle cell types or cell types meant to mature towards muscle tissue in the construct. Unless mentioned, cells were not pathological.
<sup>c</sup>Maximum *in vitro* culture time of muscle cells in 3D system (in days)
<sup>d</sup>Maximum dimension of the construct achieved (in mm)
\* Present study

The magnetic approach proposed here has successfully enabled the engineering of skeletal muscle tissue with a high level of differentiation within a few days, both with immortalized C2C12 cells and even with M-hiPS cells. This cell type has only recently been used to create 3D constructs^25,49,62^. Here, indicators of differentiation after a week, such as the presence of myogenin-positive cells, were markedly higher to those observed in M-hiPS 2D cultures. Human iPS-derived muscle cells exhibited spontaneous contractions after just three days of culture in our system, when they are usually reported after at least a week^49^. Functionality was assessed by calcium imaging in absence or presence of acetylcholine, with responses within just 3 days, comparable to those reported in other studies^49,63^ after more than a week with higher ACh concentrations. Other studies have also investigated functionality by examining the force response upon electrical stimulations^23,25,49,61–64^ (Table 1), which could also be explored in combination with mechanical stimulation in our system for future studies. Importantly, high levels of maturation in our systems as early as day 3 were further evidenced by the high percentage of myogenin-positive cells and the presence of clear sarcomeric striations in most cells. This crucial feature of muscle differentiation is usually expected after one week^49,62^ and is not always reported in muscle constructs^16,23,58,59,61^. One study^49^ showed that about 20% of nuclei belonged to striated cells after a week of culture, and 50% after 2 and 4 weeks. In contrast, we achieved here high levels after only 3 days of maturation. Furthermore, stretching significantly enhanced maturation, with alignment and differentiation improving in tissues stretched by 100%. Indeed, stretching further increased both the populations of myogenin and Pax7 positive cells after only two days in tissues generated from M-hiPSC. While the role of stretching on muscle cell differentiation has mostly been studied with C2C12 cells^29,58–61^ and with primary human skeletal muscle cells^24^ (Table 1), its impact on myoblasts derived from hiPS cells remains to be fully explored. This study shows that stretched samples from hiPS-derived myoblasts exhibit levels of myogenin positive cells similar to the ones observed after 2 weeks of culture in another study^49^. Moreover, the number of Pax7-positive cells doubles in stretched tissues compared to non-stretched ones at day 3, despite being quiescent and likely satellite-like. This, to the best of our knowledge, has not been observed before and would require further investigation to understand the impact of mechanical stress on Pax7 positive, and more specifically satellite, cells. The enhanced response to ACh in stretched samples also highlights the significant role of mechanical cues in myogenesis, with stretched samples contracting by approximately 20% - twice more than non-stretched samples - when stimulated with ACh after just 3 days. It is important to note that 100% stretching is far beyond physiological levels, especially in comparison to other approaches that rarely exceed 20% stretching (Table 1). However, in the context of *in vitro* tissue engineering, such a high strain could offer beneficial effects, and it does not require complex stretching machinery and protocols like other methods^58–61^. Due to its simplicity, our system does not offer direct force measurement through a force transducer^23,24,49,61,64^ or pillar deflection^25–27,62,63^. To assess the mechanical environment of the cells, we proposed an approach for stress evaluation in muscle tissue, originally developed for zebrafish embryos^48^, using deformable polyacrylamide beads embedded in the tissue. These beads revealed that stretching the tissues led to stresses and a mechanical state, similar to those in mature tissues, which could help explain why stretching enhances myogenesis and increases the number of Pax7-positive cells.

The magnetic approaches presented here could be further complexified for various biological applications, including disease modeling and drug screening, and could provide insights into the neuromuscular junction once innervated. For instance, magnetic bioprinting could be employed to create complex and precisely tuned cellular architectures, with innervation or vascularization integrated with the magnetic muscle constructs in co-culture approaches. The rapid magnetic bioprinting process and its potential to generate large and complex tissues offer promising opportunities for developing grafts, which could be monitored through MRI using the nanoparticles as contrast agents. A system of magnetic stents, similar to the magnetic needles, could be used to directly position these grafts *in vivo*.

## Materials and Methods

### Cell Culture

C2C12 cells were cultured in proliferation medium consisting of DMEM (Gibco) supplemented with 10% FBS (Dutcher) and 1% Penicillin-Streptomycin (Thermofisher).

M-hiPS cells were cultured in DMEM with 1000 mg/L D-Glucose (Stem Cell Technologies) supplemented with MyoCultTM-SF expansion 10X supplement (human, Stem Cell Technologies) and 0.1% Penicillin-Streptomycin on flasks coated with Matrigel (Corning). Both cell types were passaged once confluency reached between 50 and 80%, without exceeding passage 5.

### Myogenic differentiation of hPSCs

The three hiPSC lines used in the study were derived by Phenocell^®^ using the Epi Episomal iPSC Reprogramming Kit (ThermoFischer) from three different patient cell lines (Supplementary table 1). Informed consents were obtained from all the patients included in this study, complying with the ethical guidelines of the institutions and with the legislation requirements of the country of origin. Experimental protocols were approved by the french minister of health (2019-A02599-48).

hiPCs were used at passages P15-P20, maintained in StemMACS iPS-brew XF medium (Miltenyi Biotec) in vitronectin (Gibco)-coated culture dishes and were routinely tested for Mycoplasma contamination using a commercially available kit (MycoAlert, Lonza).

hiPSC skeletal muscle differentiation experiments were performed using media A, B, C, and D from the commercially available STEMdiff™ Myogenic Progenitor Supplement Kit (Stem Cell Technologies). Briefly, hiPSC colonies were dissociated into single cells with Tryple Express (Thermo Fischer Scientific) and seeded onto Matrigel (Corning)-coated 6-well plates at a density of 20,000 cells/cm² in medium A for 2 days. Cells were then switched to medium B (days 2-4), medium C (days 2-4) and medium D (days 6-30) with a daily medium change.

Day 30 skeletal myocytes were dissociated with Tryple Express and 285µg/mL Collagenase IV (Stem Cell Technologies) for 10 minutes, filtered on a 70 µm strainer and replated on Matrigel-coated flasks at a density of 15,000 - 20,000 cells/cm² in MyoCult™-SF Expansion Medium with 10 µM Y-27632 (Stemgent). When the culture reached 60 - 80% confluency, myogenic progenitors were harvested and cryopreserved.

### Terminal differentiation

C2C12 differentiation was triggered by switching their culture medium to differentiation medium, consisting of DMEM (Gibco) supplemented with 1% Horse Serum (Thermofisher) and 1% Penicillin-Streptomycin (Thermofisher).

M-hiPSC were differentiated into mature muscle cells using the MyoCult™ Differentiation Kit (human, Stem Cell Technologies) supplemented with 0.1% Penicillin-Streptomycin.

In both cases, growth medium was switched with differentiation medium on day 0, immediately after the cells were magnetically bioprinted.

### Magnetic Labeling

Cells were magnetically labeled with superparamagnetic maghemite iron oxide nanoparticles coated with citrate^29,65^. The nanoparticles were synthesized using the standard procedure of iron salts co-precipitation. In brief, 3 g of FeCl_3_ were mixed with 8.1 g of FeCl_2_ in 5 mL deionized water, supplemented with 25 % ammonium hydroxide as precipitating agent to achieve and alkaline pH of 10. The solution was heated to 90 °C for 20 minutes, then magnetically decanted, washed with acetone, washed with water, and centrifuged (8000 rpm for 10 min). The pellet was resuspended in 10 mL of deionized water. Subsequently, 2 g of citrate, dispersed in 20 mL of deionized water, was added to the solution, and the mixture was further heated at 90 °C for 1 hour. This final step is crucial for both the oxidation of nanoparticles into maghemite and the citrate coating which ensures colloidal stability. The nanoparticles diameter was measured by transmission electron microscopy to be 8 ± 1.7 nm. Citrate absorption was effective, resulting in a final negative zeta potential −35mV. The nanoparticles exhibited a saturation magnetization of 65 emu/g, without any hysteresis in the magnetization curve. Before administering the nanoparticles to cells, the stock solution was filtered-sterilized through a 200 nm filter.

C2C12 cells were labeled by incubating them in an RPMI (Gibco) solution containing nanoparticles at a concentration of 2mM iron and 2mM citrate. The citrate was included to prevent nanoparticle precipitation without affecting the solution’s pH. Incubation was performed for 30 minutes once daily over three days preceding magnetic bioprinting, as previously described^29^. During this process, cells progressively internalized the nanoparticles through the endocytic pathway.

The labeling procedure was modified for M-hiPS cells, as prolonged exposure to the 2mM iron and 2mM citrate solution caused cell detachment. These cells were labeled with the same nanoparticle solution, but for 15 minutes once daily over two days preceding magnetic bioprinting. To further enhance nanoparticles uptake, nanoparticles dispersed at 0.2 mM iron were also added to the cells’ regular culture medium at 0.2 mM iron overnight between the two labeling sessions.

### Magnetophoresis to quantify iron internalization

To measure the amount of iron internalized by the cells, magnetophoresis was performed^65^. Magnetically labeled cells were resuspended in PBS and the solution was placed next to a permanent magnet with a magnetic field B=0.145T and a gradient grad(B)=17 T.m^-1^. By recording the cells’ displacement towards the magnet and measuring their speed and diameter, the magnetic force applied to each cell was inferred, as it balances the Stokes drag force. This allowed for the determination of the magnetic moment and, consequently, the amount of iron internalized by each cell.

### Magnetic signal measurement

After bioprinting, tissues were cultured in differentiation medium in 500 µl Eppendorf tubes. The tubes fit in a bench-top magnetometer which was regularly used to measure magnetic signal of the tissues. The device assesses the magnetic response of the tubes content when exposed to two alternating fields, providing a value proportional to the amount of magnetic material within the sample. The details of the principle of this magnetic technique and its application are explained in reference^43^.

### Nickel iron pattern fabrication

A 70×50mm Corning glass slide was covered with a Ti-Cu layer (Ti 10nm, Cu 100nm) through magnetron sputtering (Plasmionique). The resulting conductive layer was spincoated with TI-Prime (Microchemicals) and then with 70µm of AZ 125nXT photoresist (Microchemicals). Photolithography with a chromium mask would then remove the resist in the shapes and positions of the desired patterns. These holes, where the copper was exposed, could be used as molds to grow NiFe (80:20) patterns through electrodeposition^29,39,66^. The substrate was immerged in a magnetically steered 30°C electroplating bath with 250g.L^-1^ NiSO_4_, 5g.L^-1^ FeSO_4_, 25g.L^-1^ boric acid, 2g.L^-1^ saccharin and 0.1g.L^-1^ sodium dodecyl sulfate. With the substrate acting as a cathode and a pure nickel anode (Goodfellow), a 7 mA.cm^-2^ current density resulted in NiFe growing by about 3µm.h^-1^ on the exposed copper on the substrate. Once the NiFe deposit reached a height of about 50µm, the resist and surrounding copper were removed using TechniStrip P1316 (Technic) at 70 °C. The patterns were then protected by spincoating the substrate with a 100µm layer of PDMS (Sylgard 184, 1:10 curing agent). Multiple wrench-shaped patterns were made on the same glass slide, and they were usually regrouped in groups of 6, packed together to fit in a 16mm wide circle.

### Magnetic bioprinting

Wells to support magnetic tissue formation were prepared from 1 cm thick PDMS (Sylgard 184, 1:10 curing agent) by punching out discs with an inner diameter of 16 mm and an outer diameter of 20 mm. These hollow disks were plasma-bonded to 100 µm thick 22×22mm VWR coverslip to create the wells.

Each well was treated with anti-adherence solution (StemCell Technologies) for 1 hour and then placed on top of a group of 6 patterns. Magnetically labeled cells were detached using TrypLE Express 1X (Gibco) and resuspended in differentiation medium. Each wrench-shaped pattern has an area of 4.3mm^2^. To achieve a density of 10^5^ cells per mm^2^ of pattern, 2.6×10^6^ cells are needed for 6 wrench-shaped patterns. 500µL of the cell suspension containing 2.6×10^6^ cells were added to each well. The wells on top of the patterns were then immediately positioned in between two strong magnets (110.6 × 89 × 19.6 mm, remanence Br = 1.35 T, Ref. Q-111-89-20-E, Supermagnete), spaced by 3.6 cm to minimize the gradient between them. This system, presented in Supplementary Figure 1C, was left for 3 hours in this configuration to allow the magnetic bioprinting process to occur.

### Chip Fabrication

Chips were fabricated by casting transparent Ecoflex (00-31 Near Clear, Smooth-on) in 3D-printed molds (Supplementary Fig. 4), into which needles were previously inserted in dedicated holes. After curing for 3 hours at 75°C, the needles were removed, and the chips were unmolded. Nickel-plated steel needles (0.6 mm wide, Bohin) were coated three times with transparent nail polish to prevent rusting in cell culture medium and then dried overnight at 120°C to remove any traces of solvent. These needles were placed into the chips’ pre-made holes so that two needles in each compartment faced each other at a distance of 2 mm. Once inserted, the needles were bonded to the chip using Loctite SI 5398 adhesive in the holes designated for stretching clamps. The needles were then coated with polydopamine by plasma-treating the chips and then filling the compartments with 5 mg/mL dopamine hydrochloride (Thermo Scientific) in Tris-HCL buffer (pH=8.5). After 3 hours, the compartments were rinsed with PBS. The activated surface of the needles was then coated with 0.2 mg/mL collagen I (Corning, rat tail) in PBS for C2C12 cells or Matrigel for M-hiPSC for 2 hours at room temperature, while the magnetic bioprinting process was conducted in parallel. Just before removing the substrate from the magnets, the chips were rinsed with PBS and filled with warm differentiation medium.

### Magnetic Trapping of a wrench shaped tissue

A row of 8 6×6mm N48 Neodymium magnets (Supermagnete, S-06-06-N) was placed on each side of the chip in 3D-printed slots, with one magnet aligned per needle. All magnets were oriented identically so that adjacent magnets repelled each other while the pair of magnets magnetizing needles facing each other attracted one another, creating magnetic field lines that extended from one needle tip to the other. After removing the substrate from the magnets, the six tissues within a well would either detach spontaneously or could be gently detached by applying a flow with a micropipette. The cohesive tissues were then aspirated using a 5 mL pipette and released in one of the chip’s compartments, in between the two needles where the tissue would clip onto. After incubating the tissue overnight, the magnets were removed for the rest of the maturation period.

### Stretching

Once the tissue had adhered to the needles overnight, the samples could be stretched. A pair of 3D-printed clamps was used to secure the chip onto a translation stage (Thorlabs). Stretching the chip increased the distance between the two needles, thereby stretching the tissue. Stretching the chip by 2 mm resulted in an approximate 1 mm increase in the distance between the needle tips, equivalent to 50% stretching. Stretching by 4 mm led to a 2 mm increase between the needle tips, corresponding to 100% stretching.

### Tissue Fixation and Staining

Culture medium was removed from the chips and replaced with 4% Paraformaldehyde (v/v) in PBS. Samples were incubated for two hours at room temperature while they were still attached to the needles, and, when applicable, in a stretched position. This allowed to preserve the exact configuration of the samples prior to fixation, and upon detachment from the needles using tweezers, their original shape was maintained. After rinsing the samples three times with PBS, they were incubated for 1h in blocking solution with 4% w/w BSA (Sigma) and 0.5% v/v Triton X-100 (Fisher) in PBS. Following another PBS rinse, the samples were incubated overnight at 4°C in blocking solution with 1:1000 Phalloidin AlexaFluor 555 (Thermofisher) and 1:300 DAPI (Invitrogen). For further immunostaining, samples were immersed in a saturation solution containing 1% (w/w) BSA and 0.3% (v/v) Triton X-100 in PBS, for one hour at room temperature. After rinsing with PBS, samples were incubated overnight in saturation solution with the desired primary antibodies: 1:200 Desmin (goat, R&D, AF3844), 1:100 Pax7 (mouse, DHSB, Pax7-c), 1:100 MF20 (mouse, DHSB, MF20-c), 1:200 Myogenin (mouse, DHSB, F5D), or 1:500 Alpha-Actinin (mouse, Sigma, A7811-.2ML). After washing with PBS, samples were incubated overnight in saturation solution with secondary antibodies, 1:750 donkey anti-goat 647 (Thermofisher Scientific, A32849) and 1:750 donkey anti-mouse 488 (Thermofisher Scientific, A-21202), and then washed with PBS. In the case of samples stained with EdU to detect proliferative cells, they were first incubated with 10µM EdU in culture medium for 2 hours before fixation. Once fixed, EdU incorporation was revealed using the Click-iT™ reaction cocktail from the Click-iT™ EdU Cell Proliferation Kit (Invitrogen). Afterward, samples were labeled with Hoechst dye and then incubated with antibodies, as previously described.

### Live/Dead Assay

The impact of the magnetic bioprinting and trapping over time on cell viability was assessed with LIVE/DEAD cell imaging kit (488/570, Invitrogen). Tissues that had just been patterned were transferred to a well of a 48-well plate filled with the reagents. For tissues suspended between the needles, the cell culture medium was replaced by the reagents. Tissues were incubated for 30 minutes and then imaged via confocal microscopy. Non-trapped tissues were first transferred to a fluorodish for imaging, while trapped tissues were imaged in situ within the chip, through the Ecoflex near-clear bottom layer. Live cells were visualized in green using a 488 nm excitation wavelength and dead cells were visualized in red using a 552 nm excitation wavelength.

### Microscopy

Fixed samples were positioned flat onto a fluorodish using a micropipette and tweezers. They were then imaged with a 25x water immersion objective on a Leica DMi8 inverted confocal microscope. Live microscopy was conducted within the incubator using a CytoSmart Lux 3 FL system. For imaging samples or systems from above, a DinoEye camera adapted to a Stemi 508 Greenough Stereo Microscope was used. Calcium imaging was performed using a Spinning Disk microscope, which allows for image acquisition at 100ms intervals.

### Gene expression analysis

Tissues were detached from the needles using tweezers. The central fiber of the tissue was cut and flash-frozen dry at −80°C. Total RNA was isolated using the RNeasy Micro Kit (Qiagen) and cDNA was synthesized using SuperScript III (Invitrogen) according to the manufacturer’s instructions. Quantitative real-time PCR was performed in triplicate using a QuantStudio 12K Flex RT-PCR system (Applied Biosystems) with the Luminaris HiGreen qPCR Master Mix (Thermo Fisher Scientific). Primer sequences are listed in Supplementary Table 2. Experiments were performed with at least three replicates per condition and expression levels were normalized to 18S. Relative expression compared to 2D gene expression levels at day 0 was determined by calculating the 2^−ΔΔCt^.

### Magnetic field simulations

Simulations to determine the magnetic field and gradient around the NiFe patterns during magnetic bioprinting and the needles during the magnetic trapping process were performed using COMSOL Multiphysics (Magnetic Field No Currents module, Licence number 6464550).

### Designing and printing 3D parts

The molds, chips and various 3D-printed parts used in our set up were designed using Autodesk® Inventor® (Supplementary Fig. 4 and 16). These designs were then 3D printed using a Digitalwax 028J Plus® 3D printer (DWS).

### Fabrication of fluorescent polyacrylamide beads

Beads were fabricated by adapting previously established protocols^48,67^. Briefly, an aqueous solution containing 0.63% w/w Acrylamide (Sigma A4058), 1.13% w/w Bis-acrylamide molar ratio 19:1 (Sigma A9926), 10 mM of Tris buffer pH7.5 and 0.15% v/v ammonium persulfate, and an HFE 7500 fluorinated oil solution with 0.4% v/v TEMED (Sigma), 1% w/v FluoroSurf surfactant (Emulseo) and 0.1% w/v N-hydrosuccimide ester (Santa Cruz Biotechnology Inc.) was prepared, vortexed and filtered. Using a microfluidic chip similar to the one used in the original study^67^, the acrylamide solution was flowed through 20 x 35 µm channels at 300 µL/h. This flow was disrupted by an 800 µL/h flow of the oil solution, resulting in the formation of an emulsion of acrylamide droplets in oil. This emulsion was then collected in an Eppendorf tube with mineral oil and incubated at 60°C for 16 hours. The following day, the bottom and top layers of HFE and mineral oils were removed. The emulsion was then washed at least 3 times with 20% Perfluoro-octan-1-ol in HFE oil, at least two times with Hexane with 1% (w/v) Span-80, and finally with three washes with PBS. This process resulted in the formation of spherical deformable polyacrylamide beads with an average diameter of 42±5 µm. The beads were then incubated overnight in a rotating Eppendorf tube with 0.1 mg/mL FITC-labeled poly-L-lysine (Sigma) in sterile 50 mM HEPES (pH 8.2). Finally, the beads were washed with PBS. Just before bioprinting the tissues, the beads were mixed into the cell suspension at a concentration of one bead per 500 cells. Although the beads were not magnetic, they became easily embedded in the tissues during the bioprinting process.

### Calcium imaging and Acetylcholine assays

Tissues were incubated with Fluo-4, AM, a cell-permeant dye (Thermofisher Scientific), in HBSS buffer, for one hour. After incubation, tissues were rinsed with HBSS buffer and differentiation medium, and then transferred in 240 µL of differentiation medium in a 96-well plate for imaging. For tissues trapped between needles, they were carefully detached from the needles and allowed to recover and stabilize for 5 minutes before imaging. Samples were imaged using a Spinning-Disk microscope with a 488nm laser, capturing images every 100 ms for a total duration of 3 minutes. Approximately one minute into the imaging session, 60 µL of a 5 mM acetylcholine (ACh) solution was added to the well, achieving a final concentration of 1 mM ACh without causing excessive movement of the tissue.

### Image analysis

Image analysis was performed using Fiji. The orientation distributions of actin were analyzed with the Directionality plugin, normalized, and plotted as probability distribution curves, where the proportion of filaments between two angles corresponds to the area under the curve between those angles. 3D segmentation to measure the numbers of nuclei per cell and their lengths was performed manually using the ROI manager tool. Bead deformation was quantified using a simple model by fitting ellipses to the bead projections. Aspect ratio and orientation of the ellipses were extracted using the Analyze Particles tool. Frequencies of calcium fluctuations were measured in five cells exhibiting fluctuations from each sample without ACh, and within 2min following ACh injection. Cells that did not exhibit fluctuations were excluded from the analysis. The relative increase in calcium signal intensity was calculated between the intensity just before ACh addition and the maximal intensity after addition, normalized by the background intensity. Intensity increase was assessed over a large area of the tissue, rather than focusing on individual cells. The numbers of Myogenin- and Pax7-positive cells were counted manually using the Cell Counter plugin.

### Statistical analysis

Data were processed using Microsoft Excel and represented using GraphPad Prism 8 (GraphPad Software, La Jolla, California). Values were averaged from at least three independent replicates. Errors bars represent ±standard deviation. Significant differences between two conditions were evaluated using two-tailed unpaired Student’s t-tests with a 95% confidence interval, except for frequencies of calcium fluctuations which were evaluated with two-tailed unpaired Welch’s t-tests which do not assume equal standard deviations. Statistical significance is indicated with * for P value < 0.05, ** for P < 0.01, *** for P < 0.001 and **** for P < 0.0001.

## Acknowledgements

This work was funded by the French Agence Nationale de la Recherche (ANR) under grant ANR-19-CE09-0029. This work has also received the support of “Institut Pierre-Gilles de Gennes” (laboratoire d’excellence, Equipex, “Investissements d’avenir” program ANR-10-IDEX-0001-02 PSL and ANR-10-LABX-31-34). This work benefited from the technical contribution of the engineers of the joint service unit CNRS UAR 3750 at Institute Pierre Gilles de Gennes (IPGG) and the authors would like to thank the engineers of this unit. The authors also thank Morgan Gazzola from I-STEM for help with hiPS derived myoblasts and calcium imaging.

## Author contribution

N.D. and C.W. conceptualized the study. N.D., L.M., G.G., Si.D., C.W. performed experiments. St.D. and C.M. assisted in designing the experiments and proofread the manuscript. C.W. and St.D. supervised the study. N.D. and C.W. wrote the manuscript.

