## Supplementary information for "Magnetic printing and actuation of stretchable muscle tissue"

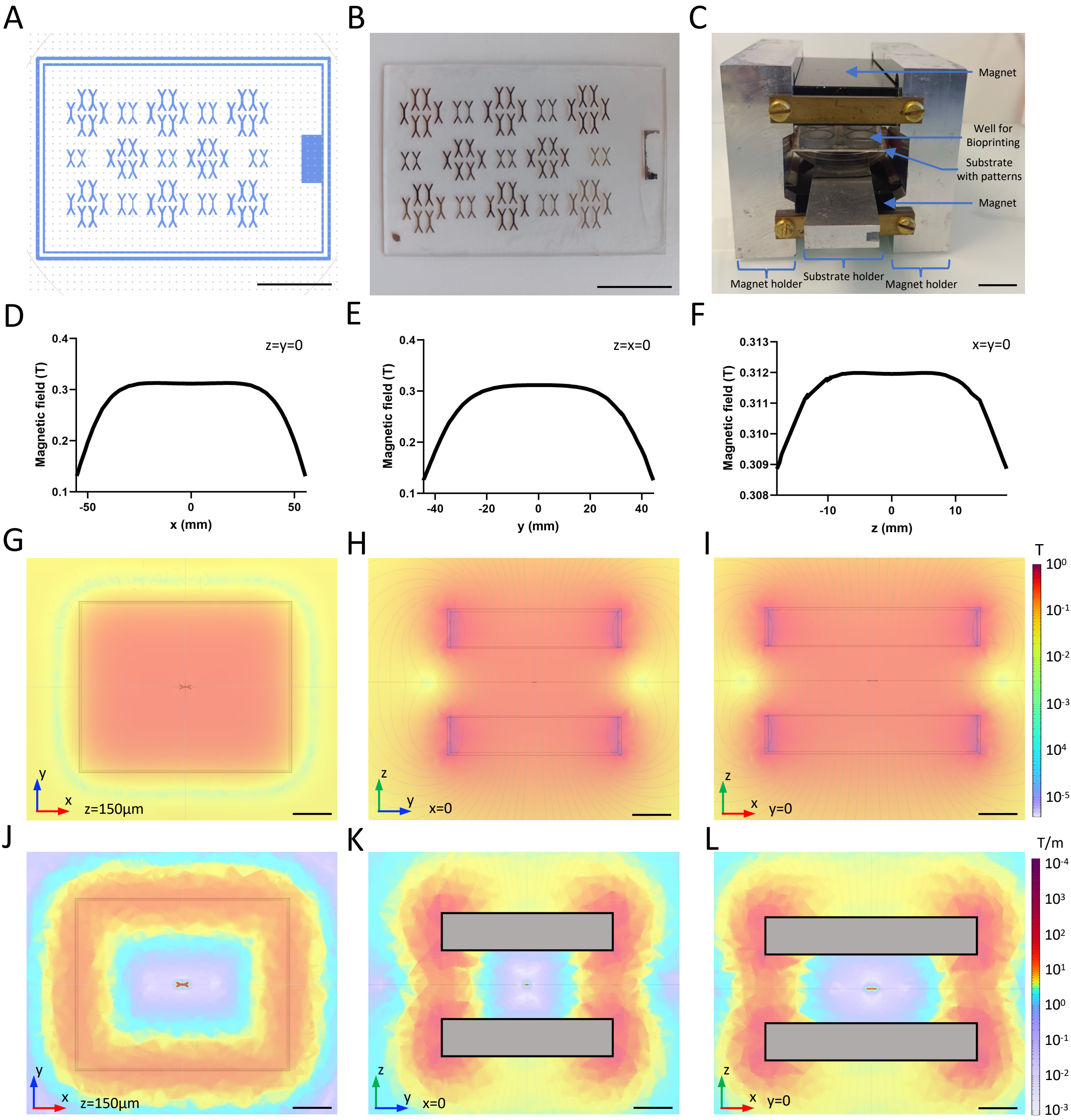


**Supplementary Figure 1 .** (A) Design of the photolithography mask on Clewin to generate (B) the substrate with the matching NiFe patterns after photolithography and NiFe electroplating. (C) Magnetic bioprinting set-up and the graphs of the simulated uniform magnetic field generated at the center the magnets without any patterns in it, plotted along the (D) x, (E) y, and (F) z axes. Heatmaps of the magnetic field (G, H, I) and gradient (J,K,L) with the field lines generated by the two strong patterns with a single NiFe pattern in the center, visualized in 3 different planes. (Scale bar=2cm)

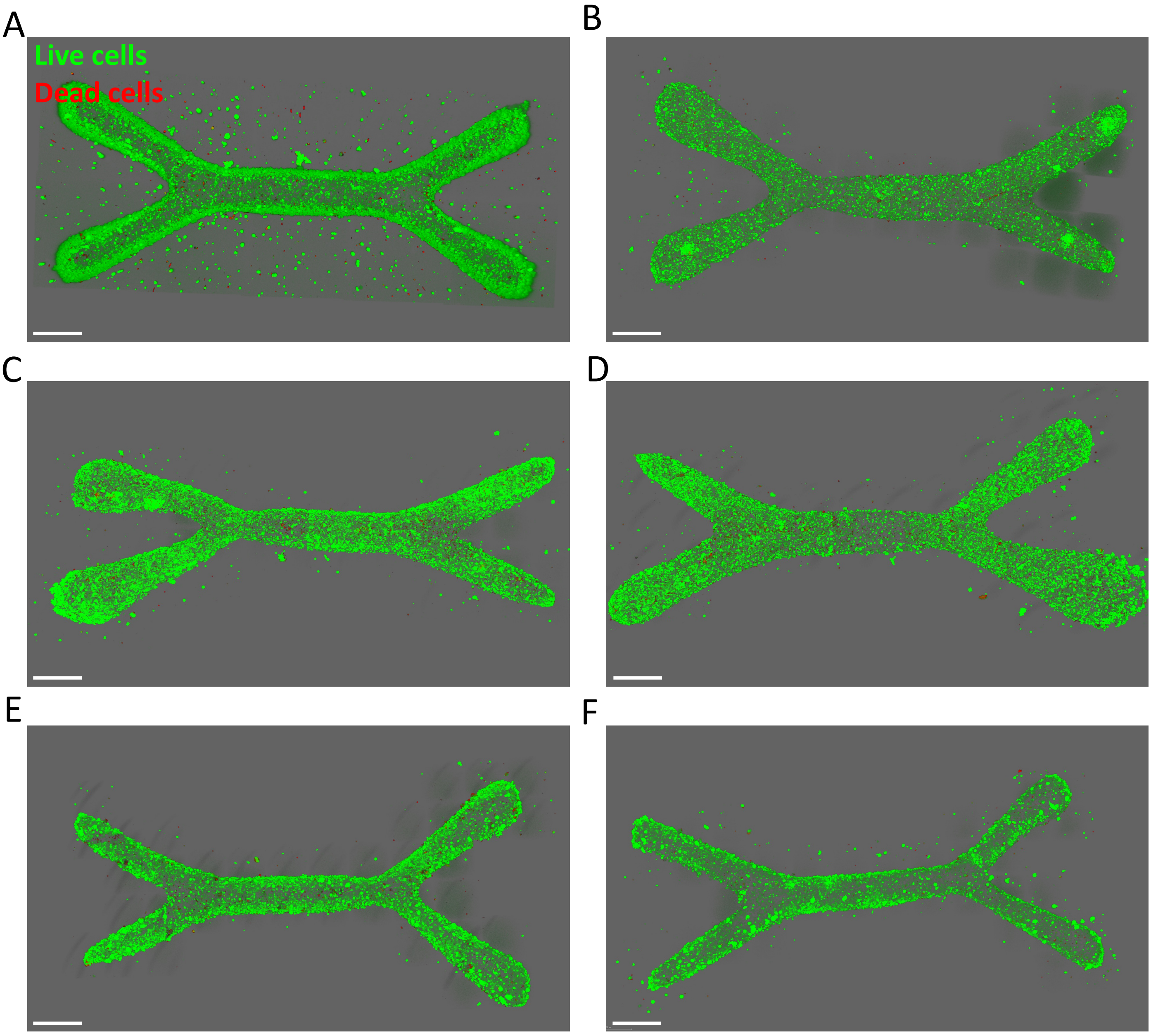


**Supplementary Figure 2.** (A-F) 3D reconstructions from confocal imaging of LIVE/DEAD assays performed on different tissues made from C2C12, just after they were magnetically bioprinted (Live cells in green, dead cells in red, scale bar = 500µm)

A

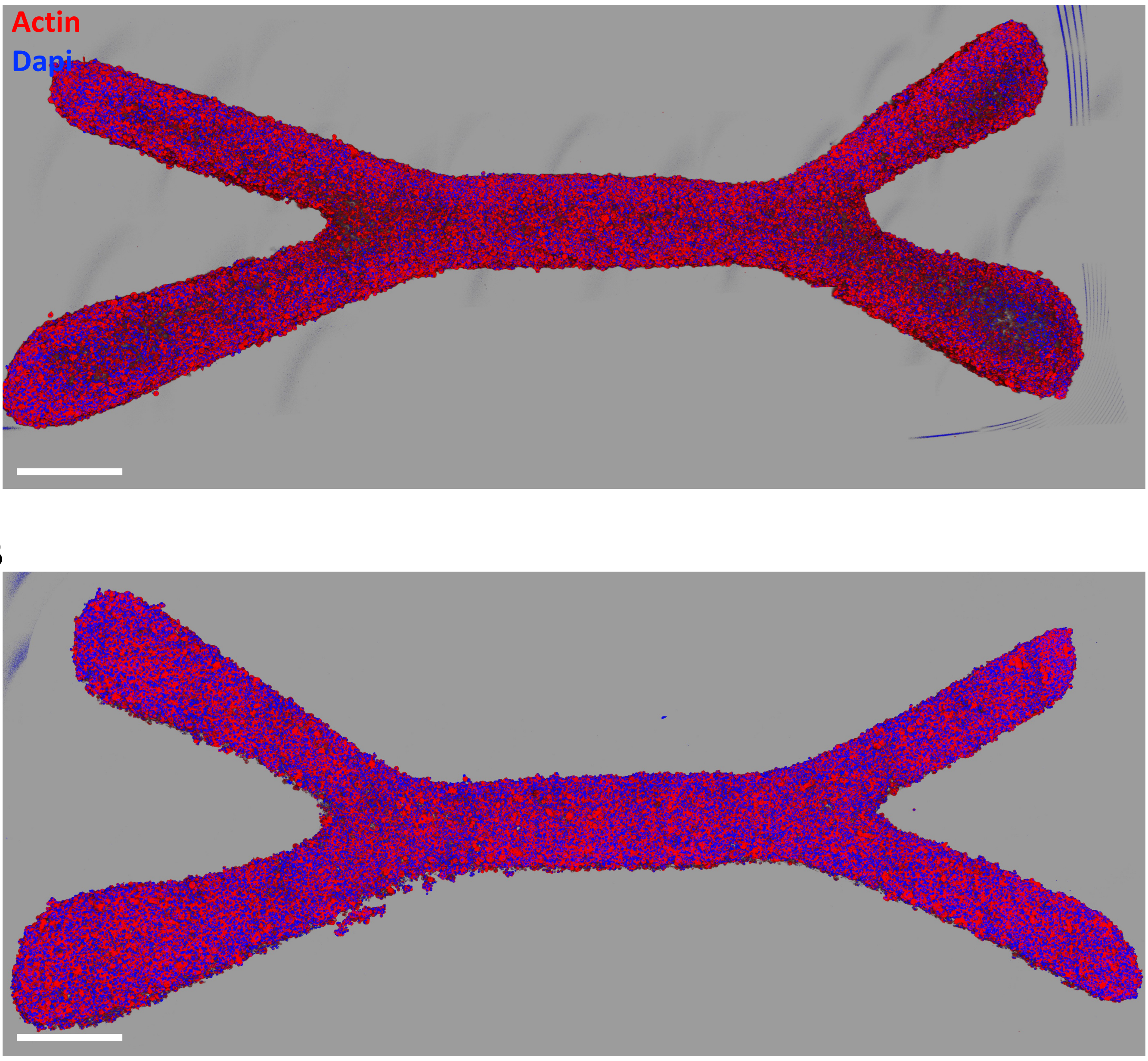


B

**Supplementary Figure 3.** (A,B) 3D reconstructions from confocal imaging of different tissues made from C2C12 cells, just after they were magnetically bioprinted (Actin in red, nuclei in blue, scale Bar=500µm)

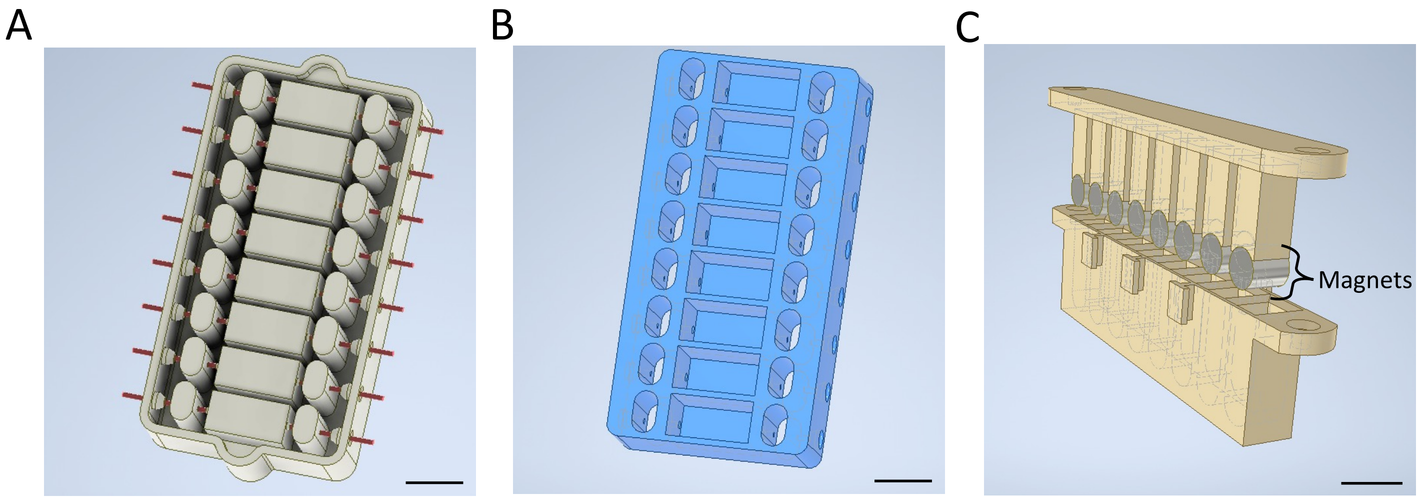


**Supplementary Figure 4 .** Designs of (A) the mold with needles inserted in it, (B) the resulting chip and (C) the opened-up system to hold the eight 6x6mm cylindrical magnets together, made with Autodesk Inventor (Scale bar=1cm)

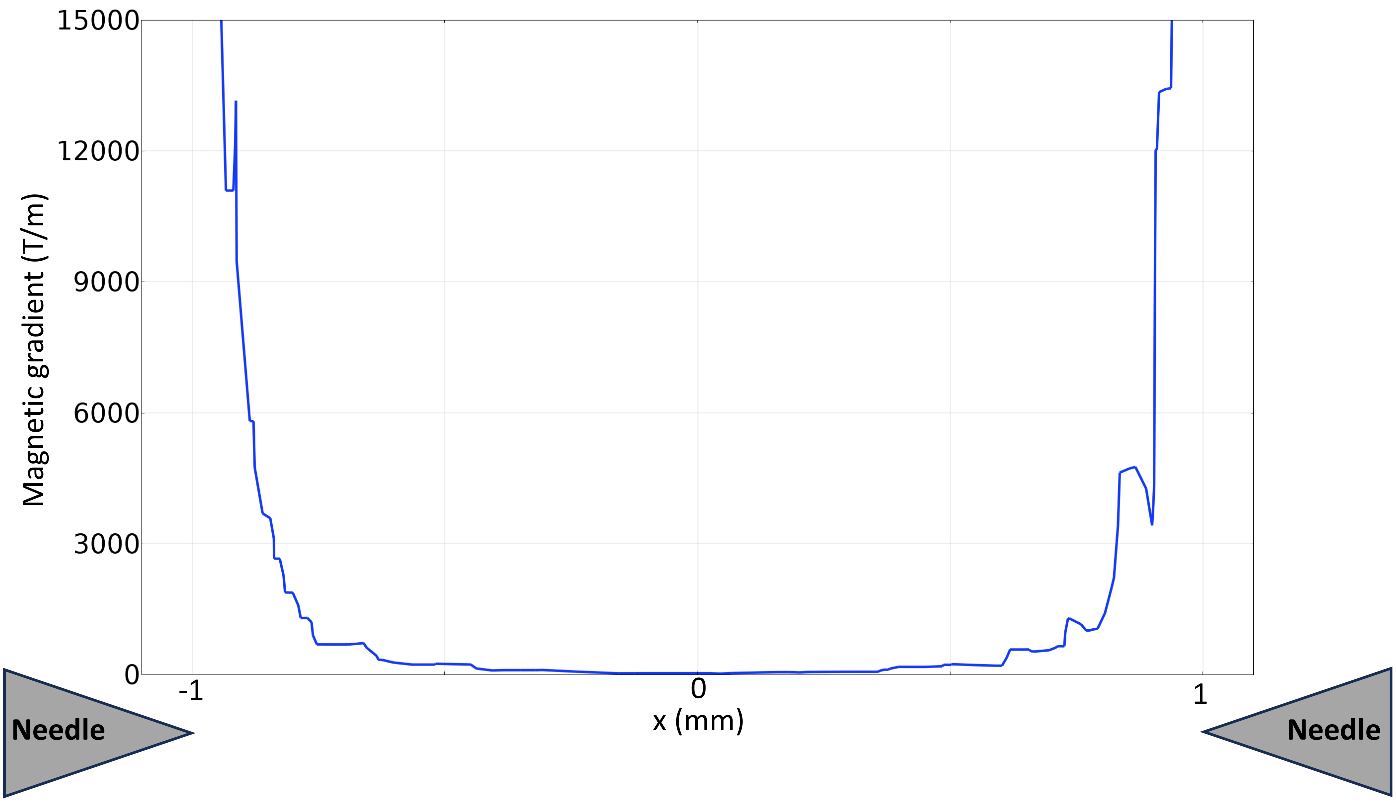


**Supplementary Figure 5 .** Plot of the magnetic gradient between two needles in the chip along their axis, generated from Comsol simulations

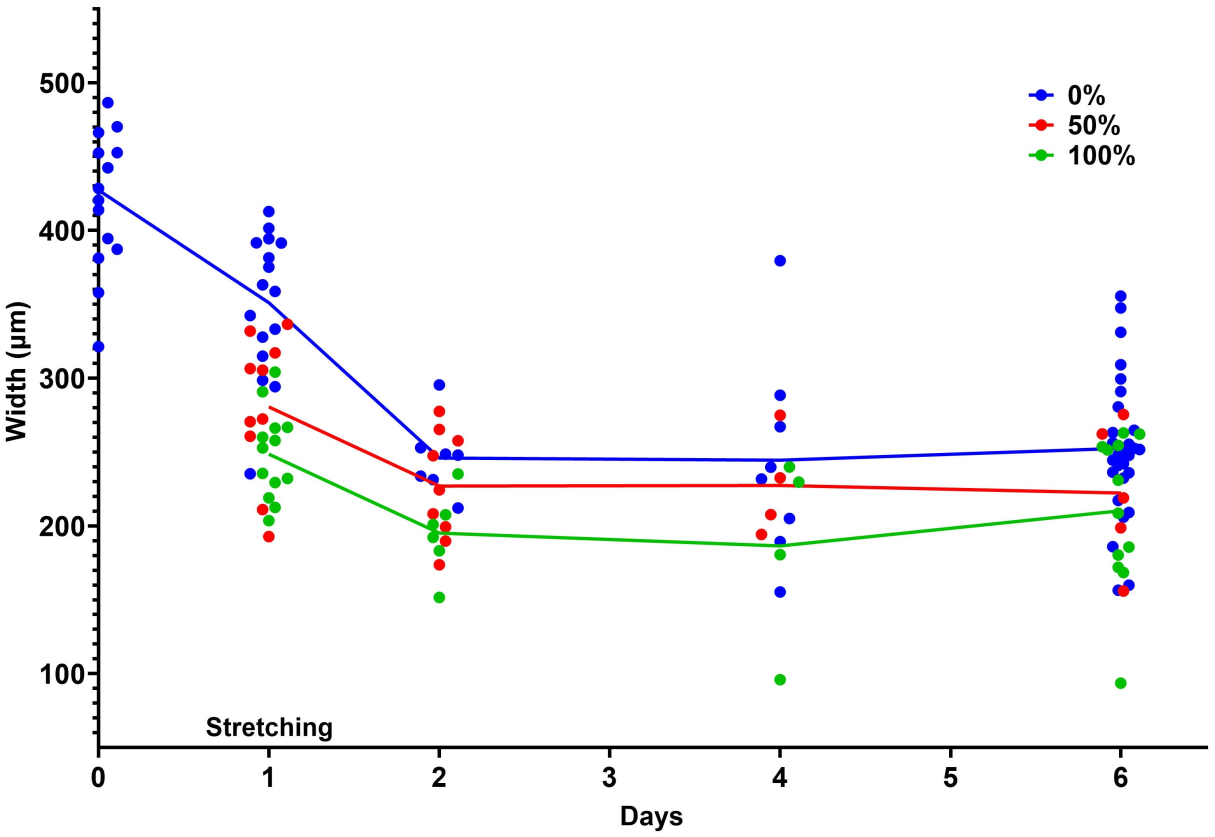


**Supplementary Figure 6.** Evolution of the widths of tissues made of C2C12 cells from day 0, after they were bioprinted, to day 6, whether they were stretched by 0%, 50% or 100% at day 1

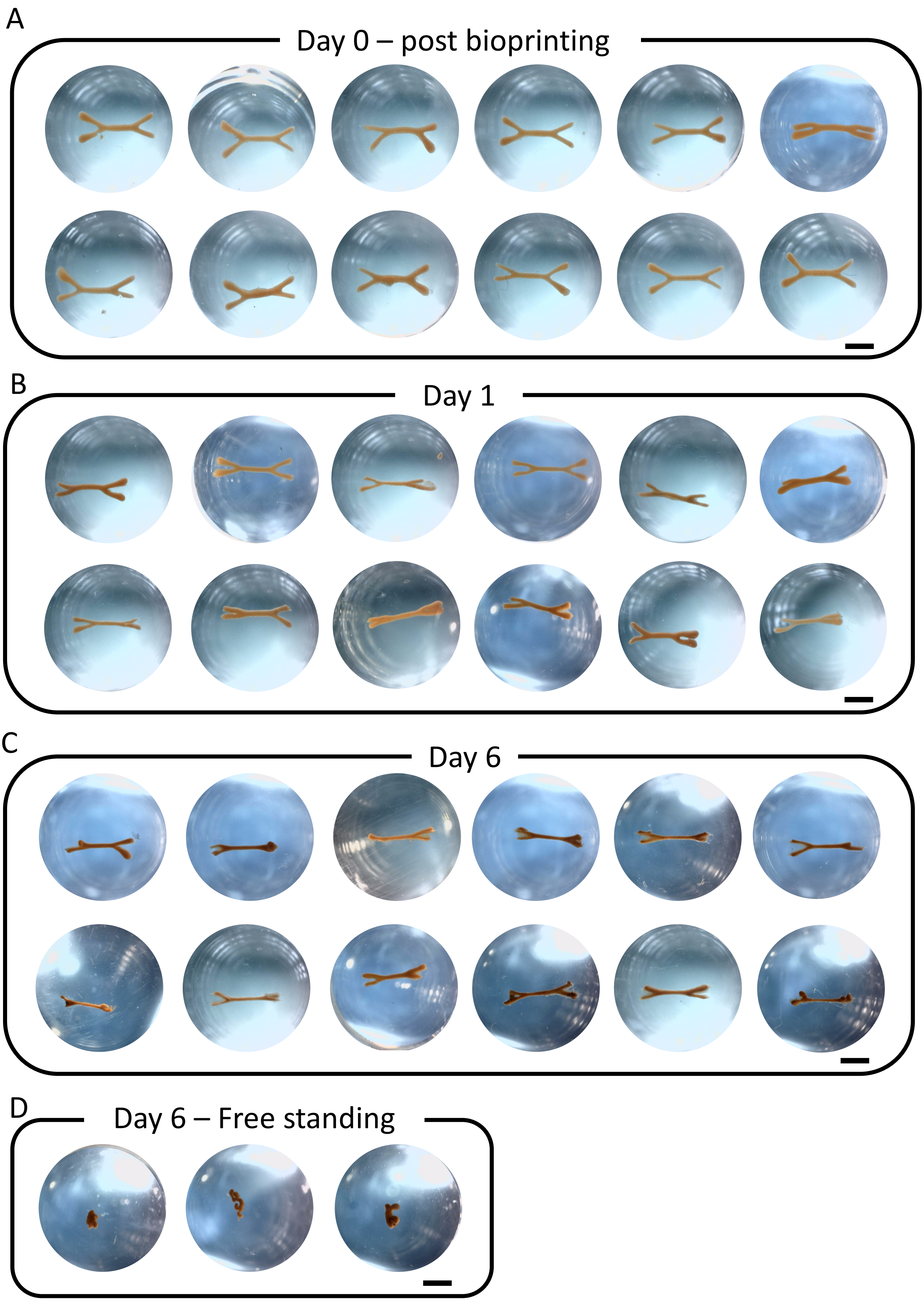


**Supplementary Figure 7.** Different tissues made of C2C12 cells that were fixed and imaged in PBS in the wells of the 48-well plate in which they were preserved. 3D issues that were fixed (A) at day 0 just after being bioprinted, and tissues cultured trapped on the needles fixed (B) at day 1 and (C) at day 6 . (D) Tissues that were cultured free standing in a non adherent well collected at day 6. Tissues that were trapped between needles were fixed still trapped in the chip, then removed from the needles by scrapping the clamps with tweezers and placed in PBS filled wells. They were imaged from the top using a magnifier and a dino camera. (Scale bar= 2mm)

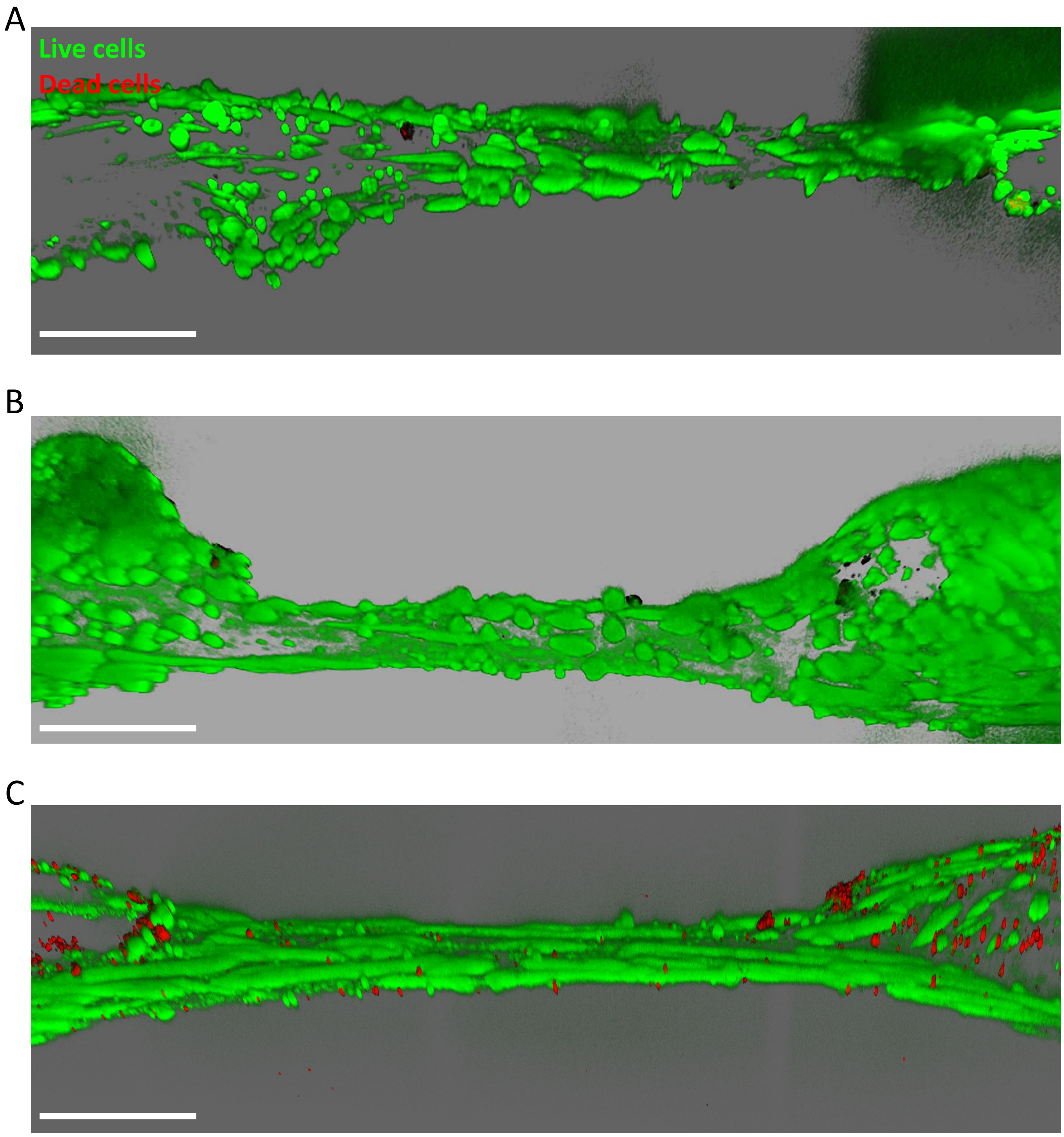


**Supplementary Figure 8.** (A-C) 3D reconstructions from confocal imaging of LIVE/DEAD assays performed on different non stretched 6 day-old tissues made from C2C12 cells, imaged in the chip still trapped on the needles (Live cells in green, dead cells in red, scale bar=500µm)

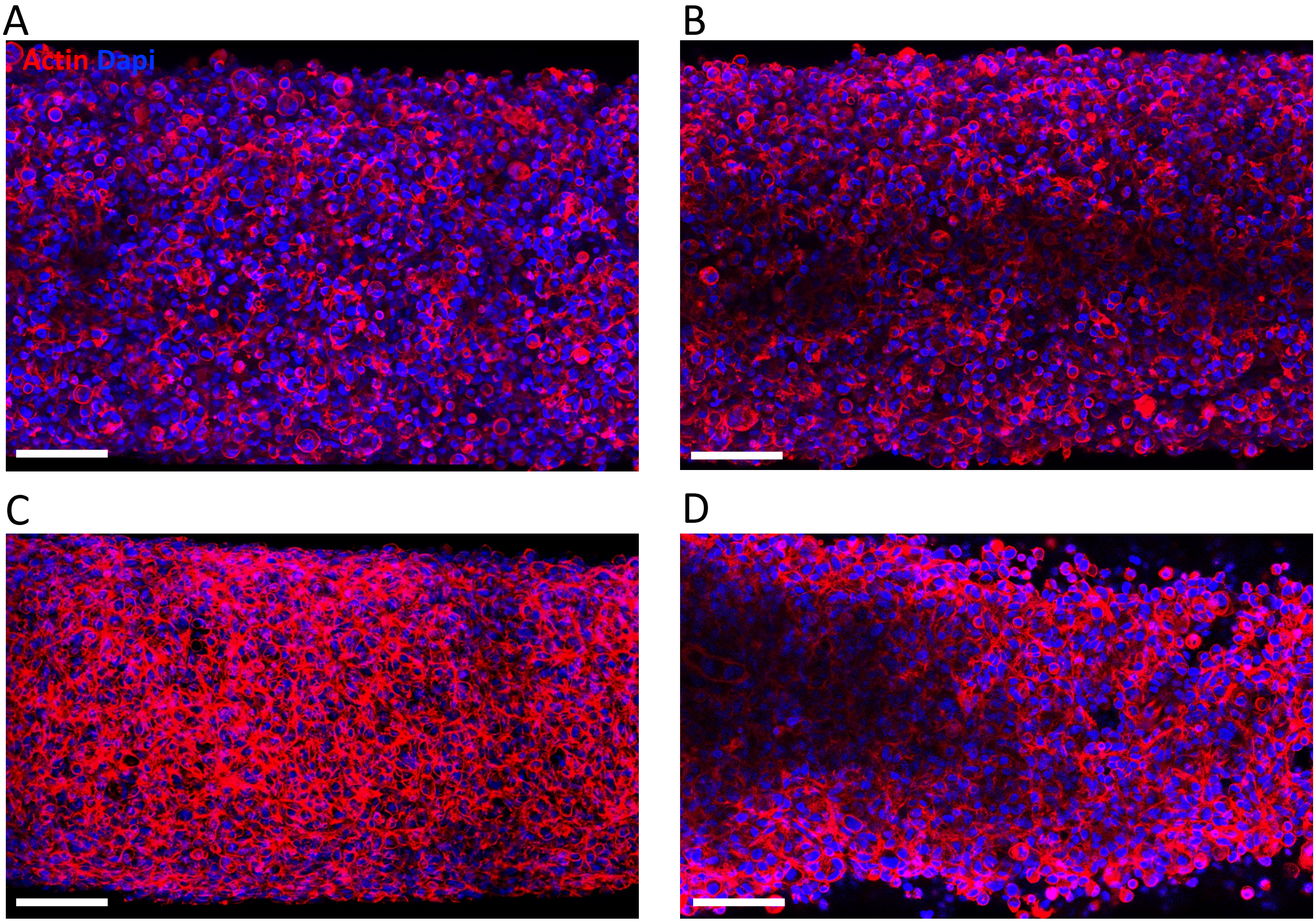


**Supplementary Figure 9.** Confocal imaging of different tissues made of C2C12 cells at day 0, just after bioprinting (Actin in red, nuclei in blue, scale bar = 100µm)

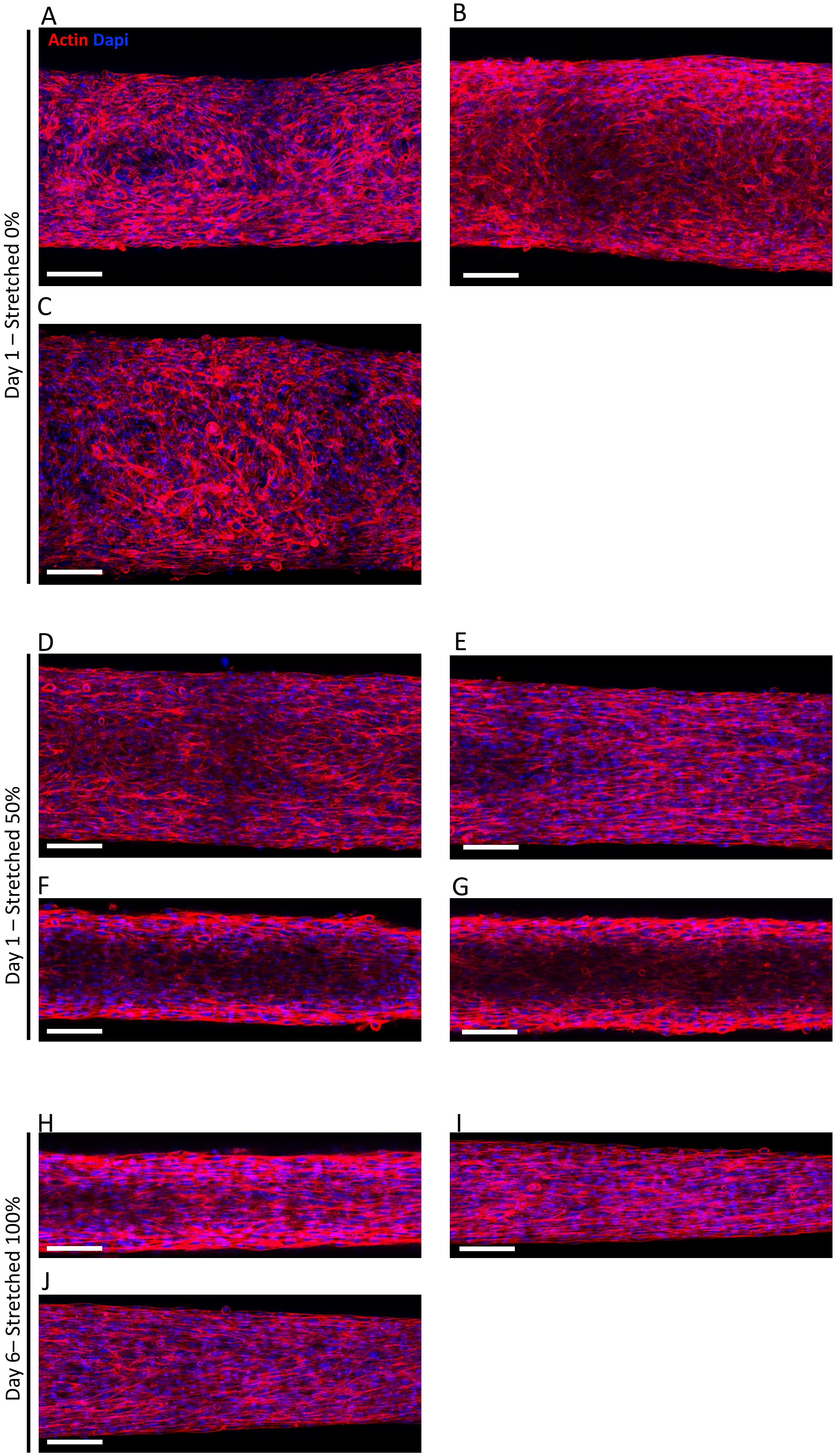


**Supplementary Figure 10.** Confocal imaging of different tissues made of C2C12 cells at day 1, which were stretched by (A-C) 0%, (D-G) 50% and (H-J) 100% (Actin in red, nuclei in blue, scale bar=100µm)

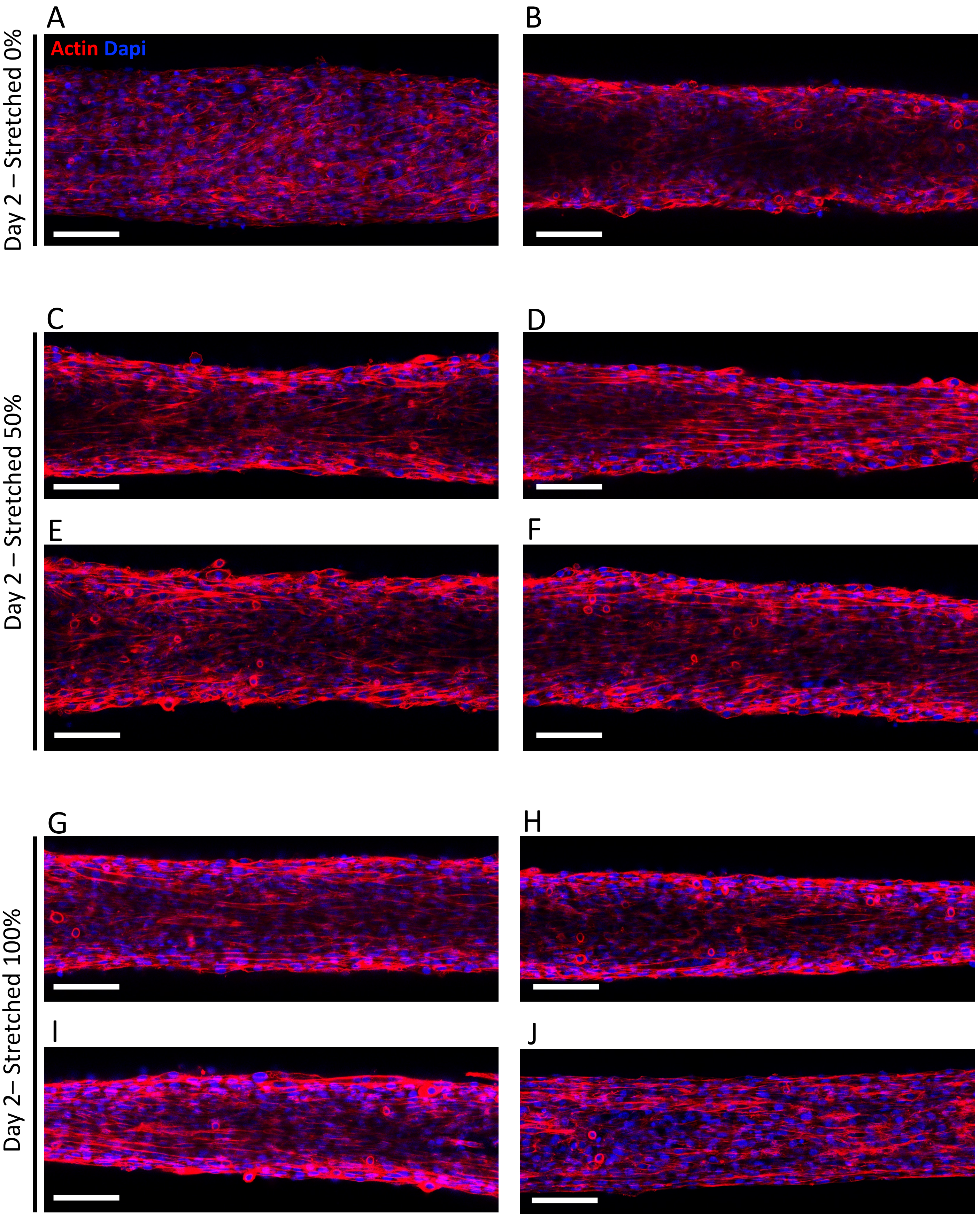


**Supplementary Figure 11.** Confocal imaging of different tissues made of C2C12 cells at day 2, which were stretched since day 1 by (A-B) 0%, (C-F) 50% and (G-J) 100% (Actin in red, nuclei in blue, scale bar = 100µm)

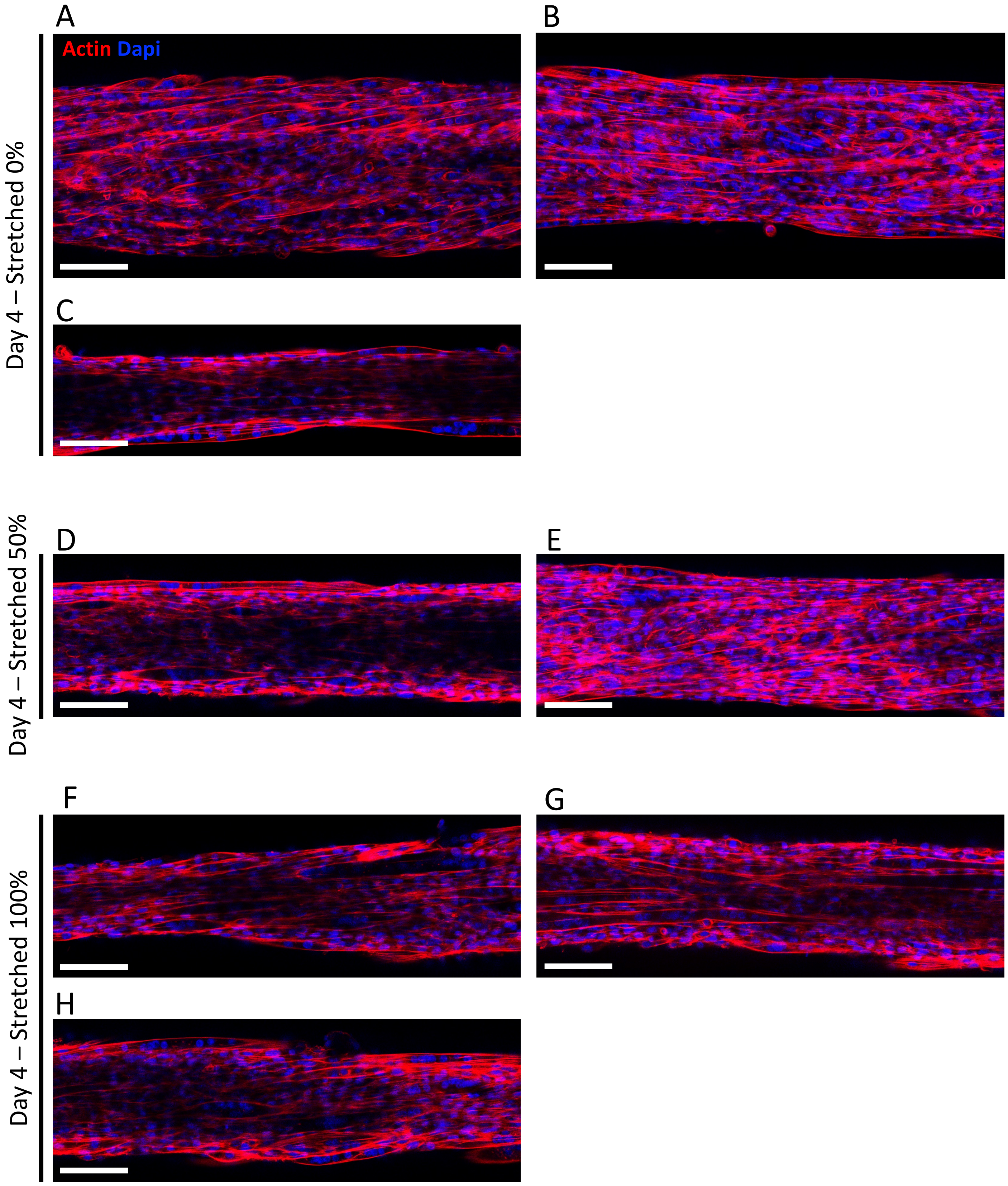


**Supplementary Figure 12.** Confocal imaging of different tissues made of C2C12 cells at day 4, which were stretched since day 1 by (A-C) 0%, (D-E) 50% and (F-H) 100% (Actin in red, nuclei in blue, scale bar = 100µm)

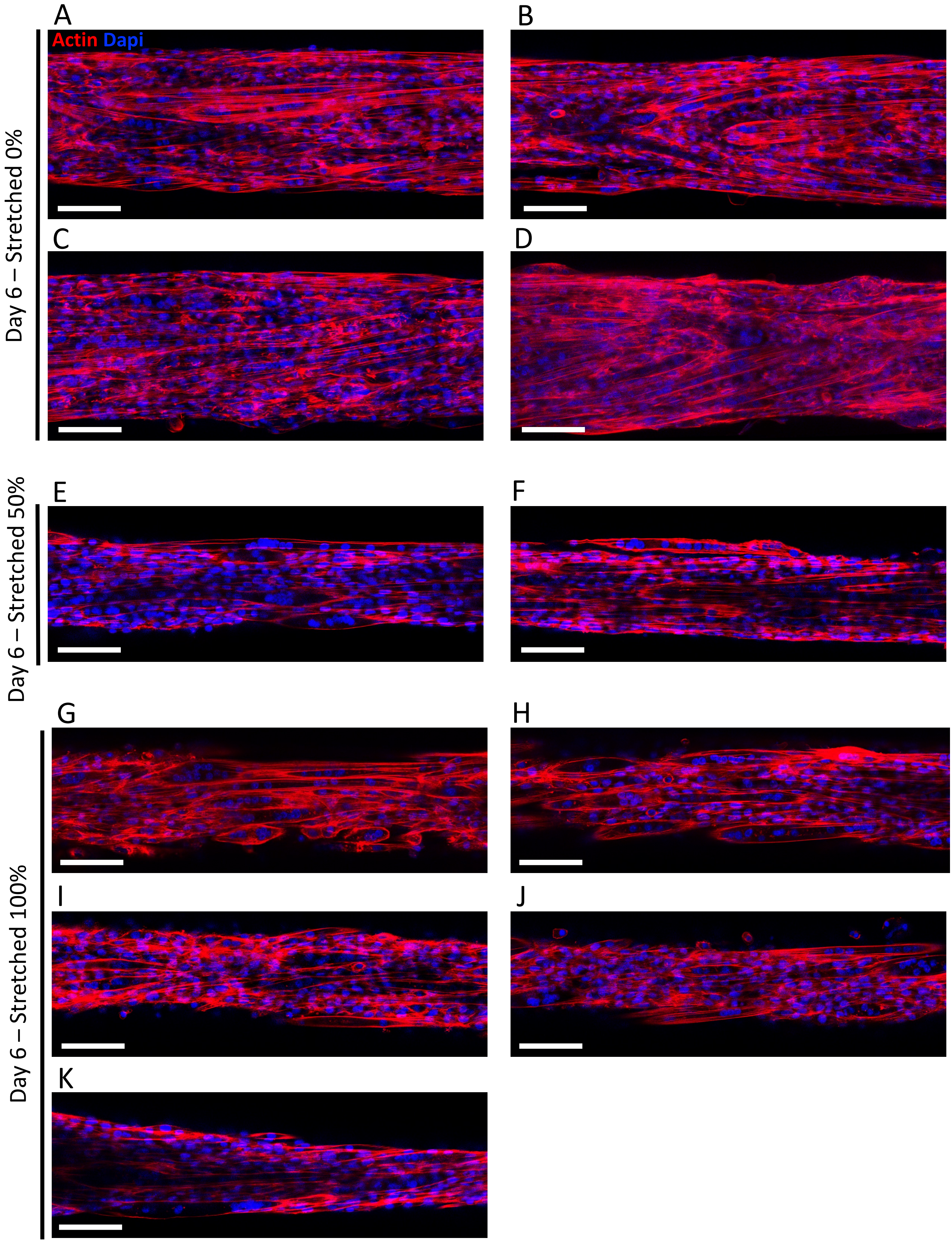


**Supplementary Figure 13.** Confocal imaging of different tissues made of C2C12 cells at day 6, which were stretched since day 1 by (A-D) 0%, (E-F) 50% and (G-K) 100% (Actin in red, nuclei in blue, scale bar=100µm)

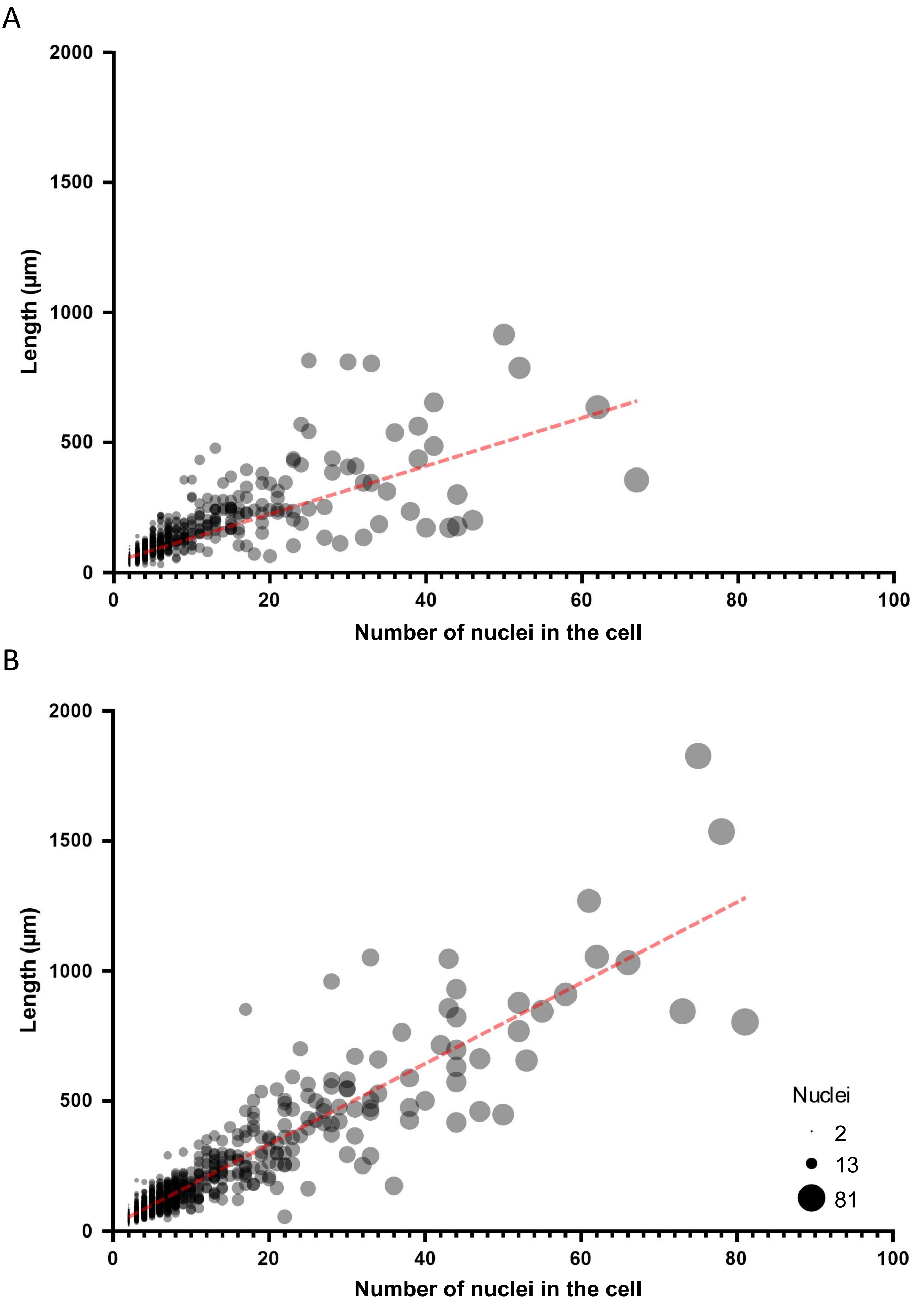


**Supplementary Figure 14 .** Scatter plots of the lengths of mulitnucleated cells as a function of the number of nuclei in the cells for 6 day old tissues that were stretched by (A) 0% (n=3) and (B) 100% since day 1 (n=3). The diameter of each point is proportionate to the number of nuclei in the cell. A trendline in red was plotted from a linear regression.

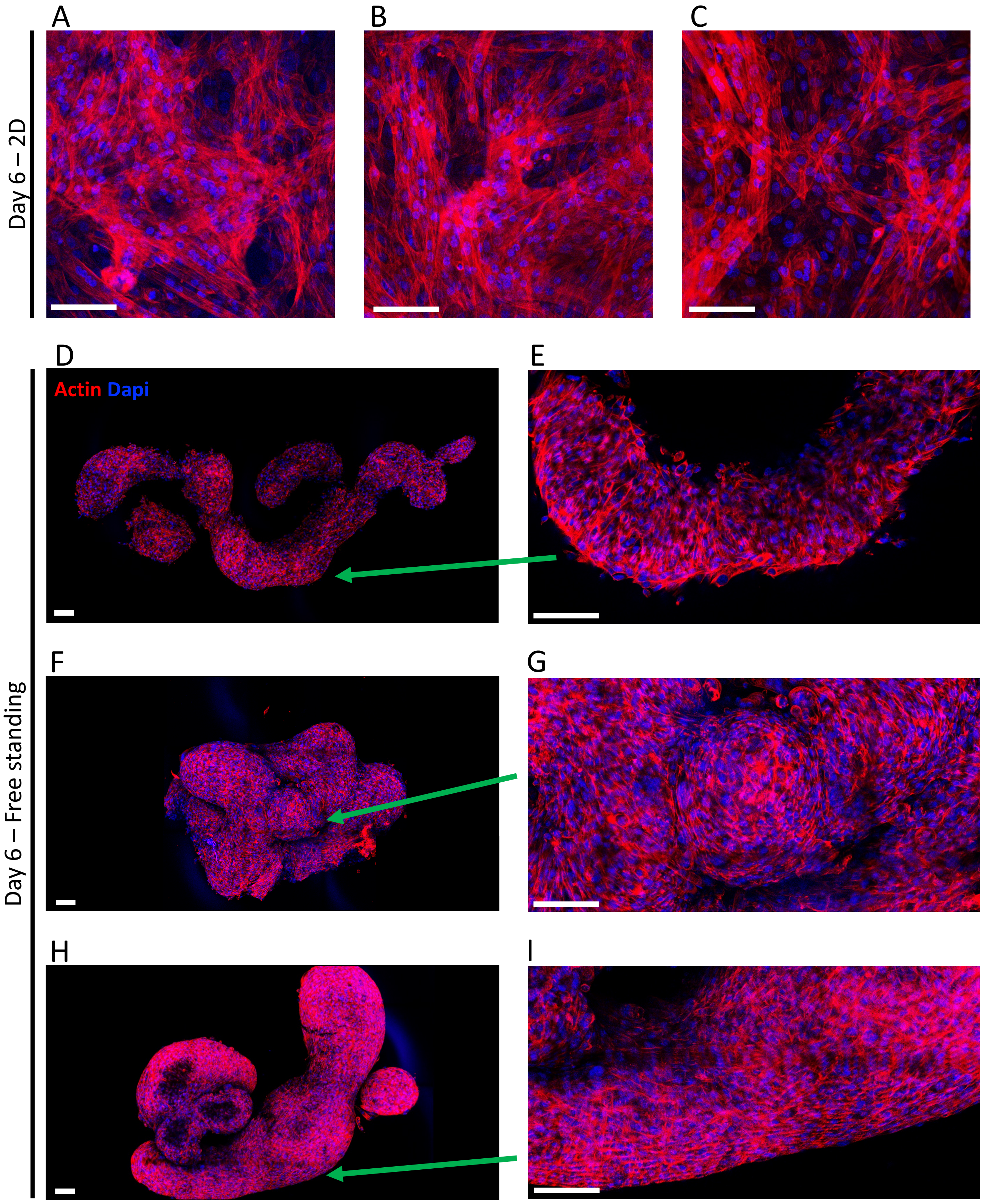


**Supplementary Figure 15.** Confocal imaging at day 6 of (A-C) C2C12 cells cultured in 2D and of (D-I) different 3D tissues which were not trapped between needles and instead were left free standing in non-adherent wells (Actin in red, nuclei in blue, scale bar = 100µm)

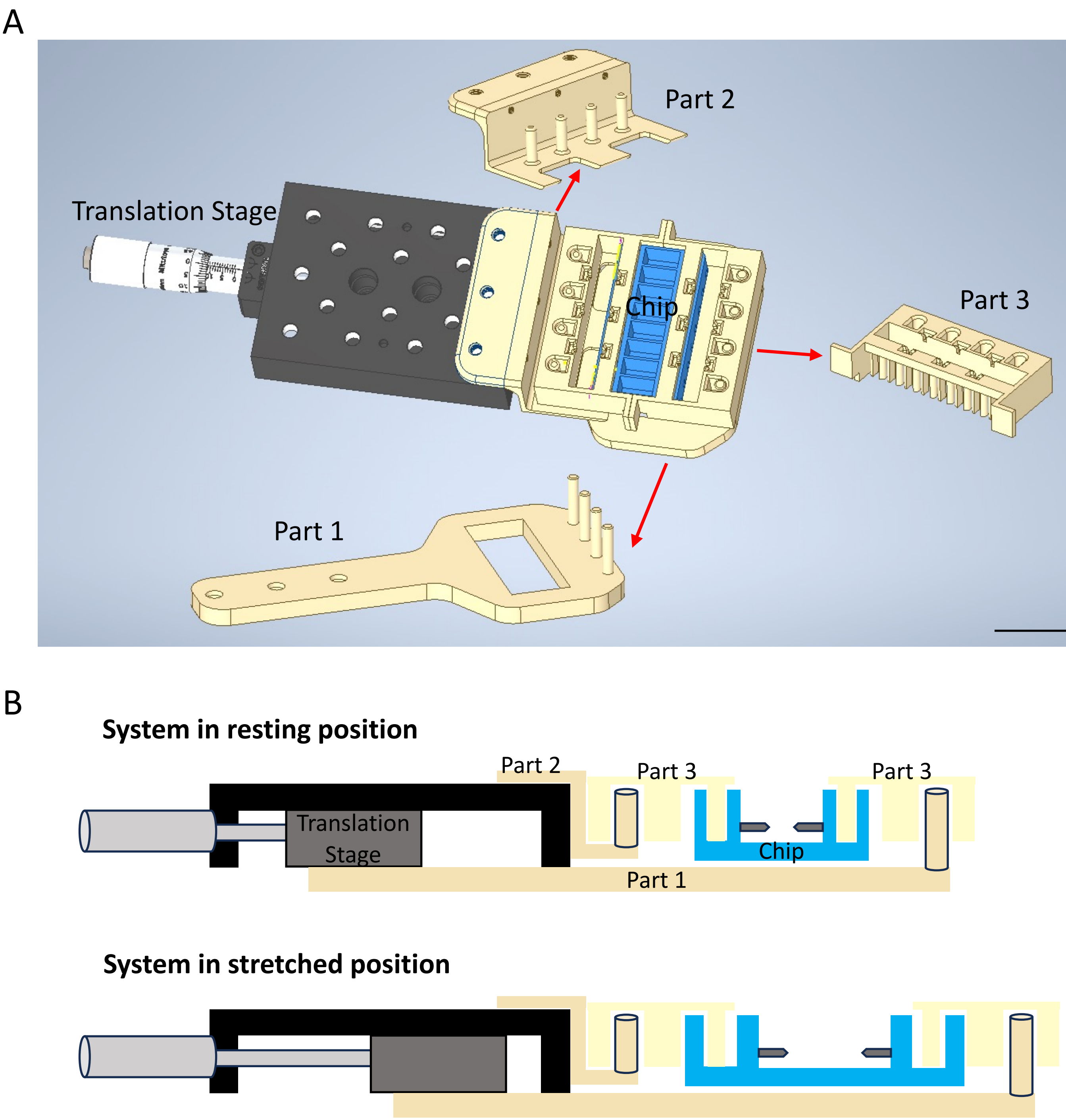


**Supplementary Figure 16 .** (A) Designs of the system stretching the chip and the individual 3D printed parts composing it, made with Autodesk Inventor (Scale bar=3cm), and (B) simplified shematics showing how these different parts are organized from a longitudinal cut when the system is in its resting position or stretched position.

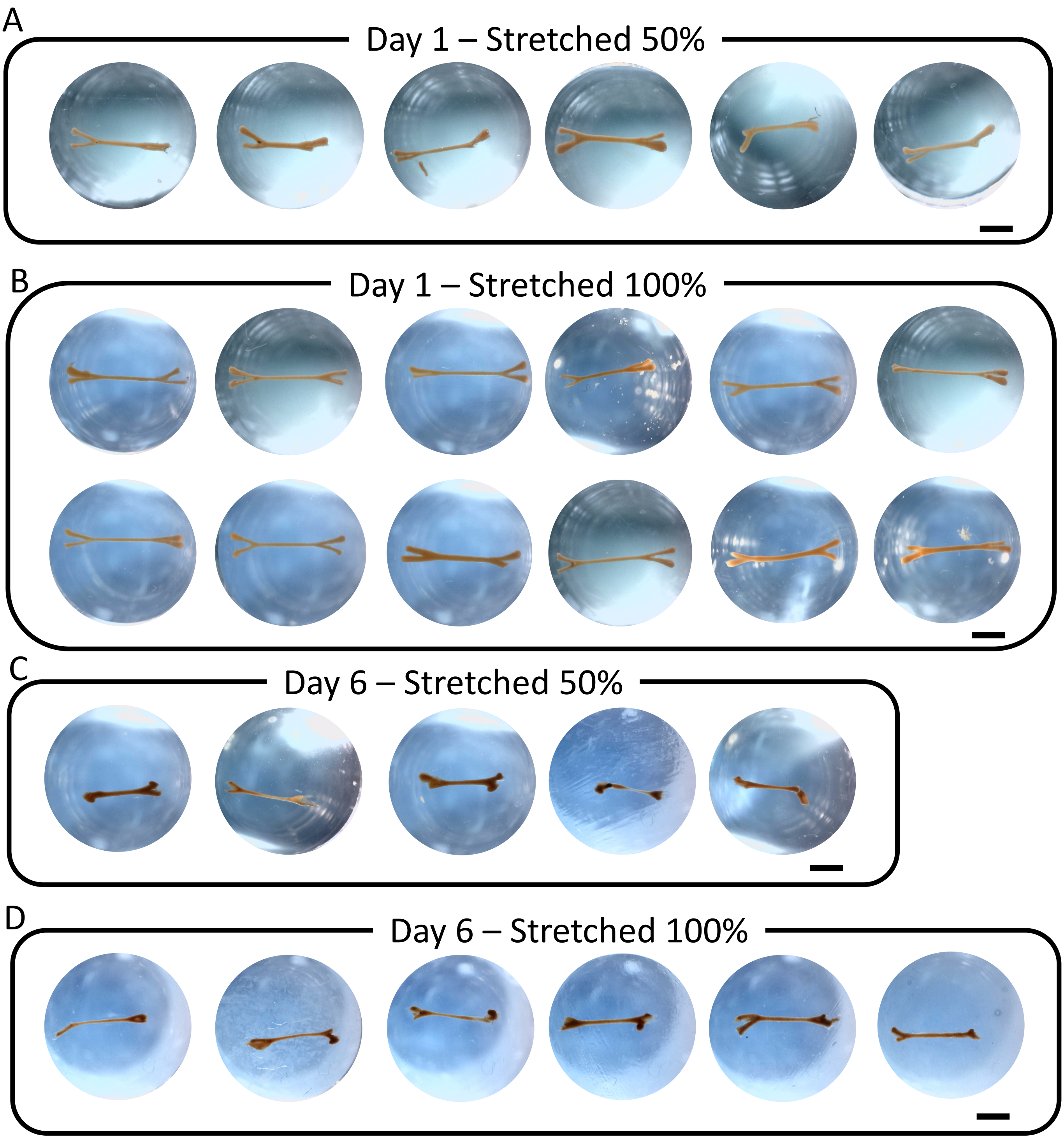


**Supplementary Figure 17.** Different tissues made of C2C12 cells that were fixed and imaged in PBS in the wells of the 48-well plate in which they were preserved. Tissues collected on day 1 just after they were stretched by (A) 50% or (B) 100% and on day 6 which were stretched since day 1 by (C) 50% or (D) 100%. Tissues that were trapped between needles or stretched were fixed still trapped or stretched in the chip, then removed from the needles by scrapping the clamps with tweezers and placed in PBS filled wells. They were imaged from the top using a magnifier and a dino camera. (Scale bar = 2mm)

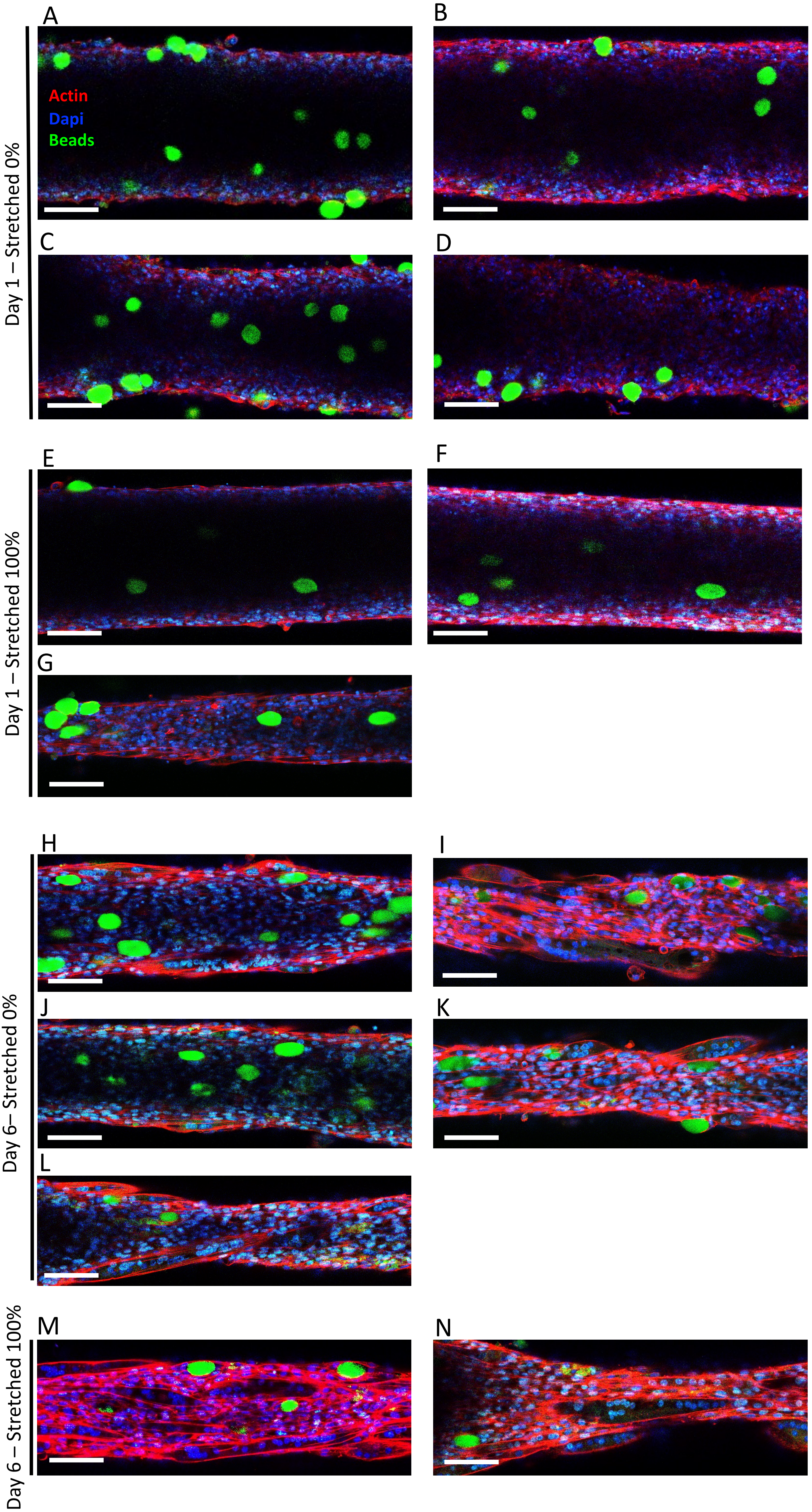


**Supplementary Figure 18.** Confocal imaging of different tissues made of C2C12 cells, in which polyacrylamide beads were embedded, at day 1 after they were stretched by (A-D) 0% or (E-G) 100%, and at day 6 which were stretched since day 1 by (H-L) 0% or (M-N) 100%. (Actin in red, dapi in blue, beads in green, scale bar = 100 µm)

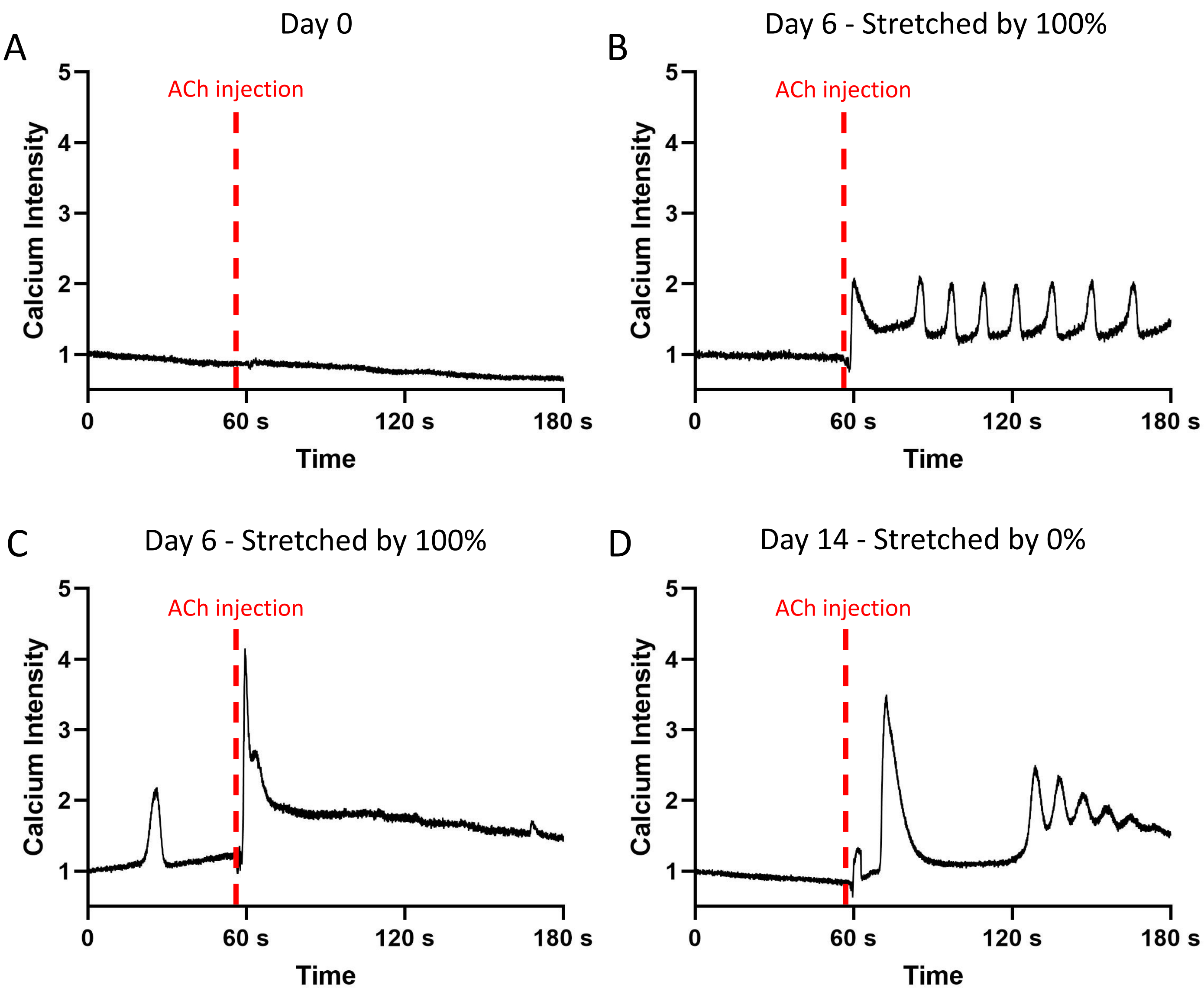


**Supplementary Figure 19.** Kymographs of the fluctuations of the calcium signal in (A) a C2C12 cell from a tissue at day 0 just after bioprinting, (B-C) in two different C2C12 cells from tissues at day 6 stretched by 100% since day 1, and (D) a C2C12 cell from a non-stretched tissue at day 14, treated with 1mM ACh after approximately 1min.

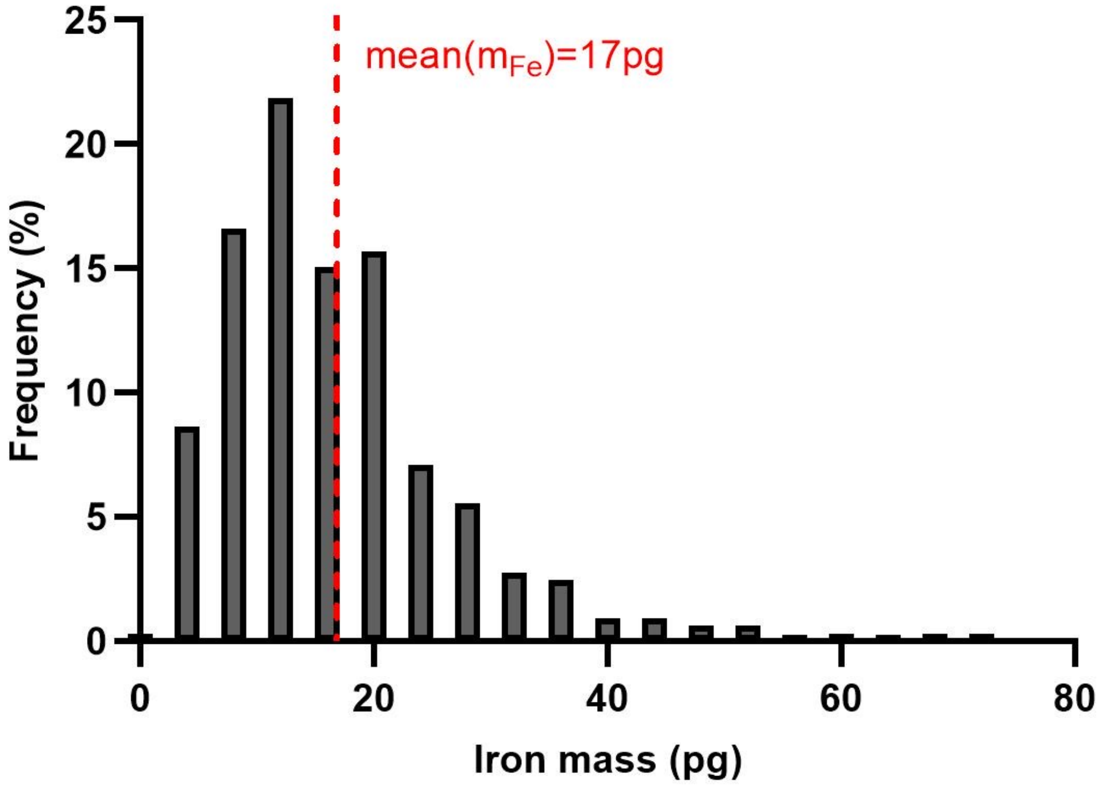


**Supplementary Figure 20 .** Histogram of the distribution of iron mass internalized per cell for M-ihPSC 1 collected just before patterning, after they were labeled with 2mM of iron and 2mM citrate in RPMI for 15mn once a day for the two previous days and with 0.2mM of iron in their proliferation medium overnight between these two days

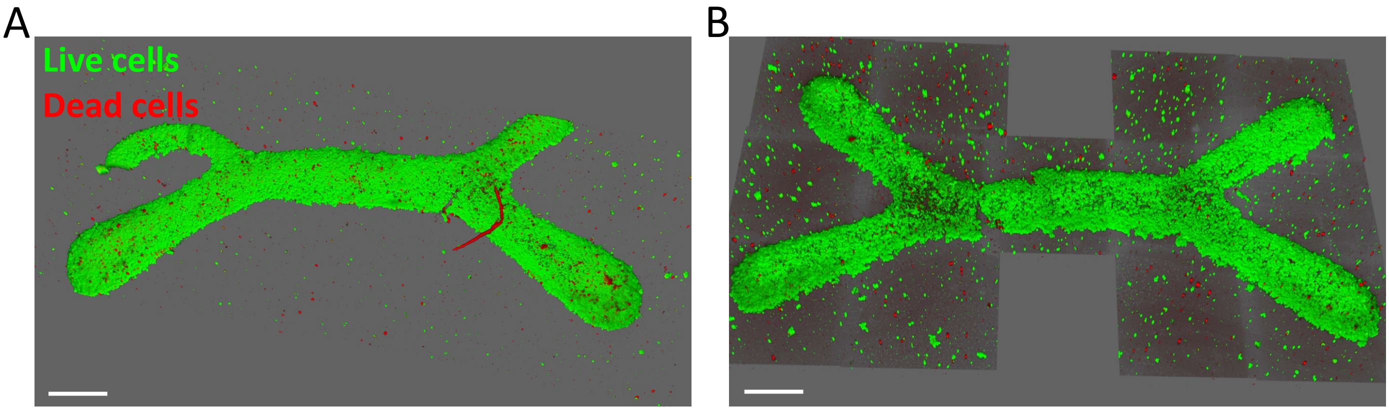


**Supplementary Figure 21 .** 3D reconstructions from confocal imaging of LIVE/DEAD assays performed on different tissues made from M-hiPSC 1, just after they were magnetically bioprinted (Live cells in green, dead cells in red, scale bar=500µm)

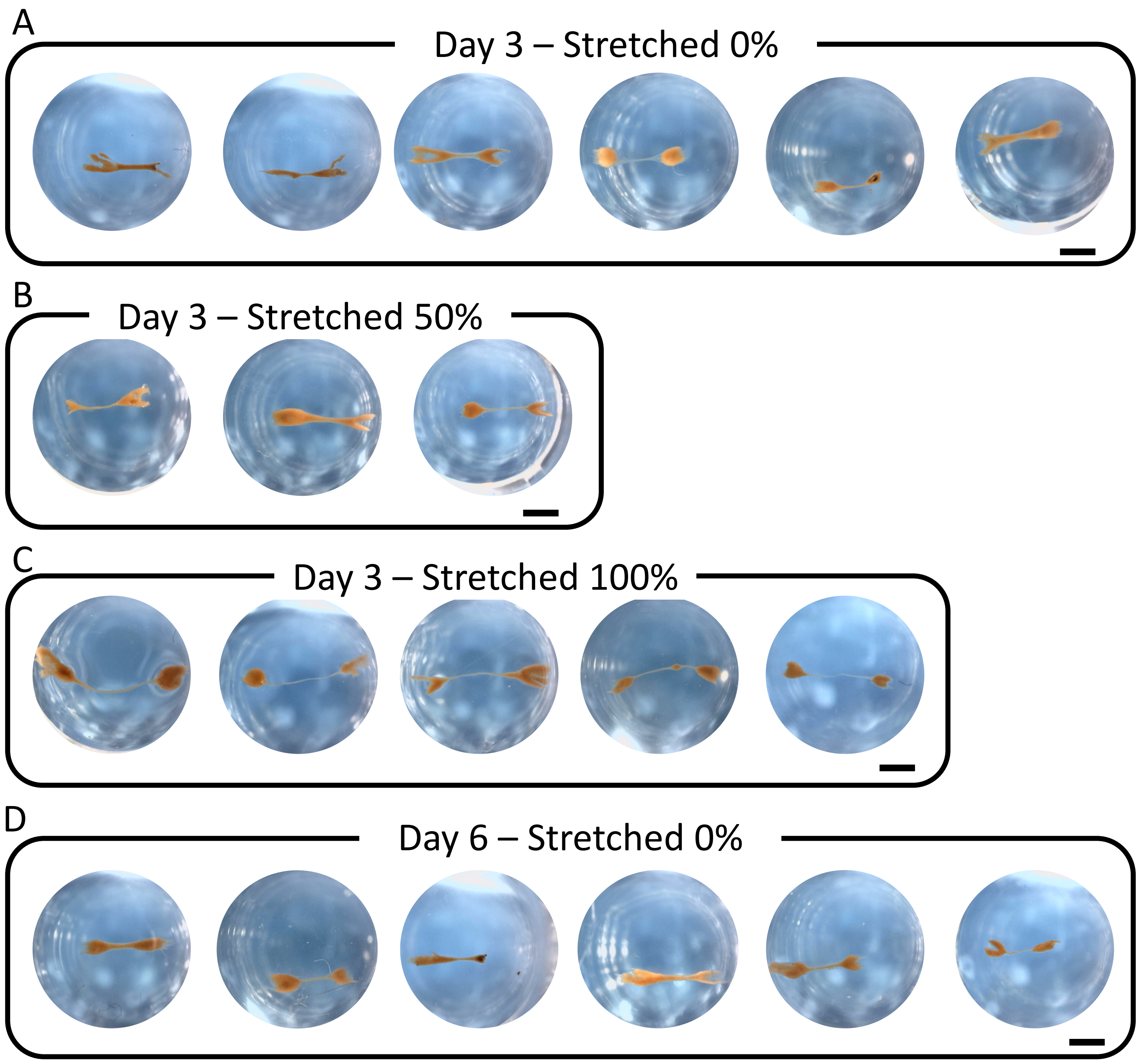


**Supplementary Figure 22.** Different tissues made of M-hiPSC 1 that were fixed and imaged in PBS in the wells of the 48-well plate in which they were preserved. Tissues that were fixed at day 3 that were stretched since day 1 by (A) 0%, (B) 50% or (C) 100% and at (D) day 6 without stretching. Tissues that were trapped between needles or stretched were fixed still trapped or stretched in the chip, then removed from the needles by scrapping the clamps with tweezers and placed in PBS filled wells. They were imaged from the top using a magnifier and a dino camera. (Scale bar=2mm)

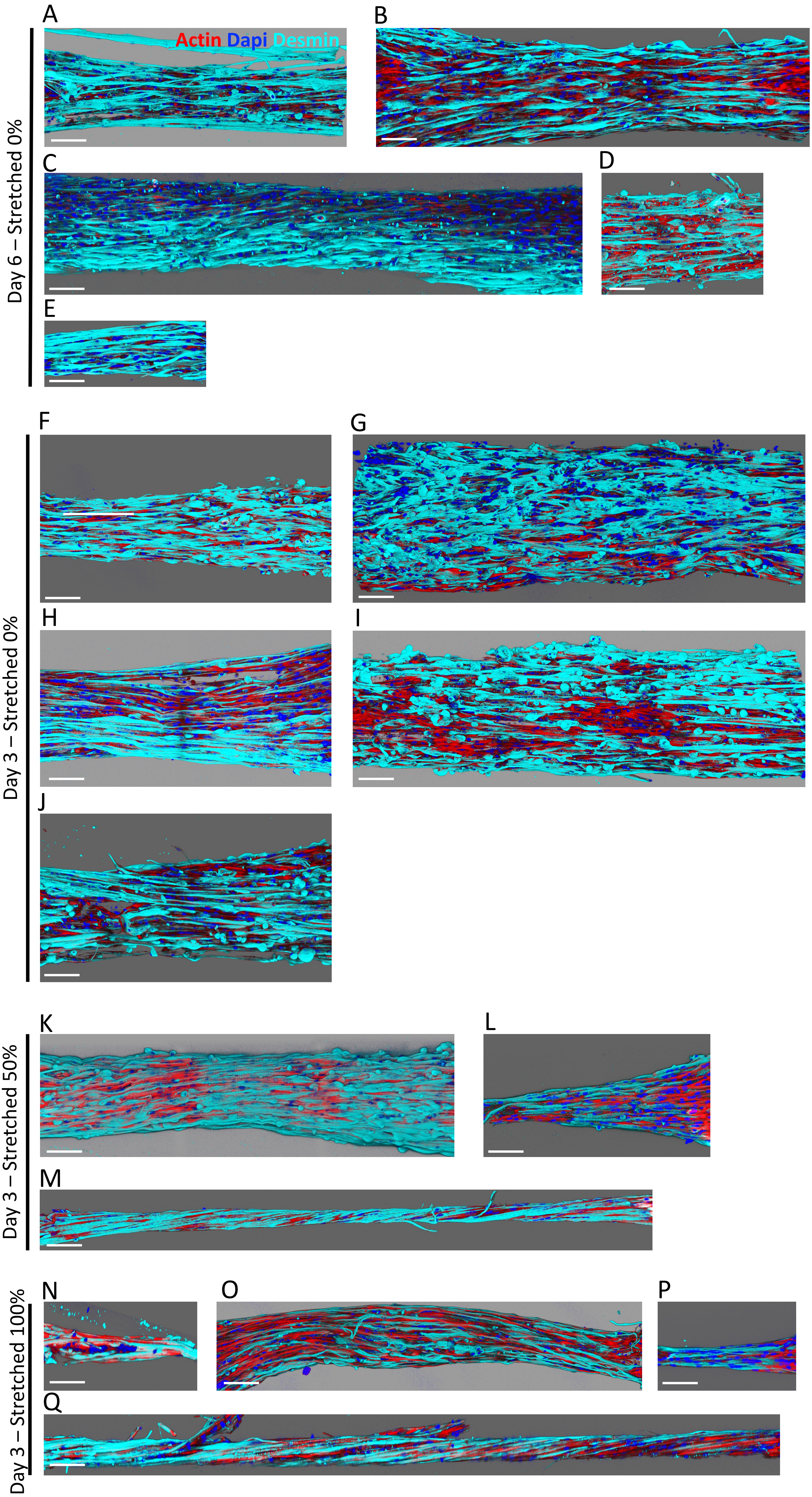


**Supplementary Figure 23 .** 3D reconstruction from confocal imaging of different desmin immunostained tissues made from M-hiPSC 1 cultured for (A-E) 6 days with 0% stretching and 3 days with (F-J) 0%, (K-M) 50% and (N-Q) 100% stretching (actin in red, nuclei in blue, desmin in cyan, scale bar=100µm)

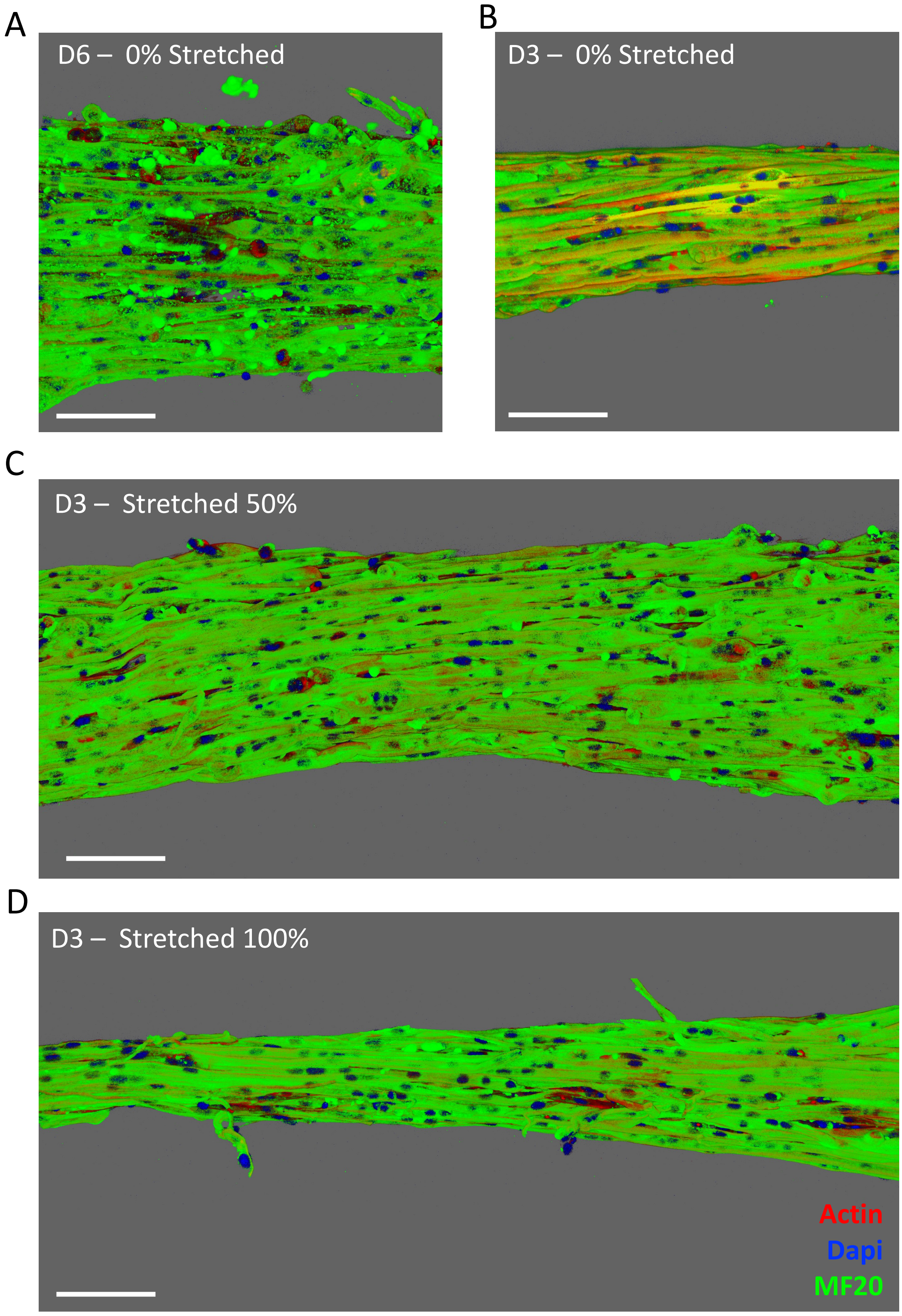


**Supplementary Figure 24 .** 3D reconstruction from confocal imaging of different MF20 immunostained tissues made from M-hiPSC 1 cultured for (A) 6 days with 0% stretching, and 3 days with (B) 0%, (C) 50% and (D) 100% stretching. They were immunostained with MF20 antibody to see myosin heavy chains. (actin in red, nuclei in blue, MF20 in green, scale bar in 100µm)

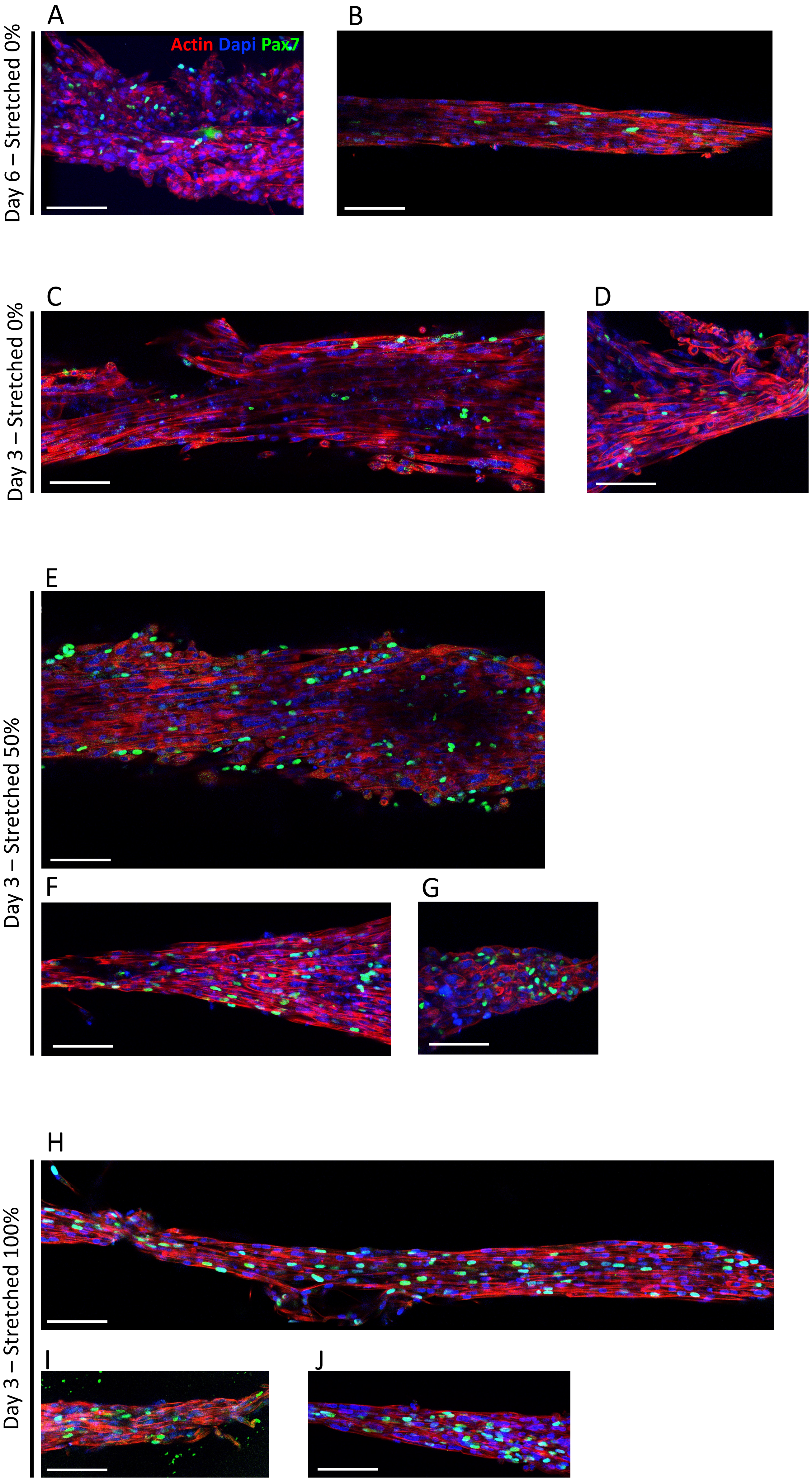


**Supplementary Figure 25 .** Confocal imaging of different Pax 7 immunostained tissues made from M-hiPSC 1 cultured for (A-B) 6 days with 0% stretching and 3 days with (C-D) 0%, (E-G) 50% and (H-J) 100% stretching. (actin in red, nuclei in blue, Pax7 in green, scale bar=100µm)

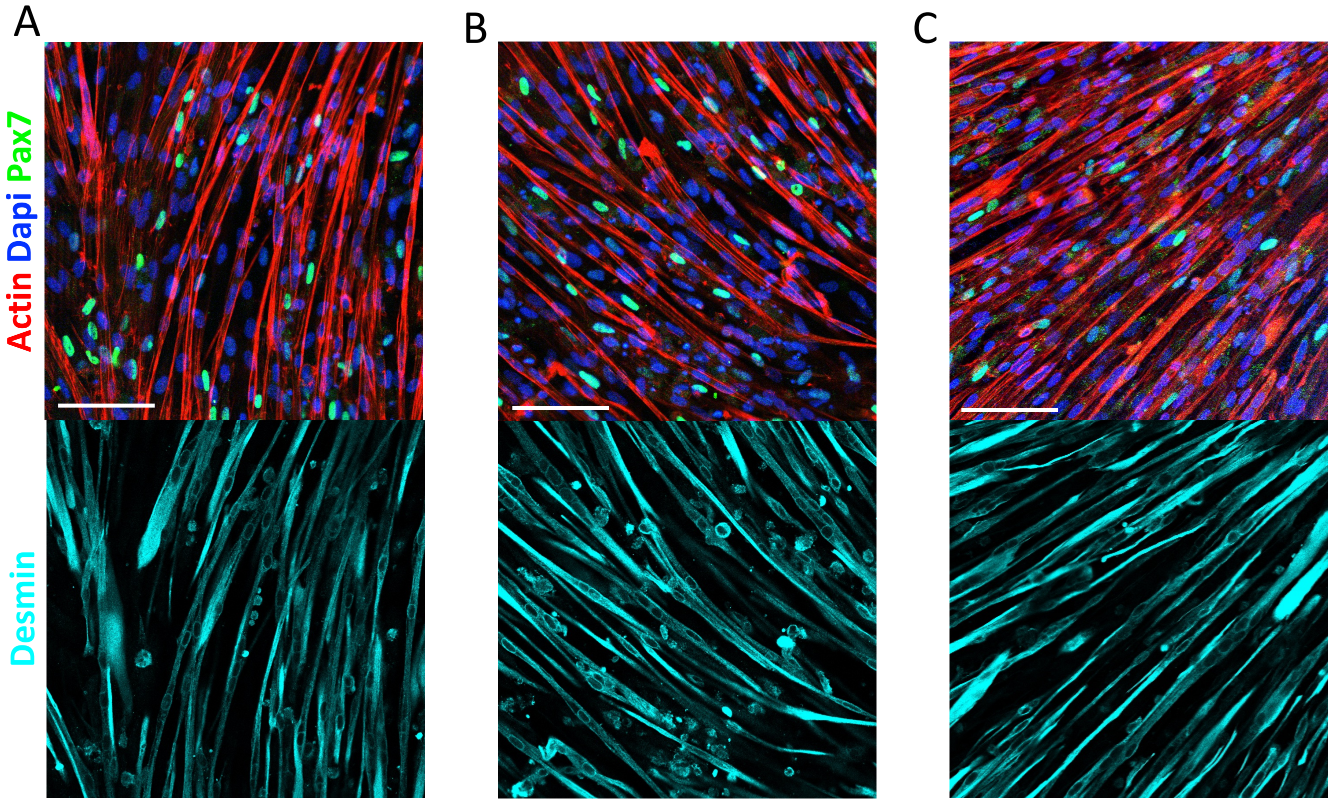


**Supplementary Figure 26 .** Confocal imaging of three different samples of M-hiPSC 1 cultured for 6 days of differentiation in 2D, immunostained for Pax 7 and desmin (Actin in red, nuclei in blue, Pax7 in green, desmin in cyan, scale bar = 100µm)

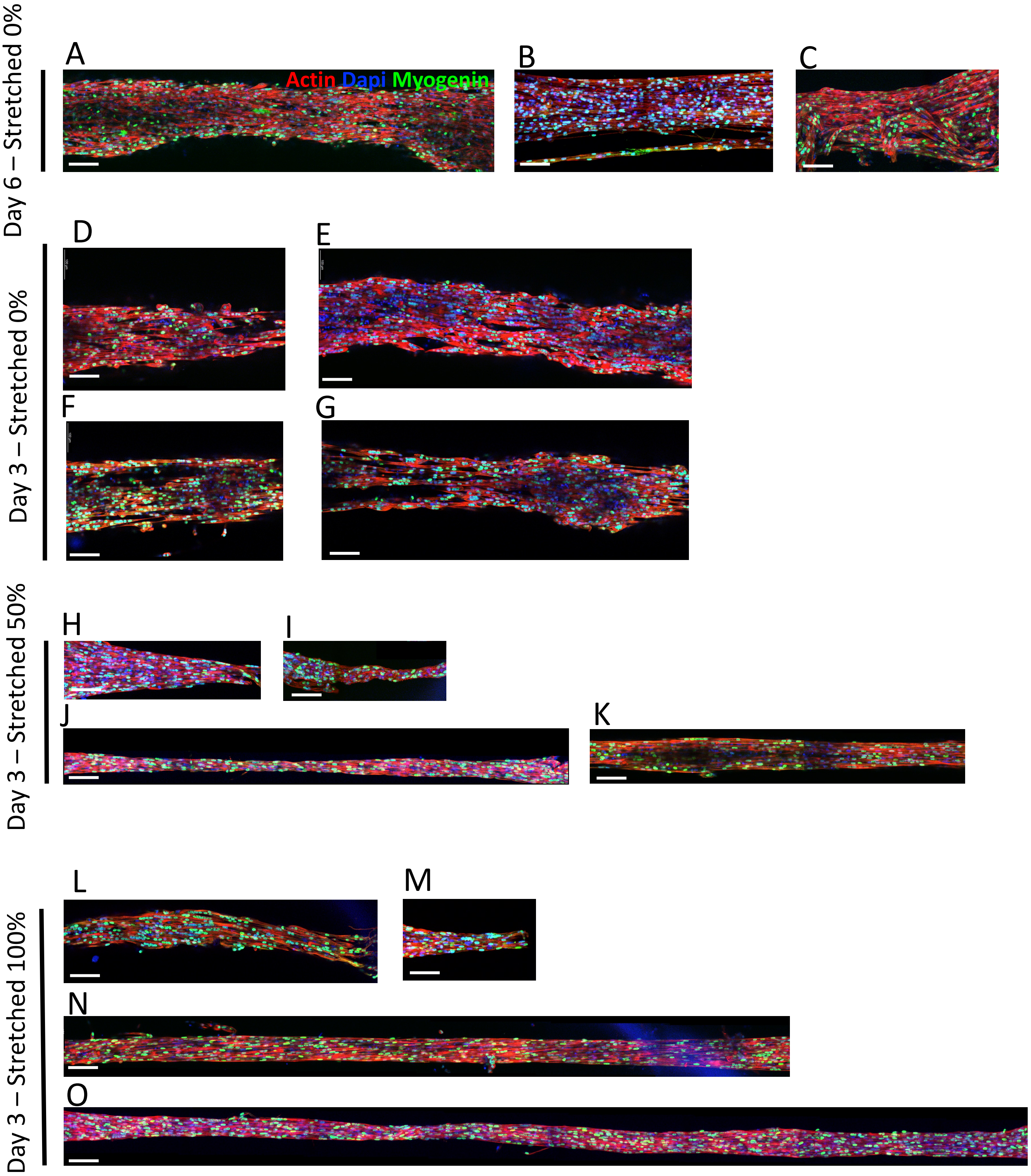


**Supplementary Figure 27 .** Confocal imaging of different myogenin immunostained tissues made from M-hiPSC 1 cultured for (A-C) 6 days with 0% stretching and 3 days with (D-G) 0%, (H-K) 50% and (L-O) 100% stretching. (actin in red, nuclei in blue, myogenin in green, scale bar=100µm)

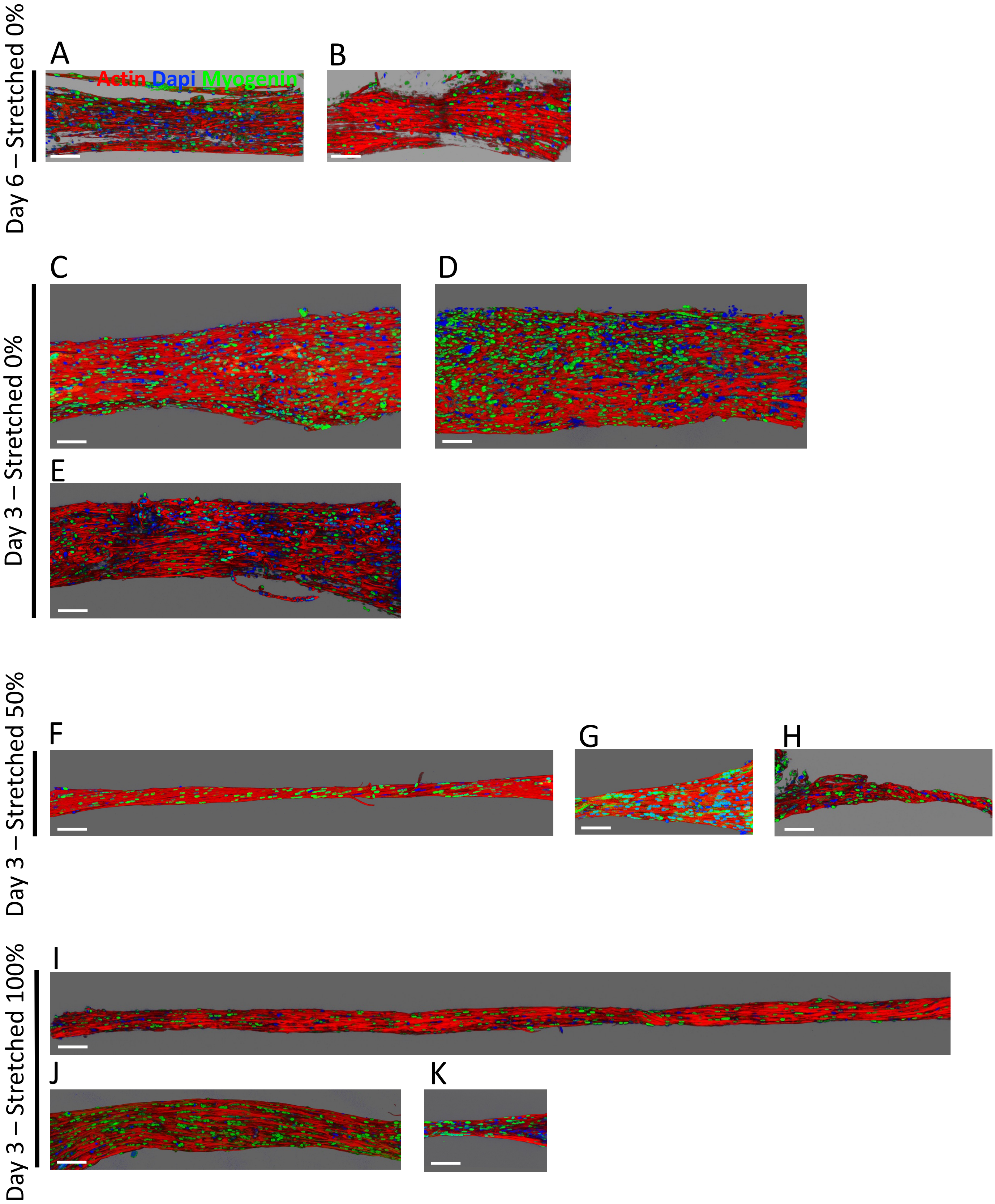


**Supplementary Figure 28 .** 3D reconstruction from confocal imaging of different myogenin immunostained tissues made from M-hiPSC 1 cultured for (A-B) 6 days with 0% stretching and 3 days with (C-E) 0%, (F-H) 50% and (I-K) 100% stretching. (actin in red, nuclei in blue, myogenin in green, scale bar=100µm)

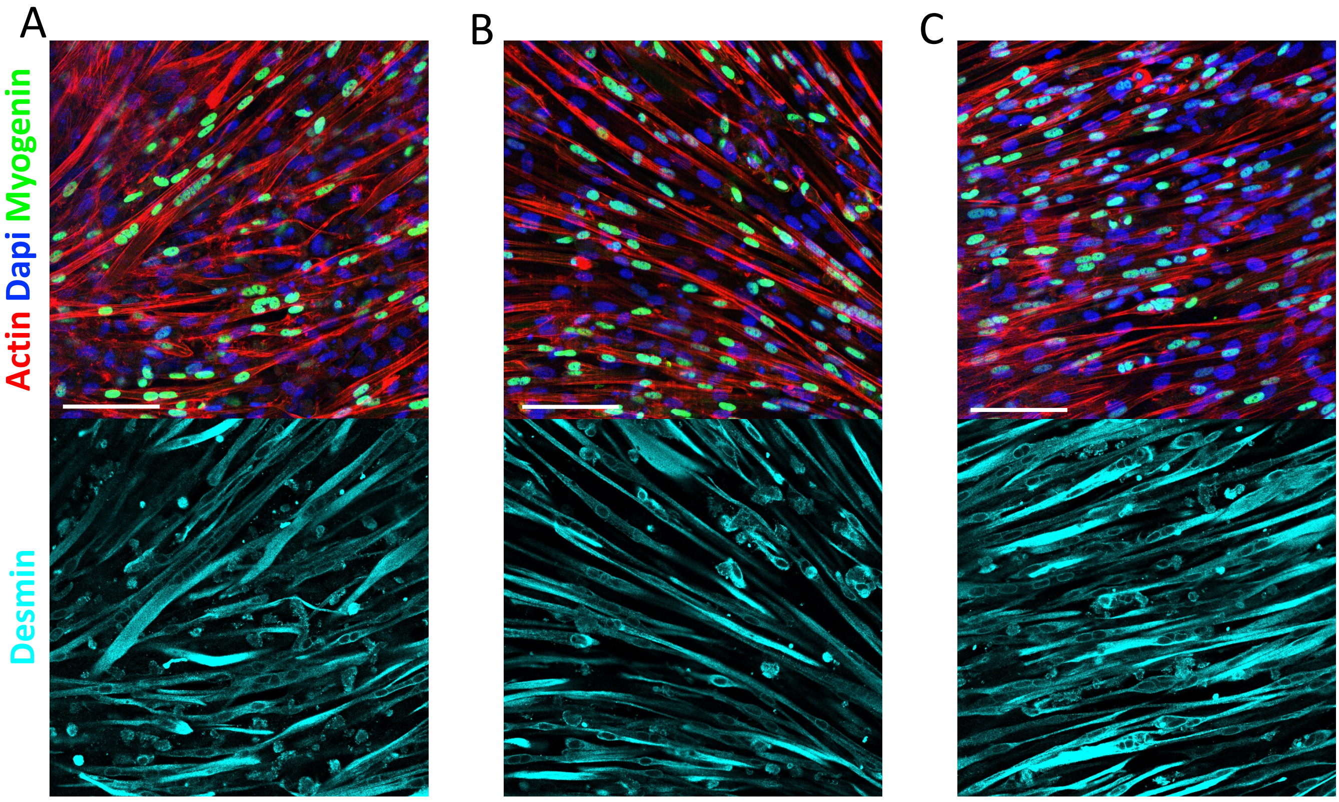


**Supplementary Figure 29.** Confocal imaging of three different samples of M-hiPSC 1 cultured for 6 days of differentiation in 2D, immunostained for myogenin and desmin (Actin in red, nuclei in blue, myogenin in green, desmin in cyan, scale bar = 100µm)

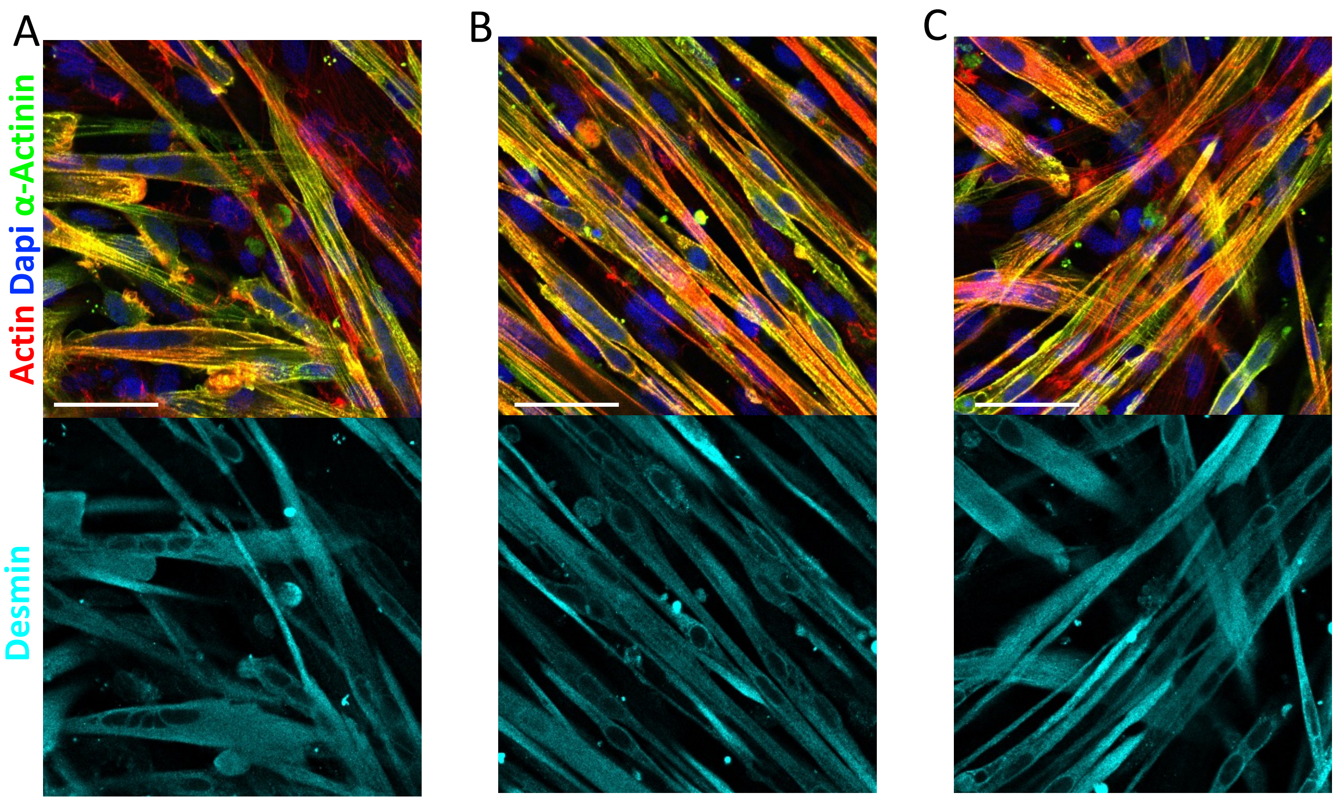


**Supplementary Figure 30 .** Confocal imaging of three different samples of M-hiPSC 1 cultured for 6 days of differentiation in 2D, immunostained for alpha-actinin and desmin (actin in red, nuclei in blue, alpha-actinin in green, desmin in cyan, scale bar = 50µm)

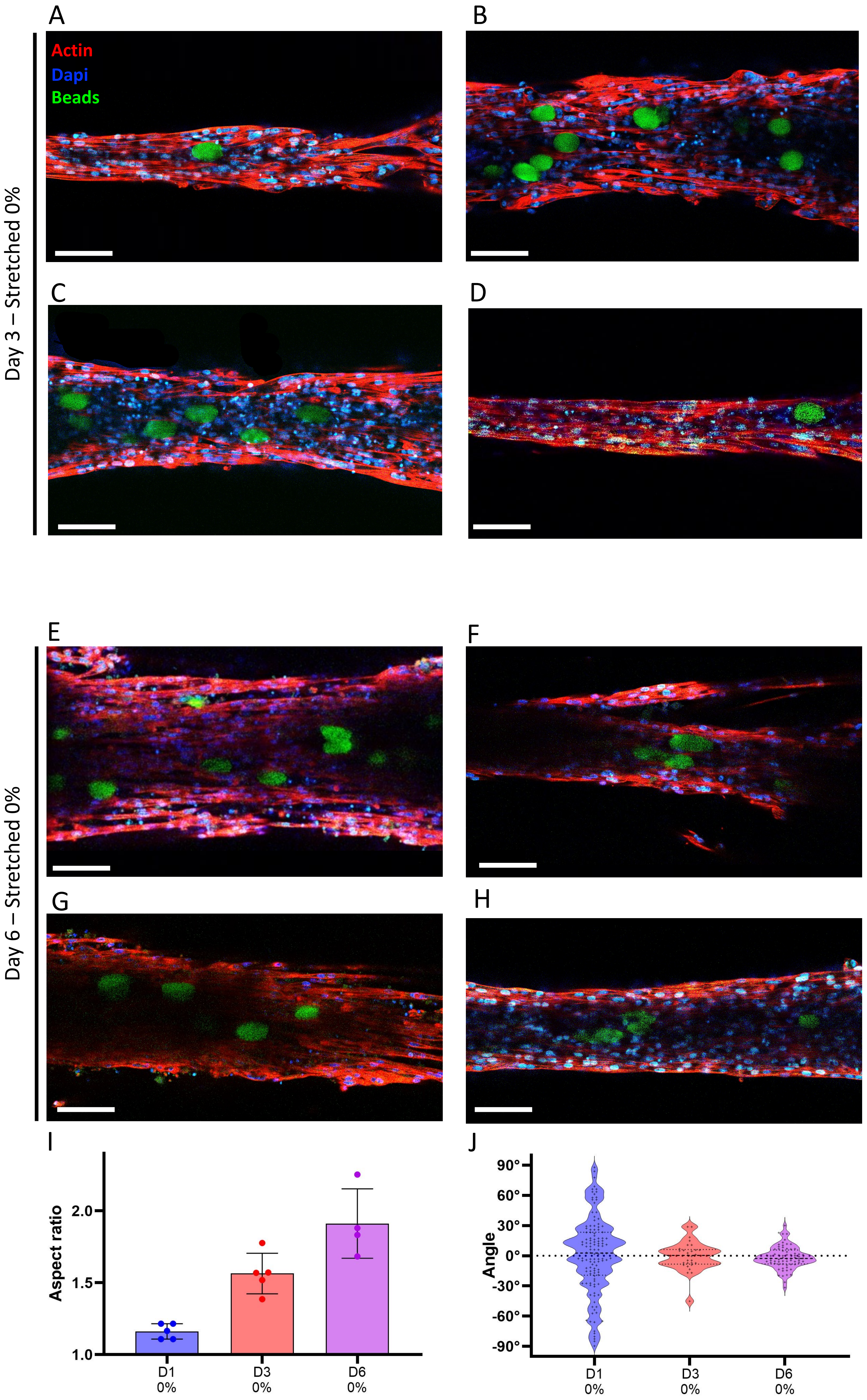


**Supplementary Figure 31.** Confocal imaging of different tissues made of M-hiPSC 1, in which polyacrylamide beads were embedded, which were cultured for (A-D) 3 days and (E-H) 6 days without stretching. (I) Aspect ratio and (J) orientation of the beads for tissues stretched by 0% on day 1, day 3 and day 6. Data points represent the average bead aspect ratio within one sample, and the angles of all beads across all samples. (Actin in red, Dapi in blue, beads in green, Scale bar = 100 µm)

A

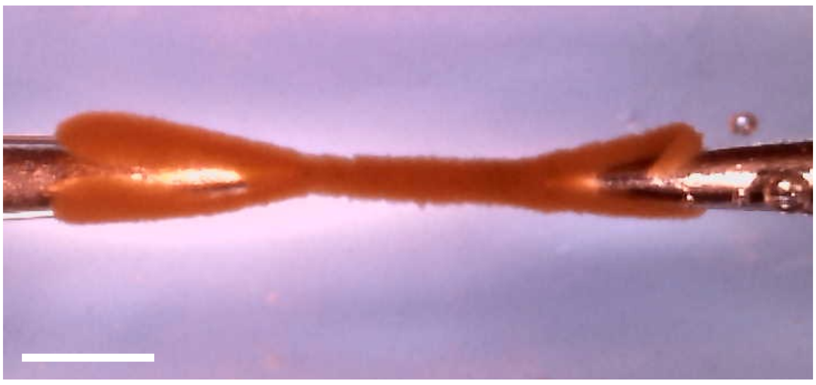

B

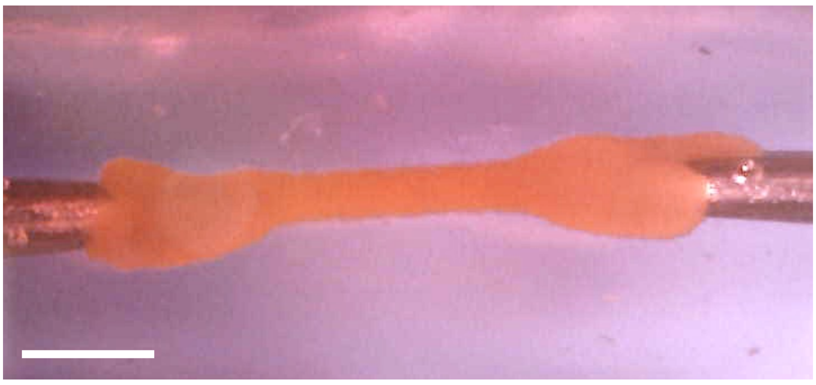

**Supplementary Figure 32.** Tissues magnetically bioprinted from magnetically labelled M-hiPSC 2 (A) and M-hiPSC 3 (B) trapped between needles. Scale bar=1mm

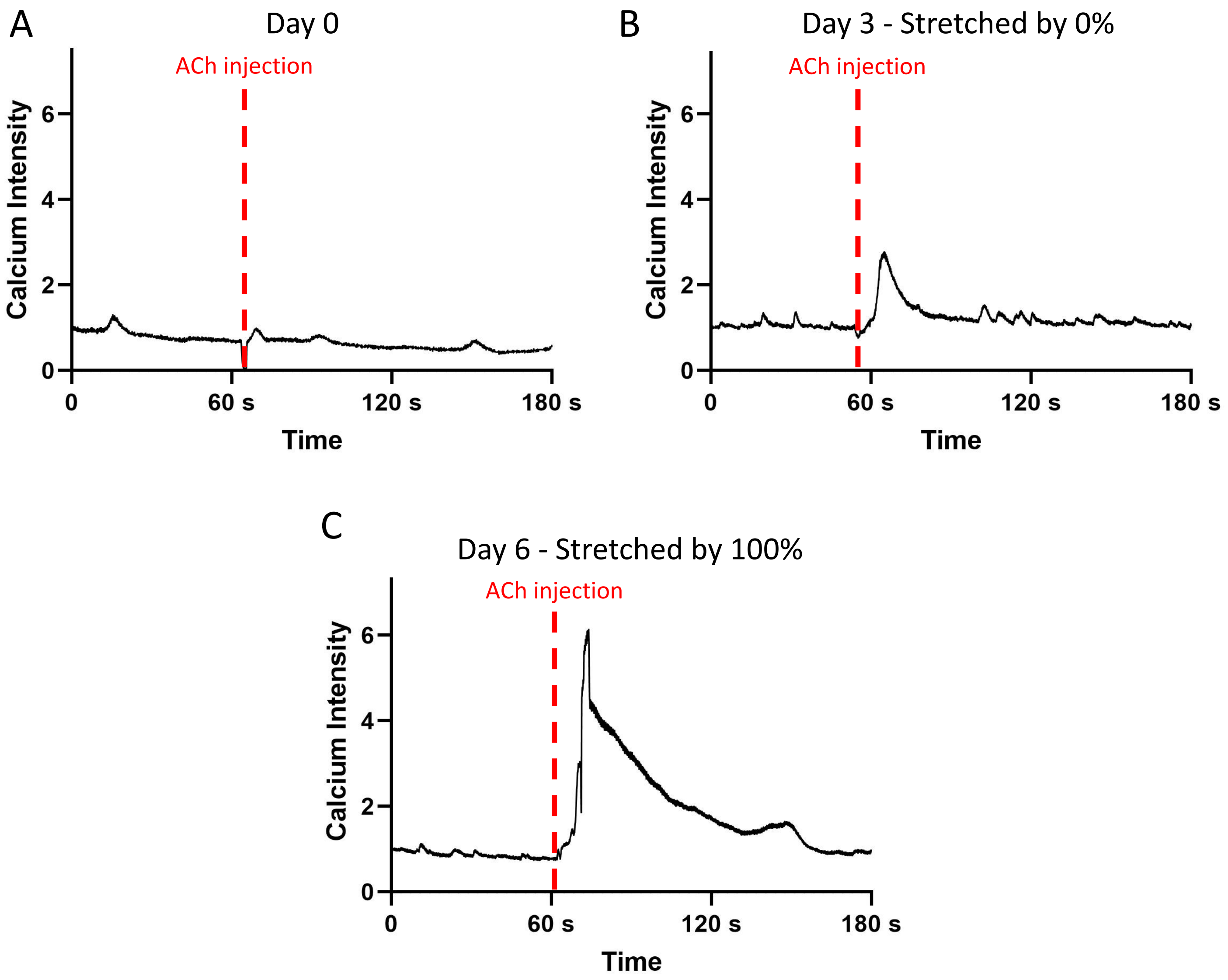


**Supplementary Figure 33.** Kymographs of the fluctuations of the calcium signal in a M-hiPS cell from (A) a M-hiPSC 2 tissue at day 0 just after bioprinting , (B) a non-stretched M-hiPSC 3 tissue at day 3, and (C) a M-hiPSC 2 tissue at day 3 stretched by 100% since day 1, treated with 1mM ACh after approximately 1min.

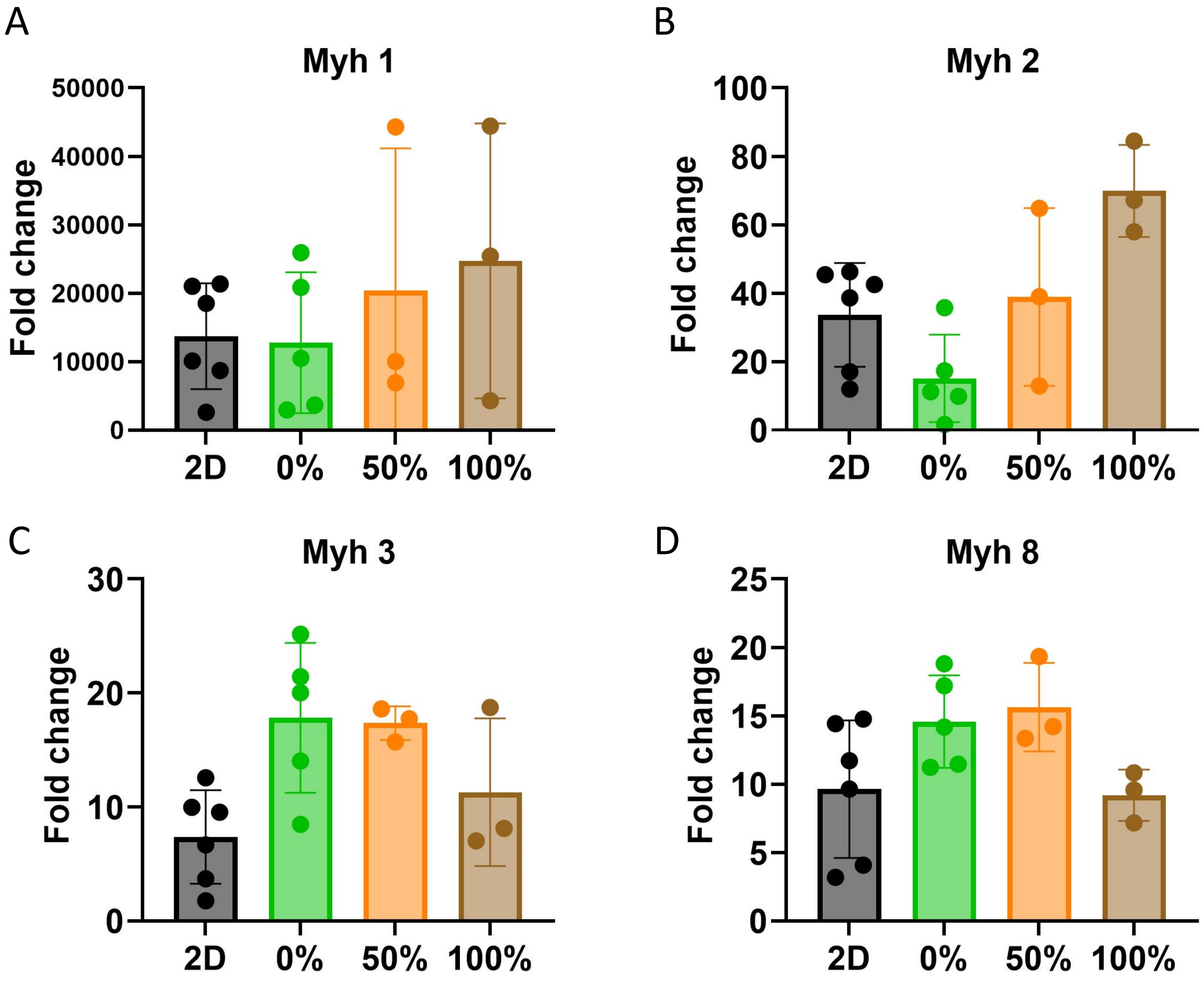


**Supplementary Figure 34.** Fold changes of the gene expression on day 3 of (A) Myh1, (B) Myh2, (C) Myh3 and (D) Myh8 for 3D tissues stretched by 0%, 50% and 100% and 2D cultures of M-hiPSC 1 , compared to the expression of cells collected at day 0.

| | M-hiPSC 1 | M-hiPSC 2 | M-hiPSC 3 |
|---|---|---|---|
| Sex of the patient | XX | XY | XX |
| Age of the patient | 67 | 22 | 48 |
| Origin of the sample | Fibroblasts | PBMC | PBMC 16525 |
| Cell supplier | Phenocell | | |
| Reprogramming method | Epi5 Kit - ThermoFischer #A15960 | | |

**Supplementary Table 1.** Information on the origin of the three patient derived hiPS cell lines from which the three M-hiPS cells were derived

| | |
|---|---|
| F_18S | GAGGATGAGGTGGAACGTGT |
| R_18S | TCTTCAGTCGCTCCAGGTCT |
| F_Myh1 | CTCTTTGTTGGGGCAACGG |
| R_Myh1 | TGAGTGCTCCTCAAGTTGGTC |
| F_Myh2 | TGAAGGAGAGGGAGCTGGT |
| R_Myh2 | ATGGGTACTCCTGAGGTTGGT |
| F_MYH3 | AGAGGAACTTTGACAAGGTGTTG |
| R_MYH3 | GGCTTCCTCGTAGGCATTTT |
| F_Myh8 | TGCTGAAGCAGATAGCAGCG |
| R_Myh8 | TACGAAGTGAGGGTGTGTGC |

**Supplementary Table 2.** Primers used for qRT-PCR.
